# The dinucleotide structure of NAD enables specific reduction on mineral surfaces

**DOI:** 10.1101/2024.10.11.617347

**Authors:** Delfina P. Henriques Pereira, Xiulan Xie, Tuğçe Beyazay, Nicole Paczia, Zainab Subrati, Jürgen Belz, Kerstin Volz, Harun Tüysüz, Martina Preiner

**Affiliations:** Microcosm Earth Center, Max-Planck-Institute for Terrestrial Microbiology, Hans-Meerwein-Str. 4; 35032 Marburg (Germany); Department of Chemistry, Philipps University Marburg, Hans-Meerwein-Str. 4; 35032 Marburg (Germany); Heterogeneous Catalysis, Max-Planck-Institut für Kohlenforschung, Kaiser-Wilhelm-Platz 1; 45470 Mülheim an der Ruhr (Germany); Metabolomics and small molecule mass spectrometry, Max-Planck-Institute for Terrestrial Microbiology, Karl-von-Frisch-Str. 10; 35043 Marburg (Germany); Department of Physics, Philipps University Marburg, Hans-Meerwein-Straße 6; 35032 Marburg (Germany); IMDEA Materials Institute, C/ Eric Kandel 2, 28906 – Getafe, Madrid (Spain)

**Keywords:** nucleotide cofactors, hydrogen, emergence of life, mineral catalysis, 2D NMR

## Abstract

Nucleotide-derived cofactors could function as a missing link between the informational and the metabolic part at life’s emergence. One well-known example is nicotinamide dinucleotide (NAD), one of the evolutionarily most conserved redox cofactors found in metabolism. Here, we propose that the role of these cofactors could even extend to missing links between geo- and biochemistry. We show NAD^+^ can be reduced under close-to nature conditions with nickel-iron-alloys found in water-rock-interaction settings rich in hydrogen (serpentinizing systems) and that nicotinamide mononucleotide (NMN), a precursor molecule to NAD, has different properties regarding reduction specificity and sensitivity than NAD. The additional adenosine monophosphate (AMP) “tail” of the dinucleotide, a shared trait between many organic cofactors, seems to play a crucial mechanistic role in preventing overreduction of the nicotinamide-bearing nucleotide. This specificity is also connected to the used transition metals. While the combination of nickel and iron promotes the reduction of NAD^+^ to 1,4-NADH most efficiently, in the case of NMN, the presence of nickel leads to the accumulation of overreduction products. Testing the reducing abilities of both NADH and NMNH under abiotic conditions showed that both molecules act as equally effective, soluble hydride donors in non-enzymatic, proto-metabolic stages of life’s emergence.

## Introduction

To form connections between undirected geochemical reactions and the first, directed, potentially even autocatalytic chemical reactions ultimately leading to metabolism, organic cofactors have dropped in and out of focus of research for decades.^[1–6,6–10]^ These cofactors are a common denominator between “information first”^[1,11]^ and “metabolism first” emergence of life hypotheses and thus might help to actively bridge this dated temporal separation of life’s most central components. The prebiotic synthesis of several organic cofactors was described or hypothesized via numerous synthetic ways.^[3,9,11]^ Based on theoretical work cofactors could have preceded enzymatic reactions playing central parts of self-sustaining non-enzymatic reaction networks .^[2,12]^

Many central cofactors are structurally related, mostly via an adenosine-derived “tail” added to a chemically active group (s. Fig.1). In biology, this tail mainly seems to have a recognition function for enzymes.^[13,14]^ The chemical properties and impact of this tail to the molecular dynamics of the molecule, however, remain largely underexplored, with only a few exceptions.^[5]^ Even when cofactors do not exhibit an obvious molecular connection such as adenosine, many (e.g. pterins or folates) derive from nucleotides in their biosynthesis.^[8,15–17]^

An important aspect of evaluating the role of common nucleotide structures in organic cofactors as the adenosine-derived “tail” is to investigate its function in a non-biochemical setting.^[5]^ So to take one of the most central organic cofactors, NAD, as an example: how much are the hydride transfers onto and from the nicotinamide influenced by the non-redox active adenosine part? Getting insight into the mechanisms in question is one of the main goals of this study, focussing on extant biomolecules rather than unknown prebiotic precursors.

NAD is a central redox cofactor in metabolism and is presumed to trace back to the Last Universal Common Ancestor (LUCA) of Archaea and Bacteria – and possibly beyond.^[2,18,19]^ In a previous study we demonstrated that NAD^+^ can react with H_2_ and metal powder (Ni, Co, Fe) to specifically form the biologically relevant form of reduced NAD (1,4-NADH) under hydrothermal conditions, however using the metal powders in great excess (333:1).^[20]^ Another study achieved NAD^+^ reduction without H_2_ – but also rather low yields – with iron sulfides.^[21]^ These publications introduced organic cofactors as possible transition points between geo- and biochemistry. Prior to these studies, only application-oriented heterogeneous catalysis pathways for the hydrogenation of NAD^+^ had been described.^[22]^

How can NAD be synthesized? Looking at biology, the de novo synthesis of NAD requires the synthesis of the deaminated form of NMN, to which AMP is transferred, followed by an amination to NAD^+^. All known de novo pathways are considered aerobic and are thought to have commonly emerged from an ancestral anaerobic pathway.^[23,24]^ In many organisms, but particularly in archaea and methanogens, an alternative salvage pathway directly adds ribose-5-phosphate to nicotinamide (Nam), producing NMN, avoiding two extra steps of deamidation and amidation.^[25]^

When it comes to NAD’s prebiotic synthesis pathways, several have been proposed e.g. via mineral-assisted synthesis under hydrothermal conditions^[9,26]^ or achieved in parts via nitrile- and amino acid derived pathways.^[11,27,28]^ This study starts from the assumption that nicotinamides were synthesized under prebiotic conditions and aims to evaluate whether the described adenine-derived tail could have played a role in a non-enzymatic context as well.

Approaching prebiotic redox reactions, one has to think about a possible primary electron source in the geochemical setting. Here, we are mainly investigating the options of water-rock-interaction systems, where protons of water are being reduced to hydrogen (H_2_) gas by electrons of Fe(II) containing minerals (serpentinizing systems).^[29,30]^ Minerals found in such systems are known to promote hydrogen activation and electron transfer for CO_2_ fixation.^[31–33]^ Apart from water-rock interactions constituting a rather simple, likely wide-spread geological scenario on a mostly unknown early Earth, H_2_ as an ancient electron source makes a lot of sense from a biological perspective as well. Looking at the reconstructed physiology of LUCA, it is likely that these first cells lived as anaerobic autotrophs, so were fixing carbon dioxide (CO_2_) with the electrons provided by H_2_[^19,34–36^] – a metabolic pathway still employed by many autotrophic prokaryotes today.^[37]^ In order to access the electrons of H_2_, these organisms possess enzymes called hydrogenases, which, depending on the exact organism, employ either Fe or Fe and Ni in their active centers.^[38]^

Incidentally, serpentinizing systems are rich in Fe and, depending on the system, can also be abundant in Ni as well. Hotter systems tend to feature Ni-Fe-alloys with higher nickel composition, while cooler ones are richer in iron.^[39]^ Nickel-containing intermetallic compounds such as awaruite (Ni_3_Fe) or taenite (NiFe_3_ to Ni_2_Fe) are products of the reaction of H_2_ with Ni(II) compounds in serpentinizing systems^[39–41]^ and are also found in meteorites.^[42]^ Native metals are also not impossible to find, if the conditions on site are reducing enough.^[43,44]^ In this study, we first tested various naturally occurring Ni- and Fe-containing compounds as “proto-hydrogenases” for NAD^+^ reduction and then compared NAD to NMN under hydrothermal conditions, to assess the relevancy of the adenosine-derived tail in pre-enzymatic reduction. We investigated the stability, reducibility and specificity of these organic cofactors and reduction products.

## Results and Discussion

### SCREENING NATURALLY OCCURRING IRON AND NICKEL CONTAINING MINERALS

Ni-Fe containing minerals found in hydrothermal settings were tested for the reduction of NAD^+^ under conditions comparable to those found in mild serpentinizing hydrothermal systems (40 °C, 0.133 M phosphate buffer pH 8.5, 5 bar H_2_, SI Scheme 1). These nanoparticular mineral powders were synthesized from scratch via the nano-casting method by using tea leaves as a template (s. SI methods) and were previously characterized.^[32,45]^ Additionally, we conducted pre- and post-reaction characterization with Scanning Transmission Electron Microscopy (STEM) of single representative reactions (SI Fig. 59–62). The metal content in these reactions is equivalent to that of the cofactor (1 metal atom per cofactor). The resulting H_2_ concentration at our conditions is 3.6 mM (using Henry’s law, s. SI Eq. 1 and references ^[18,29]^), which is comparable to the H_2_ concentrations found in the effluent of serpentinizing systems.^[46,47]^ The buffer was bubbled with N_2_ for 1 h and handled inside a glove box to approximate the anoxic conditions on early Earth. Several controls were implemented, including controls without metal and H_2_, respectively. The liquid phase was analysed by ^1^H-NMR (SI Tabs. 4–6, SI Figs. 24 and 25).

After 4 h, most of the dissolved NAD^+^ was still preserved in solution under all reaction conditions except the reaction with nanoparticular NiFe_3_ (nNiFe_3_), where NAD^+^ reacted faster (Fig. 2). Under H_2_, 1,4- and 1,6-NADH formed in all samples at different yields. Control experiments starting from 1,4-NADH were conducted, showing that 1,6-NADH is very likely a product of rearrangement from 1,4-NADH – metals can influence the amount of 1,6-NADH, but the patterns are not entirely conclusive (SI Scheme 7, SI Tabs. 17 and 18, SI Figs. 38 and 39). Samples under Ar also produced reduced NAD when a Fe-rich mineral was used (nNiFe_3_, nFe). In addition to transferring hydrides from H2 to NAD^+^, iron can oxidize, donating its own electrons either by producing nascent H_2_ gas, ultimately reducing NAD^+^ or by direct electron transfer to NAD^+^. This process can also be used as a proxy for the constant H_2_-production in serpentinizing systems.^[20]^ Ni by itself is H_2_-dependent in the promotion of NAD^+^ reduction.

STEM imaging before and after the reactions (the latter including a washing and dilution step to assure true surface alteration) confirms that Fe, both under Ar and H_2_, gets associated with phosphate ions in ratio that suggests the formation of iron phosphate, while Ni does not associate with phosphate, suggesting it stays in its native form (SI Fig. 61 and 62).^[20]^

The Fe in Ni-rich minerals is expected to slowly reduce NAD under Ar conditions. However, it is likely that the resulting products do not reach the detection threshold within 4 h. Overall, bimetallic minerals are significantly more efficient than the individual transition metals when hydrogen is available. Introducing one Ni atom to a Fe atom increases the yield by 300% (nNiFe vs. nFe). Their properties, already observed in a previous study,^[20]^ seem to complement each other for the reduction of NAD^+^ with H_2_: Fe being mostly an electron donor, while Ni promotes hydride transfer from H_2_. These complementary roles have also been described in other publications, suggesting charge transfers from Fe (more electropositive) to Ni could increase the electron density in Ni.^[32,48]^

### THE “UNIVERSAL” ADENINE NUCLEOTIDE IN ORGANIC COFACTORS

Many central cofactors share an adenosine-derived tail (Fig. 1) attached to a “functional” part. In the case of NAD, we define NMN as the functional part (containing the hydride-transferring nicotinamide) and adenosine monophosphate (AMP) as the adenosine-derived tail. NAD is stable in water, with its pH range depending on the reduction state of the nicotinamide: NADH is more stable at pH>7, while NAD^+^ is more stable under acidic conditions.^[20]^ To investigate the role of the AMP-tail in a prebiotic context, several experiments were designed to compare NAD and NMN. 2D ^1^H-NMR enabled the assignment of peaks for 1,4-NMNH and NMN in accordance with literature and in comparison to the pair NAD/NADH (SI Tables 2 and 3, and SI Fig. 1–19).^[49]^ We initially focussed on nNiFe_3_, the most efficient of the Ni-Fe minerals in the above described NAD experiment (s. Fig. 2). All other reaction conditions (buffer, pH, temperature, metal to cofactor ratio) were maintained (SI Schemes 3 and 4). Reduction products were characterized through different 2D-NMR correlation spectra (SI Scheme 9 and 10; SI Figs. 42–66) and Liquid Chromatography Mass Spectroscopy (LC-MS, SI Fig. 56–58. Products were quantified via 600 MHz ^1^H-NMR spectroscopy using DSS as an internal standard (SI Table 11). Also controls with Ar instead of H_2_ and without metal were conducted (SI Fig. 29, SI Tab. 9).

**Figure 1.**
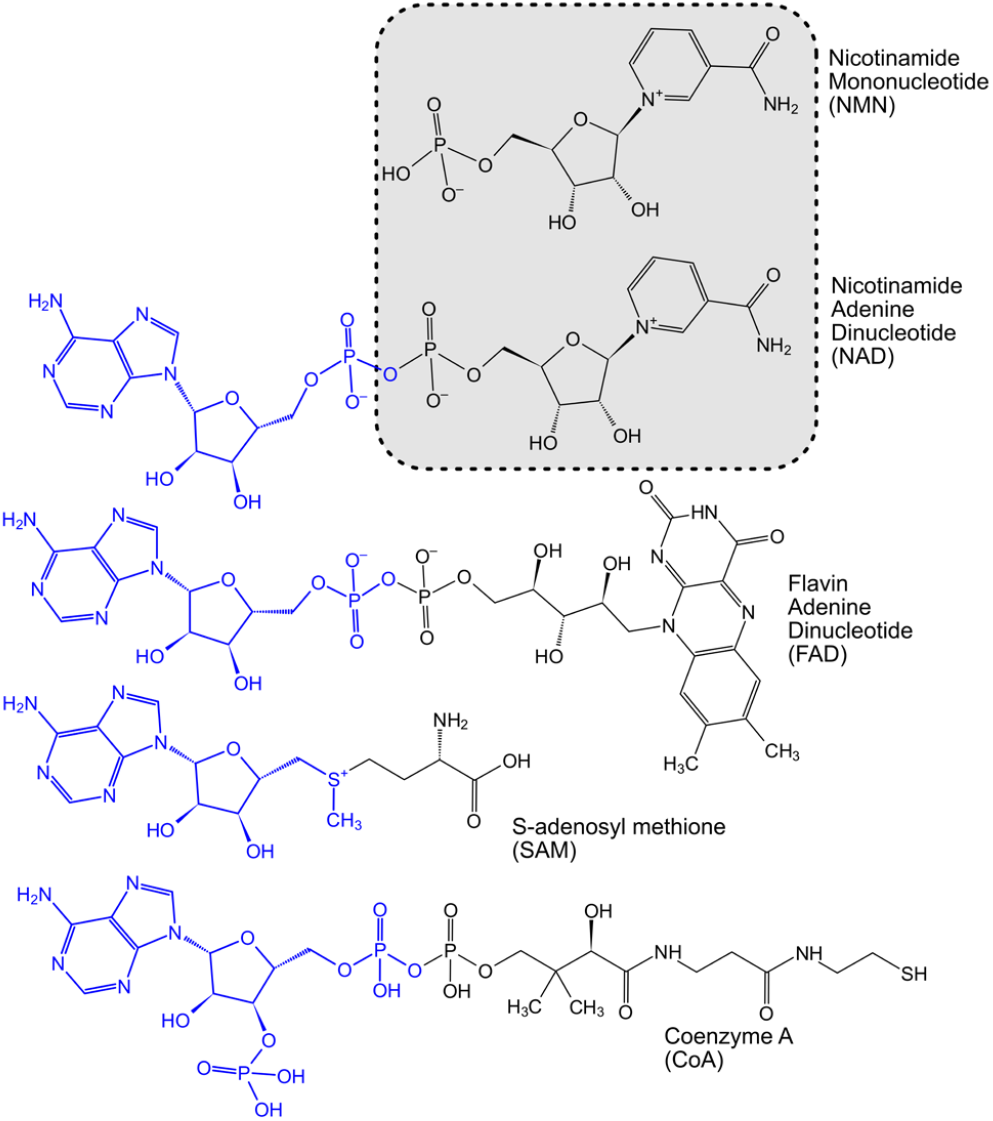
A selection of central organic cofactors with an adenosine- based tail (blue) connected to a functional part that determines the role of these cofactors in metabolism (black): Electron/hydride transfer, methyl transfer or acetyl-transfer. In grey, the function-associated half of nicotinamide dinucleotide (NAD) is highlighted: nicotinamide mononucleotide (NMN).

**Figure 2.**
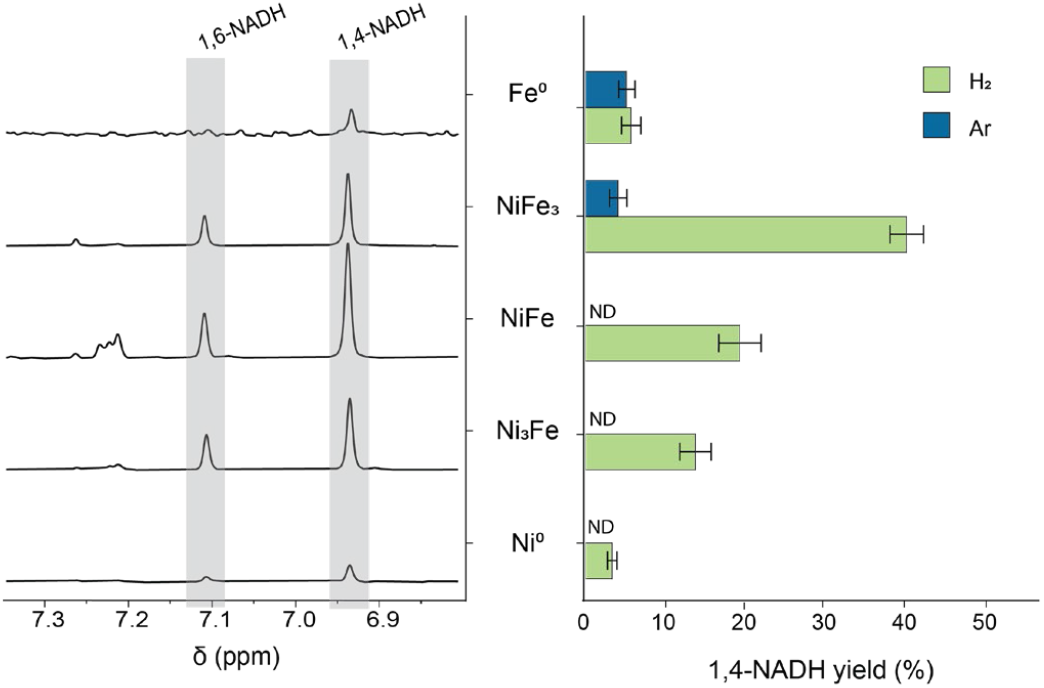
NAD^+^ reduction at 40 °C over 4 h with equimolar amounts (normalized to 1 metal atom per NAD molecule) of several Ni-Fe-alloys and 5 bar of H_2_ (or Ar). A) Segment of the ^1^H-NMR spectra where the chemical shift of the hydrogen on the 2^nd^ nicotinamide carbon is visible upon reduction. 1,4-NADH features a characteristic peak at δ = 6.9 ppm and 1,6-NADH at δ = 7.1 ppm (SI Table 4). B) Yield of 1,4-NADH obtained for several metals after 4 h under H_2_ and Ar. Reduction under Ar is detected only with minerals whose metal content is ≥ 75% Fe. With 5 bar of H_2_, all metals can facilitate 1,4-NADH synthesis. Mixed alloys are more efficient than the pure metals.

Without metals, NMN does not react and remained stable (SI Fig. 28 and 30). Under Ar, NMN still got reduced due to the abundant iron in the mineral compound, but more slowly than under H_2_ (SI Tables 10 and 11). An additional 2 h experiment with NAD^+^ under H_2_ solidified the hypothesis that the increase of 1,4- and 1,6-NADH is steady and inversely proportional to the decrease of NAD^+^ in solution (Fig. 3C, SI Scheme 4). After 4 h, on average 57% of NAD^+^ was reduced with 26% remaining oxidized. The remaining 17% can in part be attributed to nicotinamide formation but also unassigned degradation reactions and loss via surface absorption (SI Table 12 and SI Fig. 32).^[50]^

**Figure 3.**
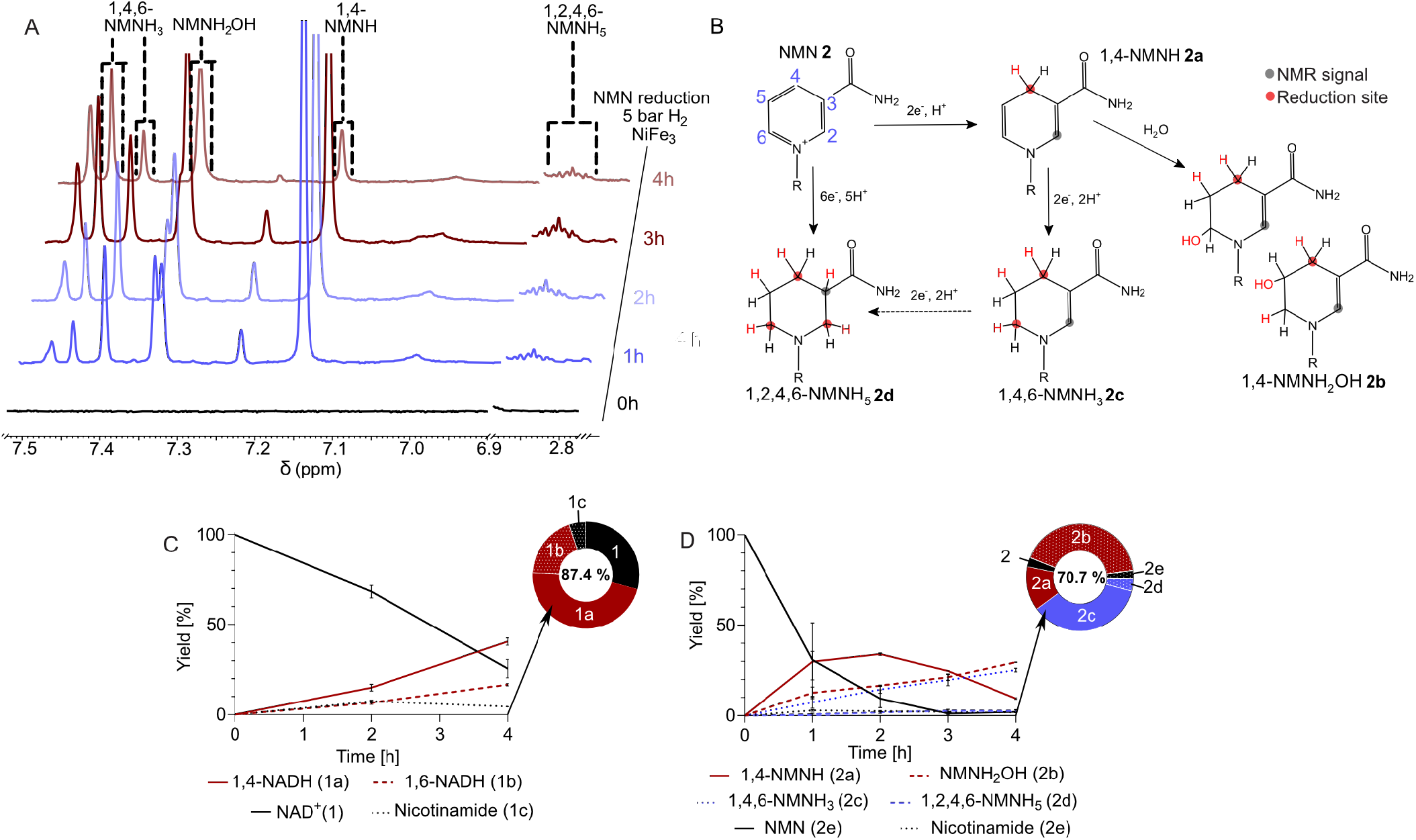
The reduction of NMN and NAD^+^ promoted with equimolar quantities of NiFe_3_ nanopowder (normalized to the number of metal atoms) and 5 bar of H_2_ (40 °C) was monitored for 4 h (SI Schemes 3 and 4). Both reactions were carried out in oxygen-free, aqueous solutions at a pH of 8.5 (0.133 M phosphate buffer). A) ^1^H-NMR (600MHz) spectra monitoring NMN reduction over time. Dashed lines mark the peaks used for identifying and quantifying reduced NMN species previously identified via 2D NMR. Due to the complexity of the mixture not all peaks could be assigned (SI Scheme 9 and 10, SI Figs. 42–56) B) Proposed mechanism for NMN reduction with H_2_ and Ni-Fe minerals. The reduction site and additional protons are highlighted in red. Full arrows represent proposed reaction mechanisms supported by the data obtained, while the dashed arrow is an additional reaction that could not be entirely excluded. Grey circles indicate the proton providing the NMR signal for quantification. In bold font numeral assignments for NMN reduction products are made. C) Time course of NAD^+^ reduction. Reduced NAD species are plotted as relative percentage to a metal-free control sample (SI Methods) – all time points represent the mean and SD of at least triplicates of the same reaction. The ring chart represents the distribution of products after 4 h, percentage in the center indicates the entirety of assigned products. D) Time course of NMN reduction. Reduced NMN species are plotted as relative percentage to a metal-free control sample – all time points represent the mean and SD of at least triplicates of the same reaction. The ring chart represents the distribution of products after 4 h, percentage in the center indicates the entirety of assigned products.

NMN, however, shows a completely different reaction profile. After only 1 h, 69% of the starting NMN had been consumed, and a variety of products was observed in 1D ^1^H-NMR (SI Scheme 5, SI Table 13 and SI Fig. 33). 2D-NMR spectroscopy facilitated the identification of the overreduction of NMN’s nicotinamide ring with two and three hydrogenation sites, so 1,4,6-NMNH_3_ (**2c**), and 1,2,4,6-NMNH_5_ (**2d**) respectively (Fig. 3A, B and D; SI Fig. 38–51).

While the fully reduced species **2d** formed quickly and its concentration remained constant over time, the concentration of twice reduced **2c** increased with once reduced 1,4-NMNH decreasing. This indicates that not all reductions are a step-wise process (s. Fig. 3B), especially in the case of **2d**.

Under Ar, **2d** did not form at all, demonstrating that H_2_ is necessary for the full hydrogenation of the nicotinamide ring (SI Tab. 9, SI Figs. 28 and 29). **2c**, however, was also forming under Ar, albeit in far lower yields (3%) than under H_2_ (25%) after 4h). The yield of 1,4 NMNH was relatively similar in both atmospheres (7% under Ar; 10% under H_2_). Transferring these observations to environmental conditions suggests that less reducing conditions could be favourable for specific NMN reduction.

The above mentioned products accumulated in different amounts, 1,4-NMNH being the main product after 1 h under H_2_ (Fig. 3A and D). Other side products formed at a comparable rate, rapidly depleting the reagent NMN. Consequently, the production of 1,4-NMNH seems to have stopped after 2 h and subsequently began to decrease in concentration. The concentration of **2c** continuously increased over time. Even though the concentration of 1,4-NMNH decreased from 35% to 9% in 2 h, the total amount of reduced NMN remained relatively stable, exceeding 60%. This suggests that 1,4-NMNH is the first and main product of NMN reduction, which can subsequently undergo further reduction to other species, mainly **2c**.

We were able to exclude two products commonly found in NAD^+^ reduction, where the 2^nd^ or 6^th^ carbon of the nicotinamide ring is reduced.^[22,51,52]^ Subsequently, reduction products presumably starting with these two one-time reduced products could be excluded (SI Scheme 9,SI Figs. 44 and 45). In the case of NADH, its 1,2-reduced form is known to be unstable, so it is likely this is the case with 1,2-NMNH as well, leading to its absence in our reaction (SI Scheme 10).^[53]^

In the case of 1,4-NMNH loss over time, we saw several routes: i) mainly the further reduction to **2c**, ii) 1,4-NMNH getting hydrolysed at it 5^th^ or 6^th^ position, and iii) 1,4-NMNH engaging in a Diels-Alder type reaction with 1,6-NMNH (SI Scheme 11). Via LC-MS we were able to exclude Diels-Alder products and confirm it to be a hydration product, most likely OH^-^ being added to 1,4-NMNH at the 6^th^ position (Fig. 4B, SI Fig. 58) Ultimately we could assign the hydration product NMNH_2_OH (**2b**) to be the peak at 7.35 ppm in the ^1^H-NMR (SI Figs. 50 and 51). Both **2b** and **2c** have several conformational isomers, which result in several peaks within the ^1^H-NMR spectrum as indicated in Fig. 4A.

**Figure 4.**
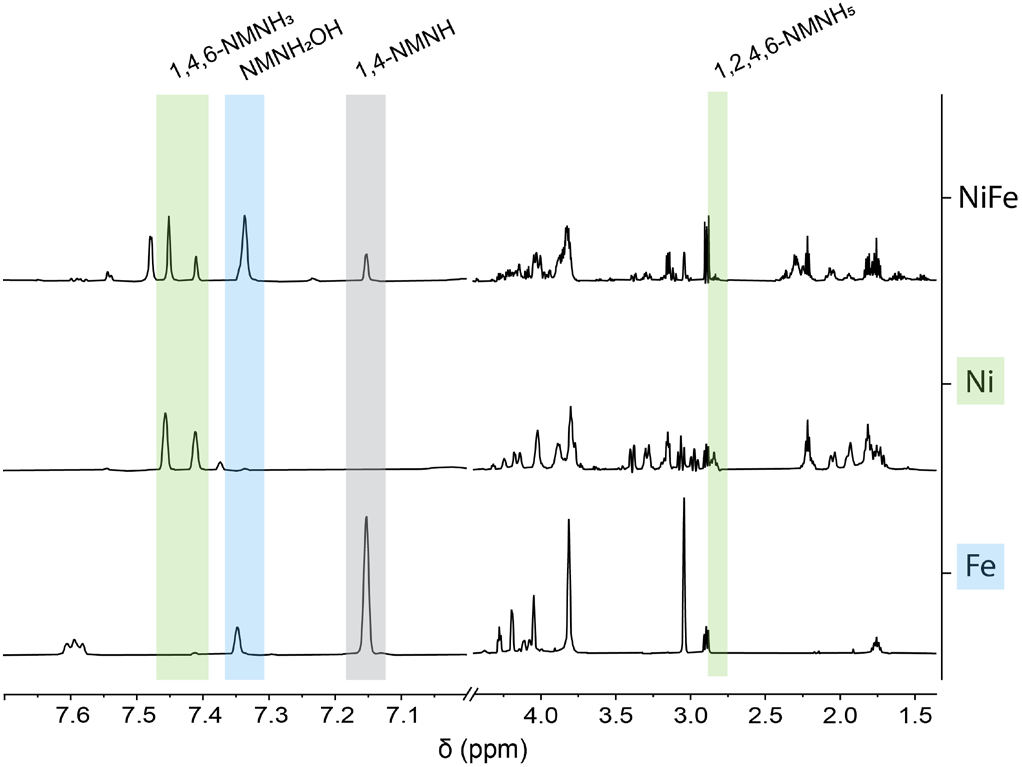
Selectivity and efficiency of NMN reduction with H_2_ gas is influenced by the type of metal prevalent. The reduction of NMN assisted by nNiFe alloys (cofactor ratio 1:1) yields products that directly correlate with the products obtained when utilizing the metals separately in micropowder form (metal-cofactor ratio 200:1). μNi^0^ promotes the accumulation of 1,4,6- NMNH_3_ and 1,2,4,6-NMNH_5_ (green) and other side products. 1,4-NMNH (white) seems to not react further when using μFe^0^, accumulating over time.

An often reported side-product of NAD reduction (e.g. via cyclic voltammetry) is a 4,4’-linked NAD dimer,^[54]^ which also qualifies as a possible side reaction of NMN reduction, too. Here, after careful interpretation of our 2D NMRs of the 1 h and 4 h reaction with NMN and the 4h reaction with NAD, we can exclude the presence of such dimers (s. SI Fig. 55; no peak at 40 to 50 ppm in ^13^C of a bridging methine corresponding to the linkage). This was also confirmed via LC-MS (no double charged molecules were detected). As these dimers are a direct result of radical-forming 1e^-^ transfers onto NAD,^[55]^ we can draw the conclusion that direct hydride or 2e^-^ transfer is the present mechanism in our reactions.

After quantification of all identified species, we can account for at least 70.7% of transformed NMN for all reactions, often more. Unidentified species encountered in lower yields can also stem from the differently reduced versions of the degradation product nicotinamide.^[56]^ It is furthermore possible, as mentioned above, that some NMN was lost due to interaction with the mineral surface.

In summary, we demonstrated that NMN is much more reactive than NAD^+^. In 1 h, more NMN is consumed than NAD^+^ in 4 h, under the same experimental conditions. The first and main product of both reactions seems to be 1,4-NADH/NMNH. However, while 1,4-NADH remains stable in solution, 1,4-NMNH quickly undergoes further reduction to form increasingly reduced products. Even when the experimental conditions are more reducing or using a higher metal to cofactor ratio, 1,4-NADH does not get overreduced and is stable.^[20]^

Where does this specificity for 1,4-NADH come from? It is known and well-described that NAD(H) in aqueous solution alternates between a folded and open conformation.^[57–61]^ This could shield the nicotinamide ring from excessive overreduction – and possibly also from side reactions such as hydrolysis and Diels-Alder reactions described earlier (SI Scheme 11).

We deduce from these results that – in a mineral-dependent prebiotic context – the adenosine-derived tail can be essential for the targeted reduction of the 1,4-position, as well as for the stability of this specific reduction product. Nevertheless, we wanted to put to the test how much the choice of catalyst influences the outcome of the reactions.

### TO EACH ITS OWN METAL AND MECHANISM

Starting from the observation that during NMN reduction, its biologically active form 1,4-NMNH is a main product decreasing with the length of the reaction, we deducted that less efficient catalysts (e.g. less surface area) might help to reduce NMN more specifically than the previously used rather efficient nanoparticular NiFe-alloys (nNiFe, nFe…).

As both nNi and nFe visibly worked less efficiently for NAD^+^ reduction but Fe promoted reduction in higher yields, we decided to work with the same nanoparticular iron powder used for NAD^+^ reduction (Fig. 2 and SI Scheme 2). While reducing NMN steadily, the nFe powder by itself (1:1 ratio to the cofactor) did not promote the formation of three times reduced **2d**, while twice reduced **2c** is only produced in almost untraceable amounts (s. SI Table 8 and SI Fig. 26). The hydration product **2b** visible as a peak at 7.35 ppm still accumulated over the 4 h reaction time. Other possible reduction products shown in SI Scheme 9 were not detected.

In order to explore whether this observation is a mere metal-dependent effect, we repeated experiments with NMN and H_2_ over commercially available Fe and Ni micropowder (μFe, particle size: <150 μm; μNi, particle size: 3–7 μm). The metal-cofactor ratio was 200:1 to guarantee the detection of even low concentration side products. The results (Fig. 4) show a remarkable trend of overreduction with μNi, while μFe mostly displays two main products: 1,4-NMNH and NMNH_2_OH.

Comparing the spectra of NMN reduction with μNi only and μFe only with those of nNiFe (Fig. 4, SI Fig. 37), it becomes apparent, that the distinct product patterns of each metal prevail within the mixture. Note that this comparison is a qualitative one as it would not be appropriate to compare the yields between nano- and micropowder. Nevertheless were the yields of the products in shown in Fig. 4 determined (SI Tab. 16).

Nickel has long been recognized as a hydrogenation catalyst,^[62]^ – but why does it, when not combined with Fe, only reluctantly reduce NAD (Fig. 2 and ^[20]^) and yet overreduce NMN, not leaving any traceable amount of single-reduced species? The answer probably lies again in the structural differences between NAD and NMN. NMN can be more easily absorbed to a hydrogenated Ni surface, possibly over the entirety of its nicotinamide ring (s. SI Scheme 13). This could also explain the fast formation of its fully reduced product **2d** shown in Fig. 4C. NAD in a staggered formation could only absorb partly on the surface, avoiding overreduction.^[63]^

If this is true, why does Fe not overreduce NMN as readily as Ni? Here, we can reflect on themechanisms postulated by us in Pereira et al.,^[20]^ that Fe both serves as a (less effective) hydrogenation catalyst and a strong electron donor, either via direct electron transfer to the nicotinamide cofactor or the formation of nascent hydride groups on its surface.

Assuming that Fe predominantly reduces NMN through direct electron transfer, the reduction process prioritizes the species with the most favourable redox potential first—namely, 1,4-NMNH (and 1,4-NADH, respectively). This hypothesis was substantiated by cyclic voltammetry measurements, which revealed that among all the reduction products obtained from NMN, 1,4-NMNH exhibits the highest oxidation potential (SI Tab. 14, SI Figs. 19–23, 32 and 33).

Another possible explanation could be that the Ni catalyst does not alter as much as the Fe surface, meaning there would be a constant supply of hydrides available. For Fe, the previously described association with phosphate could block active centers which further prevent over reduction.

While the combination of nickel’s hydrogenation strengths and iron’s electron donation increases the yield of 1,4-NADH immensely compared to only Fe or only Ni (Fig. 2, SI Tabs. 5, SI Fig. 24), the directed reduction of NMN to 1,4-NMNH does not occur in higher yields. In fact, adding nickel to the mix increases overreduction in such a way, that one could argue that NMN by itself could only have played a role as prebiotic hydride acceptor and donor in an environment depleted of Ni (SI Tab. 8, SI Fig. 26). Thus, environmental conditions such as metal availability could have influenced the prebiotic selection process of redox cofactors. This study has shown that the adenosine-derived tail of NAD stabilizes the functional nicotinamide part of the molecule. Like that, the reduction and oxidation properties of NAD could have been maintained within a broader variety of environmental conditions than without the tail. This observation was further validated by performing experiments with both NMN and NAD^+^ in the same reaction mixture using μNi and μFe as metal promoters. Also controls with NAD^+^ and NMN separately were conducted (SI Schemes 5 and 6, SI Tabs. 13 and 14, SI Figs. 34 and 35). For the mixed experiments, both cofactors (each cofactor either 6 or 12 mM) were combined with 30 mM of metal powder, leading to 50:1 and 25:1 metal to cofactor ratios, respectively. In all cases, the 1,4-NADH concentration surpassed that of 1,4-NMNH considerably (SI Scheme 8, SI Tabs. 19 and 20, SI Figs. 40 and 41). This potentially shows how the dinucleotidic structure could have been an advantage on a non-enzymatic level. However intriguing these results, reducing NAD and NMN with the help of H_2_ and metal catalysts is just one part of these cofactors’ role in the prebiotic path towards the first functioning cells – being able to act as a reductant is equally important.

### THE REDUCTION CAPABILITY OF NMNH AND NADH

Recently, it was shown, that Fe^3+^ ions (among other metal ions and also minerals) can promote the reaction of 1,4-NADH with pyruvate to lactate abiotically (SI Scheme 12).^[10]^ Here, we used this reaction as a proxy to compare the reducing capabilities of 1,4-NMNH and 1,4-NADH, showing that both molecules can reduce pyruvate to equal amounts under aqueous conditions with Fe^3+^ in 17 h at 40°C (pH<5), based on recently published experiments by Mayer and Moran.^[10]^ These results underline that the adenosine nucleotide tail does not (or at least not strongly) influence the efficiency of the catalysed hydride transfer (Fig. 5, SI Figs. 63–65)

**Figure 5.**
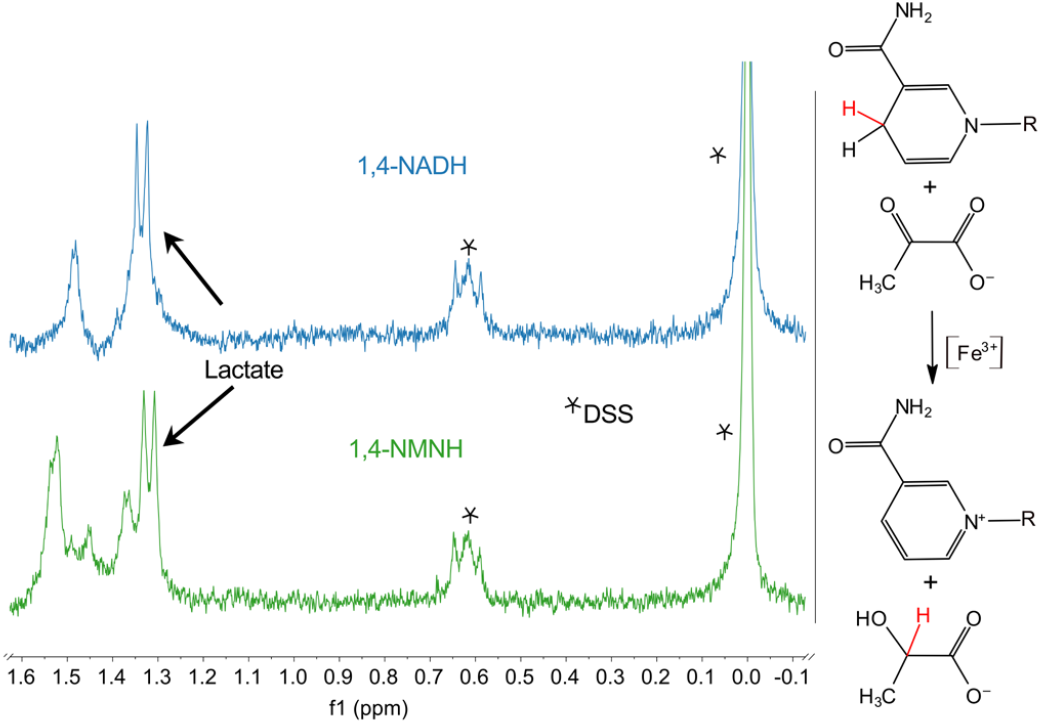
Reduction of pyruvate to lactate with 1,4-NMNH and 1,4-NADH. The method of the reaction was performed according to Mayer and Moran (2023)^[16]^ with the help of Fe(III) chloride as a catalyst in 17 h. Both nicotinamides perform equally well in this reaction. In both cases, lactate forms in comparable amounts – 2 repetitions per reaction. (for original NMR spectra s. SI Figs. 63 and 64).

Investigating the redox potential of both 1,4-NADH and 1,4-NMNH standards via cyclic voltammetry helped comparing their oxidation potential with that of the reaction mixtures of NiFe-assisted reduction of NAD^+^ and NMN with H_2_ (SI Table 12, SI Figs. 19–23, 32 and 33). In the case of NiFe-assisted NAD^+^ reduction, the resulting mixture shows only the oxidation potential of 1,4-NADH, while in the case of NMN reduction, the oxidation potential of both 1,4-NMNH and that of a second reduced species (most likely the species with the 2^nd^ highest concentration, **2c**) is measured. As the second signal has a lower oxidation potential (meaning is harder to oxidize), we postulate that in the case of NMN, as for NAD, the 1,4-species is the most relevant reductant, not only in a biological but also in a prebiotic context.^[64–66]^ One could argue that it is possible that the 1,4 position of an overreduced species would show a similar oxidation potential as a single reduced 1,4-species. However, as the oxidation of the latter leads to the aromatization of the nicotinamide ring, this reaction would be energetically favourable.

This ultimately leads to the conclusion that both 1,4-NMNH and 1,4-NADH are equally good hydride donors and could have both played a role in prebiotic hydride transfer. As the synthesis of both from their oxidized species is highly influenced by effectiveness and mechanism of the hydrogenation catalysts, it is possible that the environment had an important influence on the prebiotic selection of nicotinamides. The conditions described here for the oxidation of 1,4-NADH and 1,4-NMNH (acidic and Fe^3+^ as homogeneous catalyst) can also be found in serpentinizing systems such as the Rainbow hydrothermal vent field in the Mid Atlantic Ridge.^[46]^ How to realistically combine the conditions needed for reduction and oxidation still needs to be shown in the laboratory, but alternating physicochemical conditions on a microcompartment level within serpentinizing systems have been observed and further hypothesized as driving force for prebiotic reactions.^[67–70]^

## Conclusion

In this study, we have shown that both nicotinamide mono- and dinucleotide can be reduced under conditions found in serpentinizing systems, i.e. with H_2_ gas promoted by Fe and Ni containing minerals. We further observed that the distinct structure of the molecules influences the reduction product spectrum associated with the nicotinamide ring.

There is a significant difference in how both molecules react to the addition of nickel to iron-containing catalysts – while most of the overreduction of NMN seems to be due to the contribution of nickel, the specific reduction of NAD^+^ to 1,4-NADH and smaller amounts of 1,6-NADH with Ni-Fe-alloys is much faster than with iron or nickel alone. The overreduction of NMN, only consisting of nicotinamide and a phosphorylated ribose part, inevitably would lead to the molecule being a less effective hydride donor. The rather quick overreduction could be due to NMN being able to interact more directly with the mineral/metal surface than NAD. The additional AMP part attributes NAD its well-described staggered position in solution ^[57–60]^ which could prevent it from associating to the mineral surface the same way NMN does (SI Scheme 13). Further (theoretical) investigation will be necessary to get a definite answer.

Looking at the reverse reaction, only oxidation at the 1,4 position of the nicotinamide ring is observed in a biological context. Prebiotically, a 1,6-species could also be relevant for reduction, but this, to our knowledge, has not yet been observed in an experimental setup. However, it has been shown that the 1,4- species can participate in non-enzymatic reduction of further electron acceptors.^[5,10]^ Here, we additionally confirmed that both 1,4-NMNH and 1,4-NADH act equally well as a hydride source in a non-enzymatic context. A next step would aim for the self- sustaining combination of both reduction and oxidation of these redox cofactors.

When it comes to the concrete question of prebiotic selection of organic cofactors, our study suggests that the ability to provide hydrides might not be the bottleneck, but rather the setup of the environment in which these reactions would have taken place. Overall, in the case of NAD and NMN, the dinucleotide would be more adapt to a variety of environments while the mononucleotide might be advantageous in an environment with less efficient catalysts (i.e. easier to reduce). Of course, more research is necessary to understand the depths of molecular structure in prebiotic selection, including molecules we do not see in biochemistry today.

Lastly, our and other’s observations of a stabilizing function of adenosine-derived tails could be applied other cofactors such as FAD, CoA or SAM. It seems feasible that also there, the extended structure could have been of merit in a prebiotic setting prior to a biological function.^[14,71]^

## Supporting Information

The authors have cited additional references within the Supporting Information.^[72–78]^

## Data Availability Statement

The data that support the findings of this study are available in the Supporting Information of this article. Original analysis files (LC-MS, NMR) will be provided by the corresponding authors upon reasonable request.

## Acknowledgements

D:P:H.P an M.P. thank Bill Martin for discussions and support.

D.P.H.P and M.P thank Alicia Casitas and team for providing access and help to cyclovoltammetry measurements. D.P.H.P. and M.P. thank Kendra Belthle for exploratory surface adsorption measurements and discussions. X.X. thanks Armin Geyer for discussions and the DFG funding for NMR spectrometer NEO600 (Forschungsgroßgeräte project number 508097909). M.P. thanks the Max Planck Society and the International Max Planck Research School ‘Principles of microbial life’ for funding. H.T. thanks MPG, the Volkswagen Foundation (96_742) and Deutsche Forschungsgemeinschaft (TU 315/8-1 / TU 315/8-3). This project was supported by the European Regional Development Fund (ERDF) and the Recovery Assistance for Cohesion and the Territories of Europe (REACT-EU).

## Author contributions

Conceptualization: M.P. & D.P.H.P. Methodology: M.P. & D.P.H.P. Investigation: D.P.H.P., M.P., Z.S, T.B. Validation: D.P.H.P., Z.S. & X.X. Formal analysis: X.X., J.B., N.P., D.P.H.P. & M.P. Resources: H.T., K.V. Writing Original Draft: M.P. & D.P.H.P. Writing Review & Editing: all authors. Visualization: M.P., D.P.H.P. & X.X. Supervision: M.P.

