## Supporting Information for "The dinucleotide structure of NAD enables specific reduction on mineral surfaces"

### Methods

#### Metal preparation

The synthesis of $Ni^0$, $Fe^0$ and Ni-Fe Nanoparticles was carried out as described by Beyazay et al.[1] Commercially bought $Fe^0$ (reduced, <149 µm, Carl Roth, referred to as $\mu Fe^0$) and $Ni^0$ micropowders (3-7 micron, Thermo Scientific, referred to as $\mu Ni^0$) were treated under 5 bar of $H_2$, at 50 °C for 16 h before being used. A detailed characterization of these metals can be found in the same publication.

#### Experimental setup with Ni/Fe alloys

Under anaerobic conditions, using a glovebox (JACOMEX), 3 mL of anoxic 0.133 M phosphate buffer solution (PBS; pH 8.5; potassium phosphate monobasic and sodium phosphate dibasic, Sigma-Aldrich, in HPLC-grade water); bubbled with $N_2$ for 1 h containing 36 or 18 µmol of the organic nicotinamide (Nam) compound $NAD^+$ (>95.0%, TCI) or NMN (100% Uthever, MoleQlar; >98.0%, TCI; **SI Figure 4**) were placed in 5 mL glass vials (beaded rim) with a polytetrafluoroethylene (PTFE)-coated stirring bar. Equimolar amounts (relative to the cofactor) of metal atoms of Fe, $NiFe_3$, NiFe, $Ni_3Fe$, or Ni nanopowders were added to the bottom of each vial, with the exception of a metal-free control. Alternatively, experiments with Fe and Ni micropowders had 1.8 or 7.2 mmols of metal and 36 µmol of cofactor (metal-cofactor ratio 50 or 200:1) in 0.5 M PBS (pH 8.5). The vials were sealed with a crimp cap with a PTFE-coated membrane. To allow gas exchange between the interior and the exterior of the glass vial, a syringe needle was inserted through the crimp cap membrane before the vials were placed in the high-pressure reactor.

**SI Table 1** The molecular weight (MW) was normalized to the amount of metal atoms of each metal powder. The mg of metal used in experiments with 18, 36, and 1800 µmol are listed for each of metal used.

| Metals | MW / metal atom | 18 µmol (mg) | 36 µmol (mg) | 1.8 mmol (mg) |
|---|---|---|---|---|
| Fe | 55.85 | 1.01 | 2.01 | 100.5 |
| $NiFe_3$ | 56.56 | 1.02 | 2.04 | |
| NiFe | 57.27 | 1.03 | 2.06 | |
| $Ni_3Fe$ | 57.98 | 1.04 | 2.09 | |
| Ni | 58.69 | 1.06 | 2.11 | 105.7 |

#### Standard high-pressure reaction

After pressurizing the reactor (Berghof Reactor 300) with either 5 bar of Ar gas (99.999%, Air Liquid) or 5 bar of $H_2$ gas (99.9% Nippon Gases), the reactions were started and regulated by a controlled reactor heating system (Berghof Products + Instruments). Reactions were performed from 1 h to 4 h at 40 °C and 400 rpm, in a Berghof Reactor Heating System (BR-HS). Afterward, reactors were depressurized under anaerobic conditions and the samples (metal powders and solution) were transferred to 2 mL Eppendorf tubes and centrifuged for 20 min, at 4 °C, and 13,000 rpm (Fresco 17 Microcentrifuge). The supernatants were subjected to different analyses, which are described below.

##### Reduction of pyruvate with $Fe^{3+}$ and 1,4-NMNH or 1,4-NADH

These experiments followed the protocol described in Tables S15 and S16 of the paper Mayer et al. 2024.[2] An aqueous mixture of 0.1 mL with 0.1 M pyruvate (Pyruvic acid, Carl Roth), 0.2 M 1,4-NADH (95%, Thermo Scientific), and 0.06 M $FeCl_3$ (98% anhydrous, Grüssing GmbH) reacted overnight at 40 °C and 400 rpm (pH<5). For the removal of metal ions, it was added 0.2 mL of a thiolate/phosphate solution (100 mg NaSH, 100 mg NaOH in 10 mL saturated aqueous $Na_3PO_4$), and left to settle in the fridge (4 °C) for 3 h. Instead of a DMSO standard as used in the referenced protocol, 0.1 mL of a 7mM DSS stock solution was added at the end of the experiment. To reach a certain volume, 0.2 mL of $D_2O$ were also added before the sample was measured. $^1$H-NMR spectra were obtained by an AV III HD 250 MHz Spectrometer with a Double Resonance Broad Band (BBOF) probe head. The same experiment was repeated with 1,4-NMNH (97%, AmBeed) instead of 1,4-NADH.

##### Quantitative proton nuclear magnetic resonance (qNMR) analysis

To monitor reactions, as well as detect and quantify the formation of reduced NADH and side products we established a protocol for quantitative proton nuclear magnetic resonance ($^1$H-NMR).[3,4] The internal standard was a 7 mM solution of sodium 3-(trimethylsilyl)-1-propanesulfonate (DSS, $CH_3$ peak at 0 ppm; >98.0%, TCI) in deuterium oxide ($D_2O$ 99.8 atom%D, AcroSeal, Thermo Scientific), mixed 1:6 with the supernatant of our samples. qNMR spectra were obtained on a Bruker AVANCE-NEO 600 MHz spectrometer equipped with a 5 mm iprobe TBO with z-gradient. Thirty-two scans were made for each sample with a relaxation delay of 40 s (600 MHz) and a spectral width from -3 to 13. Analysis and integration were performed using MestReNova (v.15.0.1).

Metal-free controls (ran under the same conditions as the quantified, metal-containing samples) were used as references to the initial amount of NAD/NMN in the sample to account for evaporation and possible degradation under the given pH, temperature and time. The average initial amount of cofactor in the controls was used as t=0h and to normalize the reaction yields.

##### Product Characterization through 2D-NMR

3 mL of sample from a 1h and 4 h reduction of NMN with equimolar amounts of $NiFe_3$ (5 bar $H_2$, 40 °C, 400 rpm) were dried using a vacuum concentrator (SpeedVac DNA 130, Savant). The remaining solution and pellet were suspended in 500 µl of $D_2O$ to increase the concentration of the products and resolution of the NMR spectra. The same procedure was performed for a $NAD^+$ sample after a 4 h reaction with equimolar amounts of $NiFe_3$ (5 bar $H_2$, 40 °C, 400 rpm). Two-dimensional correlation spectra of $^1$H, $^1$H DQF-COSY (Double-Quantum Filtered COrrelated SpectroscopY), $^1$H, $^1$H TOCSY (Total COrrelated SpectroscopY),$^1$H, $^{13}$C HMBC (Heteronuclear Multiple Bond Correlation spectroscopy) were recorded with standard pulse programs.[5] Edited HSQC (Heteronuclear Single Quantum Coherence spectroscopy) spectra were recorded using sensitivity improvement with echo/anti-echo gradient selection and multiplicity editing during selection step[6,7] NOESY (Nuclear Overhauser Effect SpectroscopY) spectrum was recorded with mixing time of 1.5 s. Chemical shifts are referenced with sodium salt of

trimethylsilylpropanesulfonic acid (DSS). Spectra were obtained with the same instrument as qNMR and compared to a list of possible products.

##### Standards

$^1$H-NMR standards were prepared with 24 mM of the compound and 1 mM of DSS dissolved in $D_2O$ (**SI Figs. 1–4**). The spectra were obtained by a AV III HD 250 MHz Spectrometer with a BBOF probe head.

##### Characterization of 1,4-NMNH through NMR spectroscopy

The sample for NMR measurements was 2 mg of the authentic sample in 0.6 mL of $D_2O$. Experiments were performed on a Bruker AVANCE-NEO 600 MHz spectrometer equipped with a 5 mm iprobe TBO with z-gradient. Two-dimensional correlation spectra of $^1$H, $^1$H DQF-COSY, $^1$H, $^{13}$C HSQC and HMBC were recorded with standard pulse programs.[1] NOESY spectrum was recorded with mixing time of 1.5 s. Chemical shifts are referenced with sodium salt of DSS.

##### Liquid Chromatography Mass Spectroscopy (LC-MS)

Samples were prepared from the supernantant of a reaction with 36 µmol NMN, 36 µmol $Fe^0$ (nanopowder) in 3 mL of 0.133 M PBS (pH 8.5). The reaction ran for 4 h at 40 °C in a high pressure reactor with 5 bar of $H_2$. Afterwards the supernatant was diluted 1:200 with HPLC grade $H_2O$. 2 replicas were measured and compared to the supernatant of the control (without metal powder) and a standard with equivalent amounts of 1,4-NMN to NMN in the samples.

The chromatographic separation was performed on a Thermo Scientific Vanquish high performance liquid chromatography (HPLC) System using a SeQuant ZIC-pHILIC column (150 × 2.1 mm, 5 µm particle size, peek coated, Merck) connected to a guard column of similar specificity (20 × 2.1 mm, 5 µm particle size, Phenomenex) a constant flow rate of 0.1 mL/min with mobile phase A with mobile phase comprised of 10 mM ammonium acetate in water, pH 9, supplemented with medronic acid to a final concentration of 5 µM (A) and 10 mM ammonium acetate in 90:10 acetonitrile to water, pH 9, supplemented with medronic acid to a final concentration of 5 µM (B) at 40 °C .

The injection volume was, dependent on the expected analyte concentration, between 1 µl and 5 µl.

The mobile phase profile consisted of the following steps and linear gradients: 0 – 1 min constant at 75% B; 1 – 6 min from 75 to 40% B; 6 to 9 min constant at 40% B; 9 – 9.1 min from 40 to 75% B; 9.1 to 20 min constant at 75% B.

A Thermo Scientific ID-X Orbitrap mass spectrometer was used in negative ionization mode with an electrospray ionization source and the following conditions: $H^-$ESI spray voltage at 5500 V, sheath gas at 25 arbitrary units, auxiliary gas at 5 arbitrary units, no sweep gas, ion transfer tube temperature at 275 °C, and Vaporizer temperature at 75 °C.

Detection was performed in full scan mode using the orbitrap mass analyzer at a mass resolution of 120 000 in the mass range 330–360 (m/z). The extracted ion chromatograms of the $[M-H]^-$ were generated using Freestyle software (Thermo Scientific) applying a mass accuracy of 5 ppm. Comparative predicted natural isotope distribution was calculated using ChemCalc.[6]

Peak spectra were extracted from the apex of the peak. A Background subtraction was performed from an injection of water using the spectrum extracted at the same retention time.

##### Cyclic Voltammetry

The redox potential of NAD and NMN were compared through cyclic voltammetry performed at 1 mM of each compound in $H_2O$ at 25ºC using as electrolyte disodium hydrogen phosphate and potassium phosphate (0.133 M PBS), at a scan rate of 50 mV/s. A three-electrode electrochemical cell has been used: glassy carbon as the working electrode, platinum wire as an auxiliary electrode, and $Ag/AgNO_3$ (0.01 M) as a reference electrode.

##### Scanning Transmission Electron Microscope (STEM)

In preparation for the STEM measurement, 1 mg of Ni-Fe-particles where dispersed in 400 µL methanol (HPLC grade; Fisher Scientific) using an ultrasonic bath (Bandelin) for 5 s. Subsequently, 20 µL the dispersion where diluted with additional 90 µL of methanol, dispersed in the ultrasonic bath again. A drop of the dispersion was placed on the Cu 300 Mesh lacey carbon grid (EMS) and left to dry. In the case of particles recovered after a reaction, the particles were washed with water (HPLC grade, Fisher Scientific), dried and handled accordingly to pristine particles above. Measurements of drop-cast NiFe particles were carried out using an aberration-corrected JEOL JEM-2200FS STEM equipped with a Bruker XFlash 5060 Energy Dispersive X-ray Spectrometer (EDX) operated in analytical mode with a spatial resolution of approximately 1-2 Ångstroms.

### Henry's Law

**SI Equation 1** Calculation of $H_2$ concentration – Henry's law

$$H^{cp} = \frac{c_a}{p}$$

$H^{cp}$:Henry's constant

$c_a$: concentration

p: pressure

Henry's constant for $H_2$:

$$H^{cp}(H_2, 298\ K) = 7.8\ \times\ 10^{-4} \frac{mol}{L \times atm}$$

Temperature dependency of Henry's constant:

$$H^{cp}(H_2, T_2) =\ H^{cp}(H_2, 298\ K) \times\ e^{(500\left(\frac{1}{T_2} - \frac{1}{298\ K}\right))}$$

### Standards

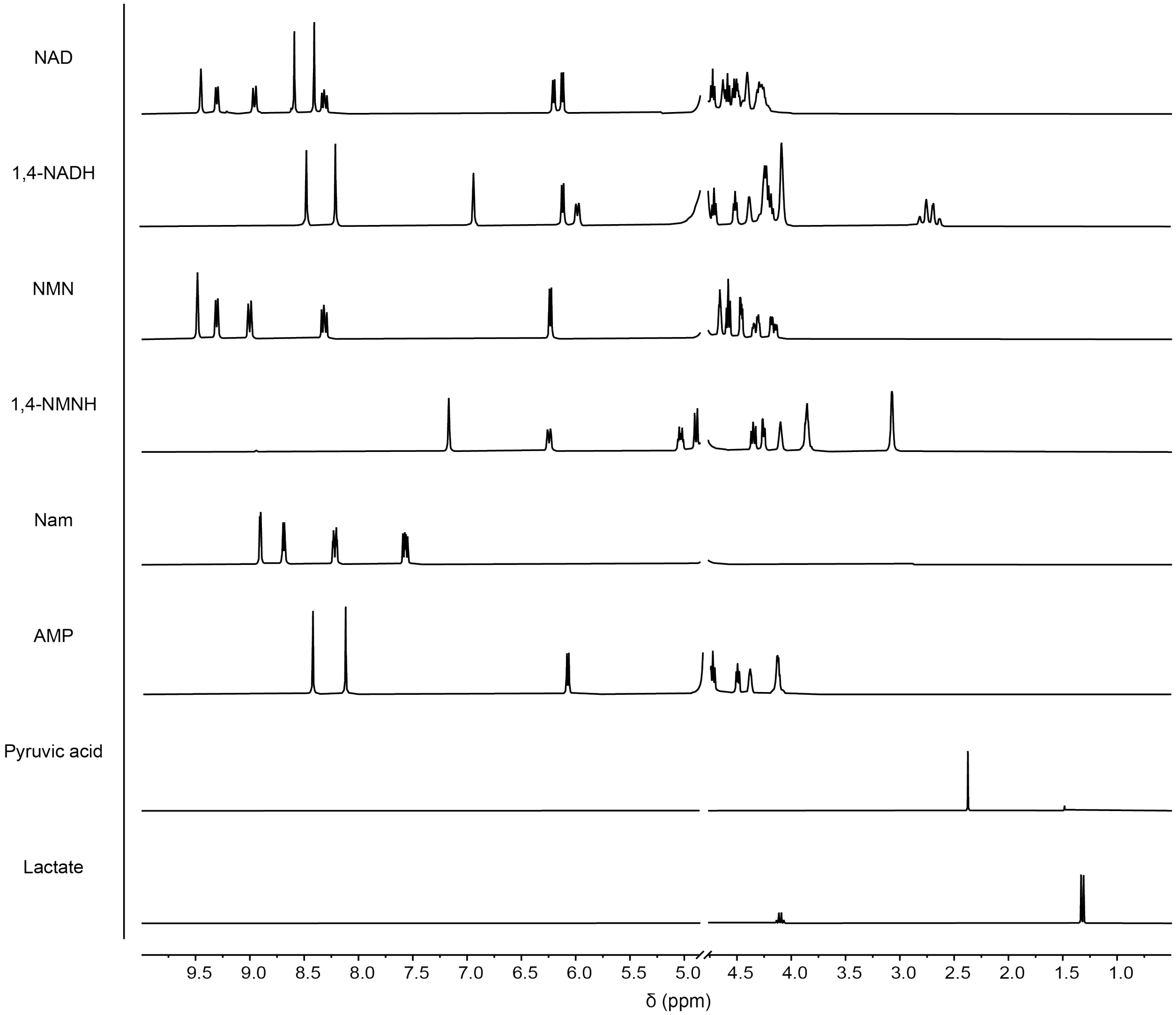

**SI Figure 1** $^{1}H$-NMR standards of several molecules in $D_2O$. Empty spectra was cut out on both sides, as well as the water peak between 4.5 and 5 ppm.

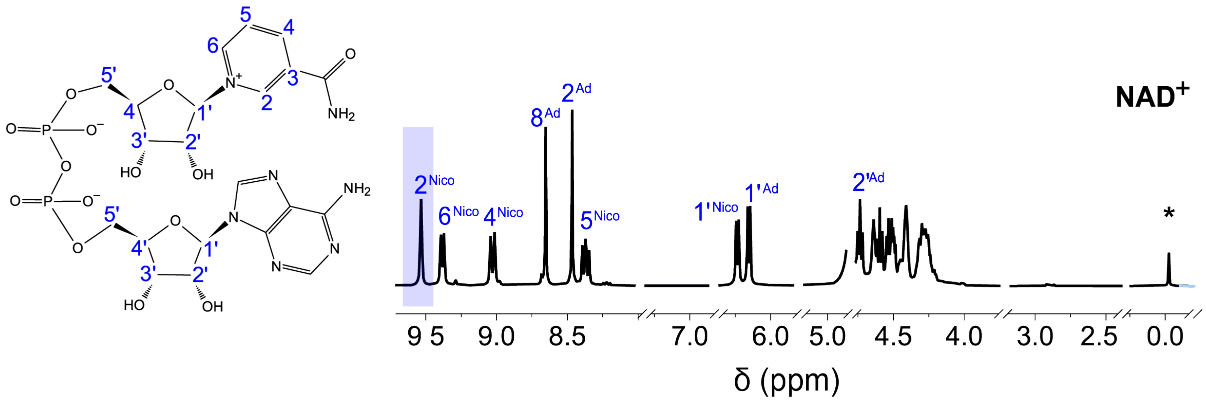

**SI Figure 2** According to 2D-NMR measurements and literature, the [1]H-NMR peaks obtained from $NAD^+$ can be assigned to hydrogens bound to the carbons indicated in the figure. Only spectra without any significant peaks were removed from the figure at several ppm values. Highlighted in blue is the peak used for qNMR. The standard DSS peak at 0 ppm is included and marked (*).

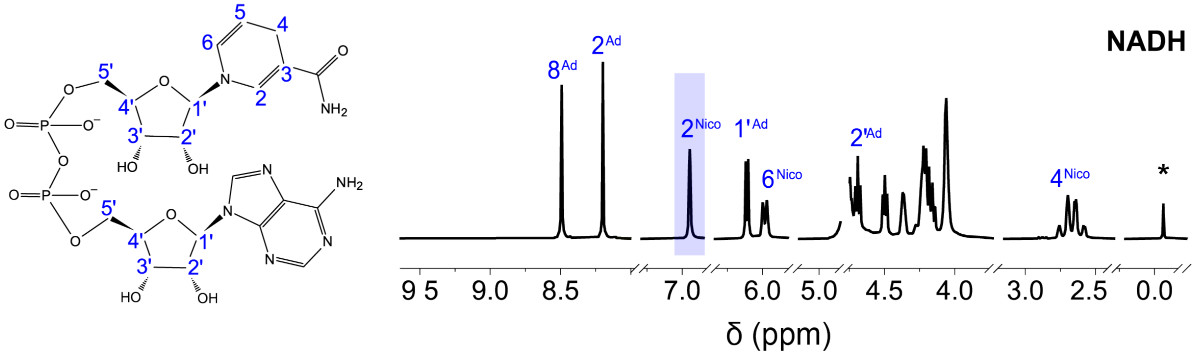

**SI Figure 3** According to 2D-NMR measurements and literature, the [1]H-NMR peaks obtained from 1,4-NADH can be assigned to hydrogens bound to the carbons indicated in the figure. Only spectra without any significant peaks were removed from the figure at several ppm values. Highlighted in blue is the peak used for qNMR. The standard DSS peak at 0 ppm is included and marked (*).

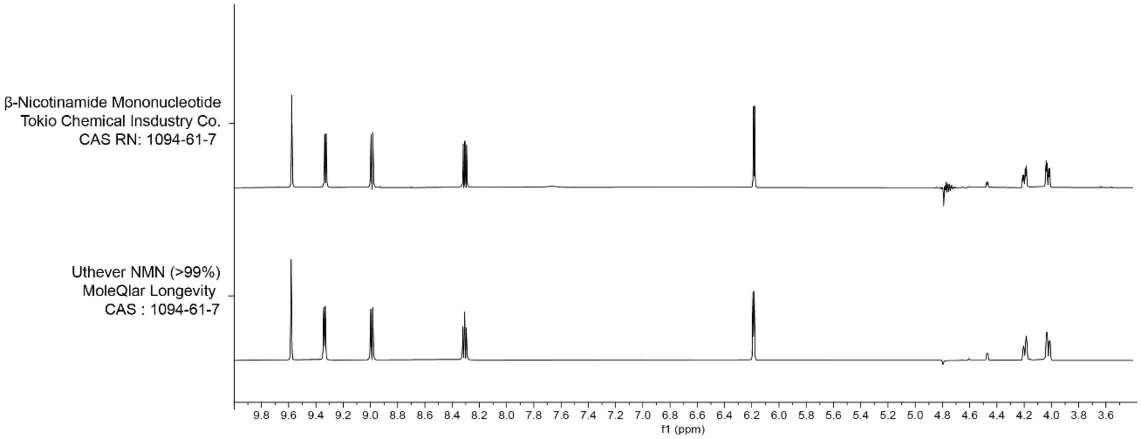

**SI Figure 4** $^{1}$H-NMR comparison of the two different NMN supplies used in all experiments (12 mM of NMN in 0.133 M PBS pH 8.5). Empty spectra was cut out on both extremities, and water suppression hides the peak around 4.8.

#### Characterization of NMN

A full assignment of the $^{1}H$ and $^{13}C$ resonance signals of NMN was fulfilled and the chemical shifts are listed in **SI Table 2**

**SI Table 2** NMR data of NMN in $D_2O$ at 298 K

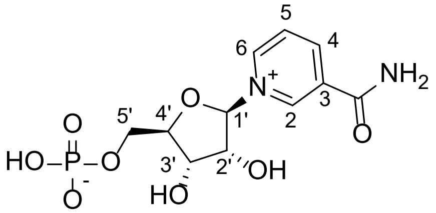

| Position | $\delta_H$ (multi., *J*) | $\delta_C$ |
|---|---|---|
| 2 | 9.47 (s) | 142.6 |
| 3 | | 136.7 |
| C=O | | 168.6 |
| 4 | 8.99 (dt, 8.2, 1.1) | 148.7 |
| 5 | 8.30 (dd, 8.0, 6.4) | 131.3 |
| 6 | 9.29 (d, 6.2) | 145.2 |
| 1' | 6.22 (d, 5.6) | 102.7 |
| 2' | 4.34 (dd, 7.6, 5.3) | 80.5 |
| 3' | 4.45 (dd, 5.1, 2.5) | 73.8 |
| 4' | 4,65 (q, 2.5) | 90.2 (d, 8.8) |
| 5' | 4.31 (ddd, 12.0, 4.3, 2.5) | 66.9 (d, 4.9) |
| | 4.15 (ddd, 12.0, 5.0, 2.1) | |

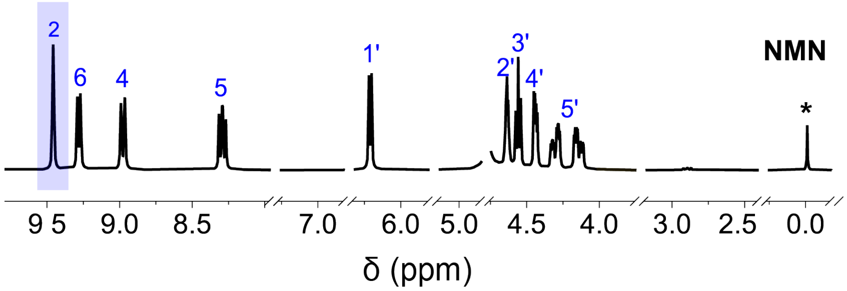

**SI Figure 5** $^{1}H$ spectrum of NMN standard in $D_2O$ with internal reference DSS at 298 K. Empty spectra was removed, highlighting the substrate peaks, labelled according to **SI Table 2**. Highlighted in blue is the peak used for qNMR. The standard DSS peak at 0 ppm is included and marked (*).

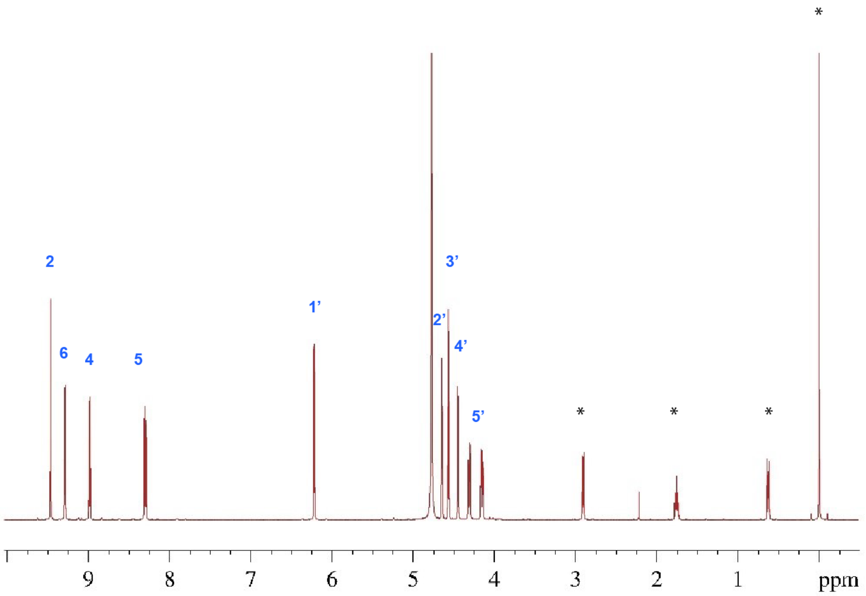

**SI Figure 6** Complete view of the $^1$H spectrum of NMN standard in $D_2O$ with internal reference DSS (*) at 298 K, labelled according to **SI Table 2**. Some $H_2O$ is visible at about 4.8 ppm.

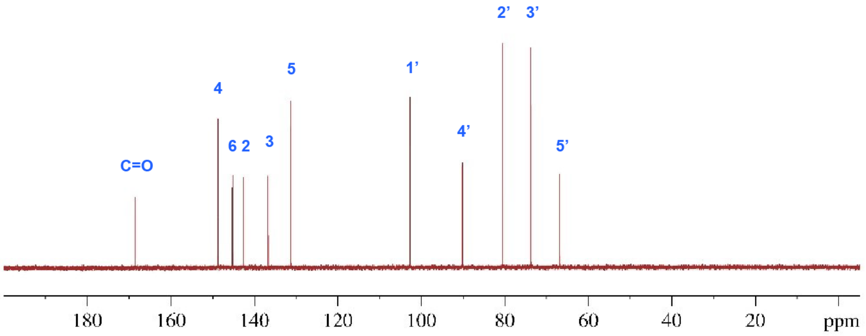

**SI Figure 7** $^{13}$C spectrum of NMN standard in $D_2O$ at 298 K, labelled according to **SI Table 2**.

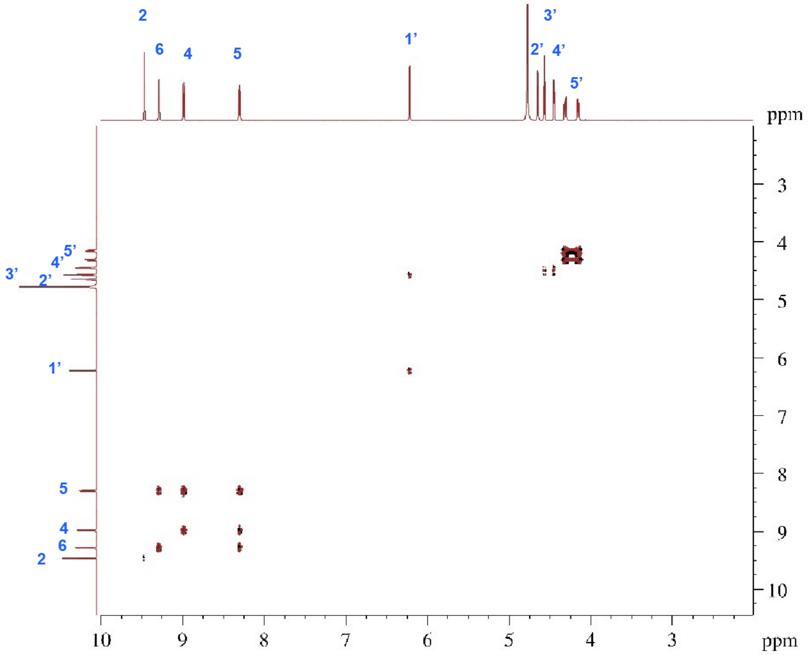

**SI Figure 8** $^{1}H$-$^{1}H$ DQF-COSY spectrum of NMN standard in $D_2O$ at 298 K, labelled according to **SI Table 2**.

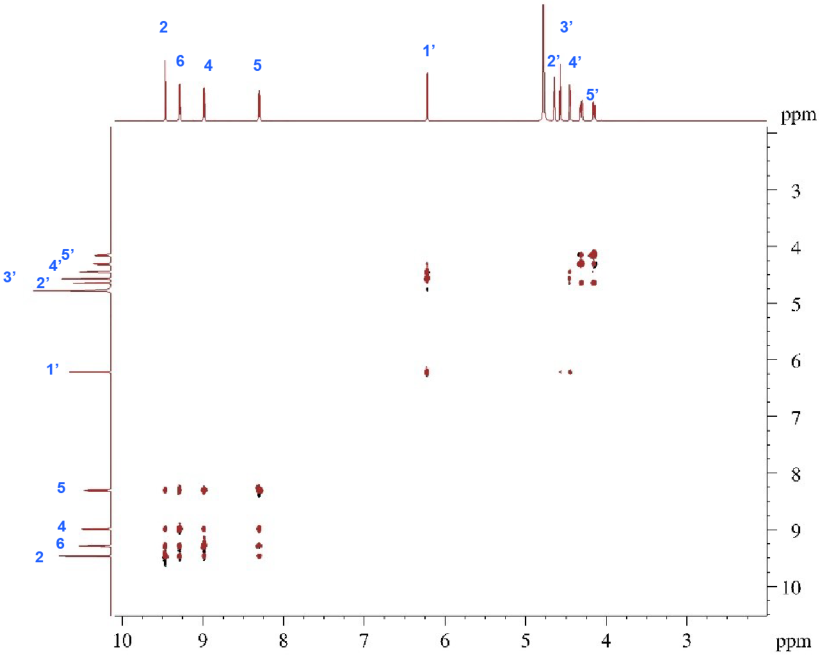

**SI Figure 9** $^{1}H$-$^{1}H$ TOCSY spectrum of NMN standard in $D_2O$ at 298 K, labelled according to **SI Table 2**.

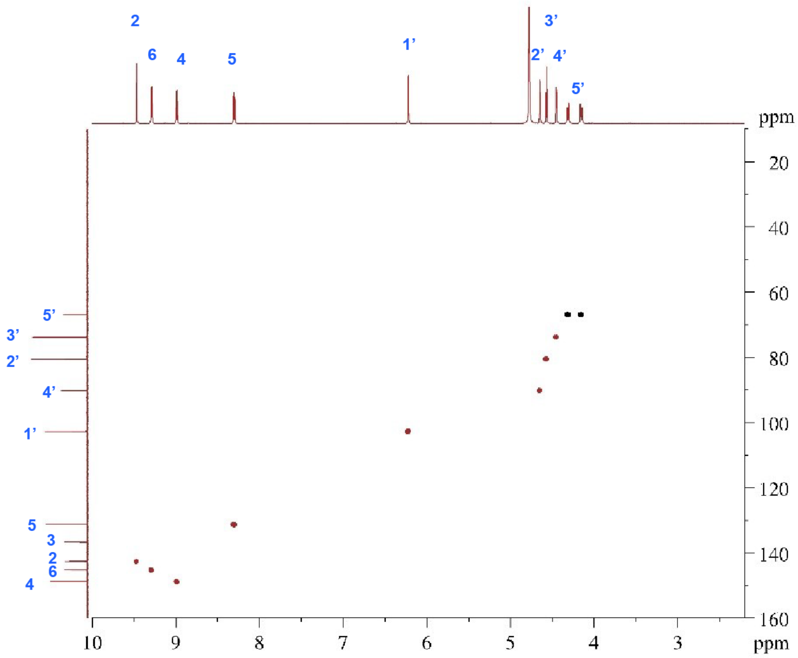

**SI Figure 10** Edited $^{1}H$-$^{13}C$ HSQC spectrum of standard NMN standard in $D_2O$ at 298 K, labelled according to **SI Table 2**.

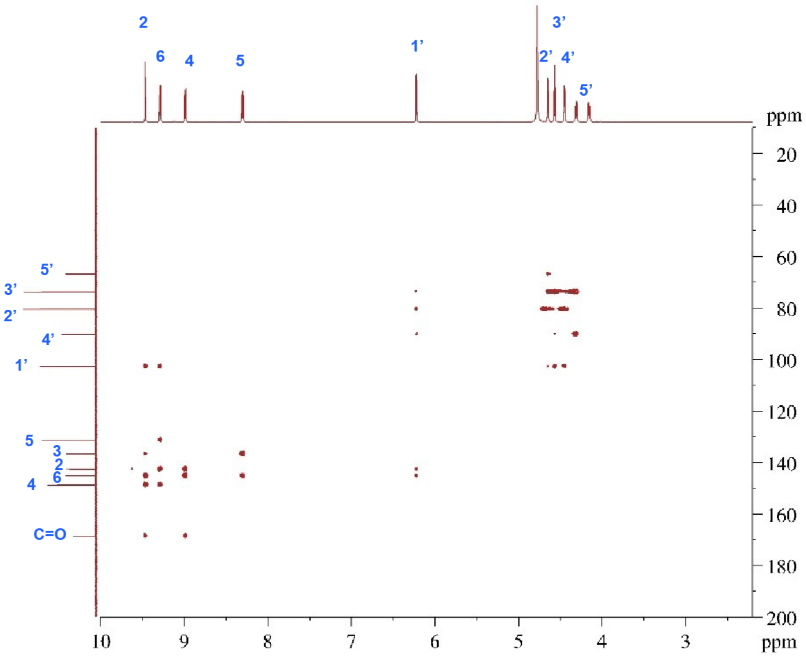

**SI Figure 11** The $^{1}H$-$^{13}C$ HMBC spectrum of NMN standard in $D_2O$ at 298 K, labelled according to **SI Table 2**.

#### Characterization of 1,4-NMNH

A full assignment of the $^{1}H$ and $^{13}C$ resonance signals of 1,4-NMNH was fulfilled and the chemical shifts are listed in.

**SI Table 3** NMR data 1,4-NMNH in $D_2O$ at 298 K

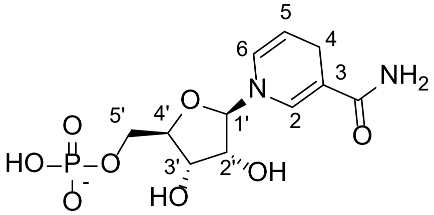

| Position | $\delta_H$ (multi., *J*) | $\delta_C$ |
|---|---|---|
| 2 | 7.15 (d, 1,6) | 141.1 |
| 3 | | 103.4 |
| C=O | | 175.9 |
| 4 | 3.056 (d, 1.7); 3.062 (d, 1.7) | 24.7 |
| 5 | 5.02 (m) | 108.2 |
| 6 | 6.24 (ddt, 8.3, 1,4, 1.5) | 127.7 |
| 1' | 4.88 (m) | 97.7 |
| 2' | 4.34 (dd, 7.6, 5.3) | 73.3 |
| 3' | 4.25 (dd, 5.3, 2.0) | 73.6 |
| 4' | 4.09 (m) | 86.0 (d, 8.8) |
| 5' | 3.87 – 3.81 (m) | 66.6 (d, 4.1) |

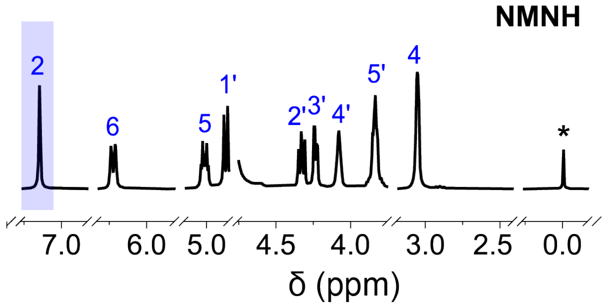

**SI Figure 12** $^{1}H$ spectrum of NMNH standard in $D_2O$ with internal reference DSS at 298 K. Empty spectra was removed, highlighting the substrate peaks, labelled according to **SI Table 3**. Highlighted in blue is the peak used for qNMR.

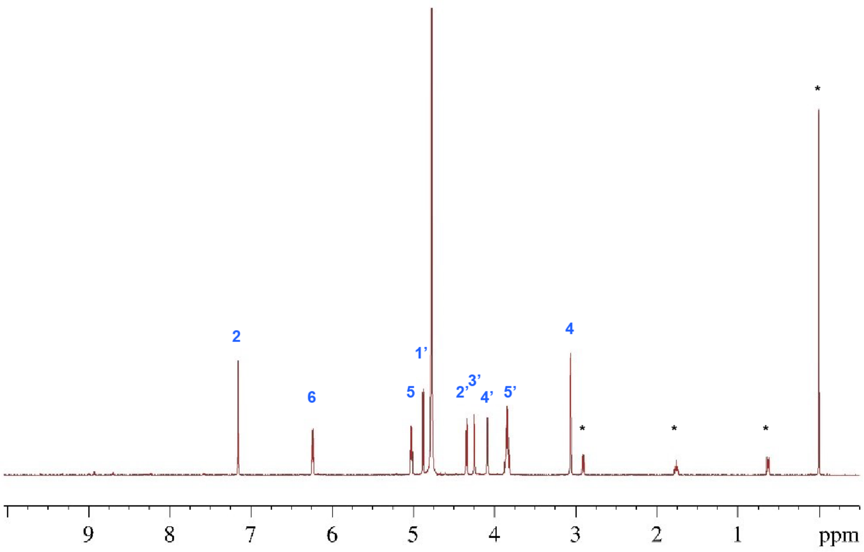

**SI Figure 13** Complete view of the $^1$H spectrum of NMN standard in $D_2O$ with internal reference DSS (*) at 298 K, labelled according to **SI Table 3**. Some $H_2O$ is visible at about 4.8 ppm.

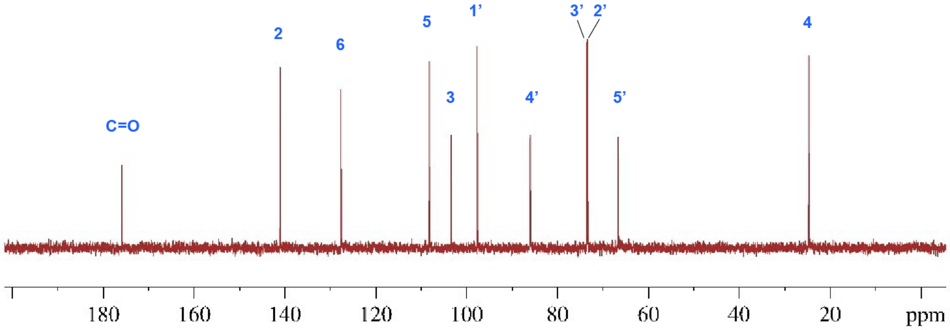

**SI Figure 14** $^{13}$C spectrum of 1,4-NMNH in $D_2O$ at 298 K, labelled according to **SI Table 3**.

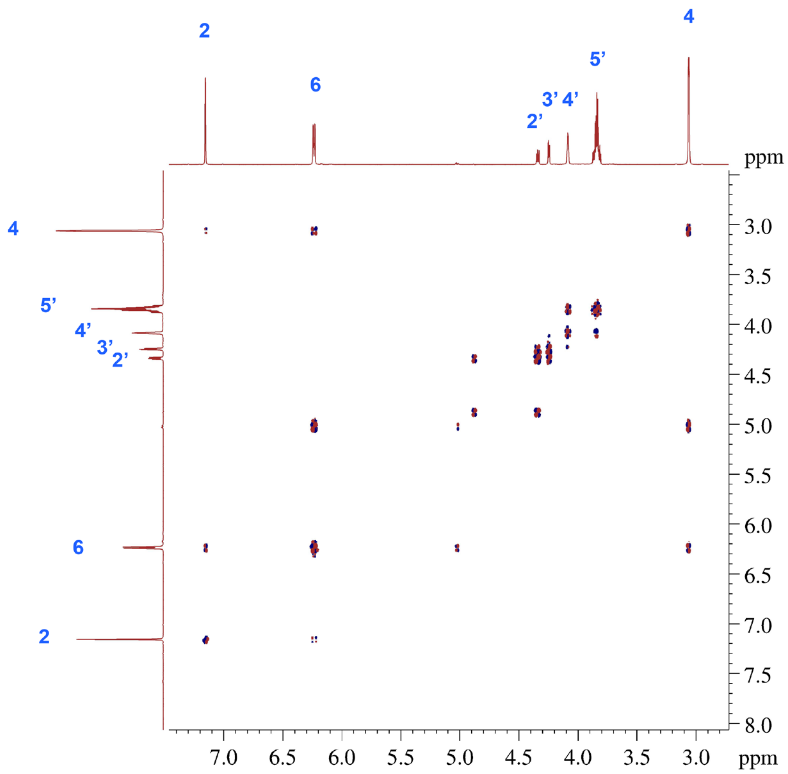

**SI Figure 15** $^1H$-$^1H$ DQF-COSY spectrum of 1,4-NMNH in $D_2O$ at 298 K, labelled according to **SI Table 3**

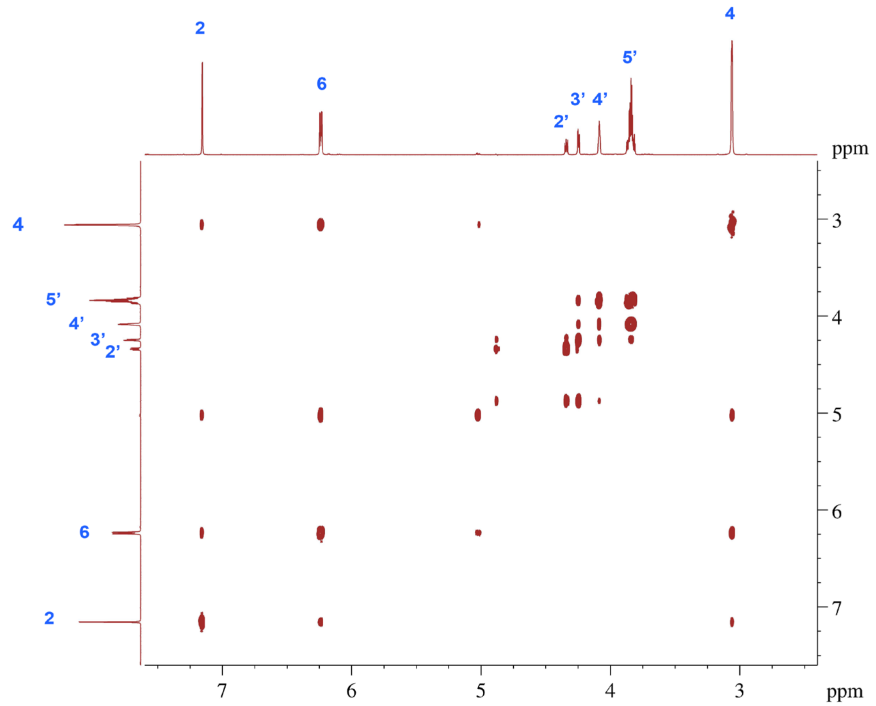

**SI Figure 16** $^1H$-$^1H$ TOCSY spectrum of 1,4-NMNH in $D_2O$ at 298 K, labelled according to **SI Table 3**

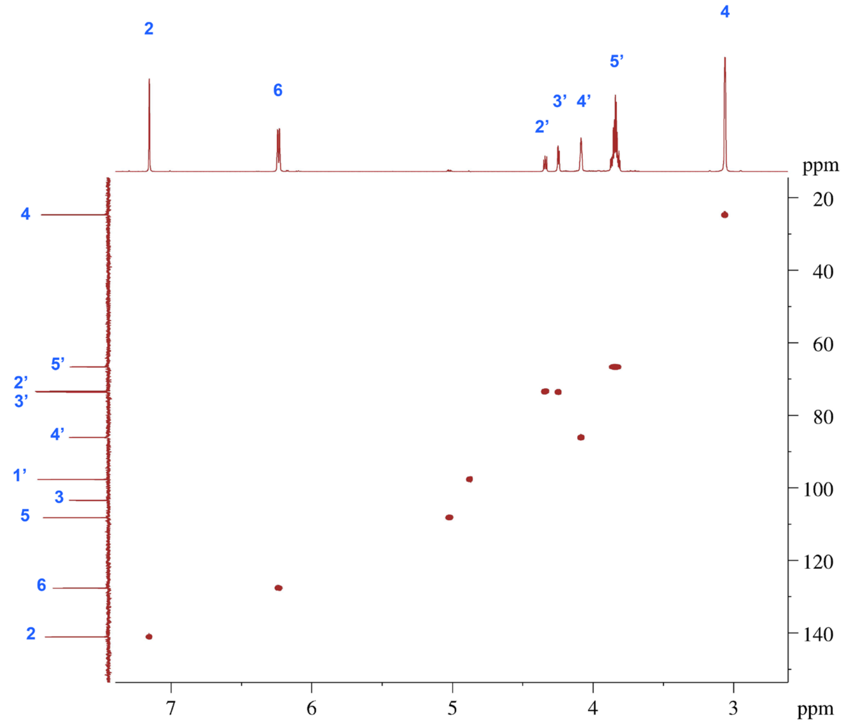

**SI Figure 17** Edited $^{1}H$-$^{13}C$ HSQC spectrum of 1,4-NMNH in $D_2O$ at 298 K, labelled according to **SI Table 3**

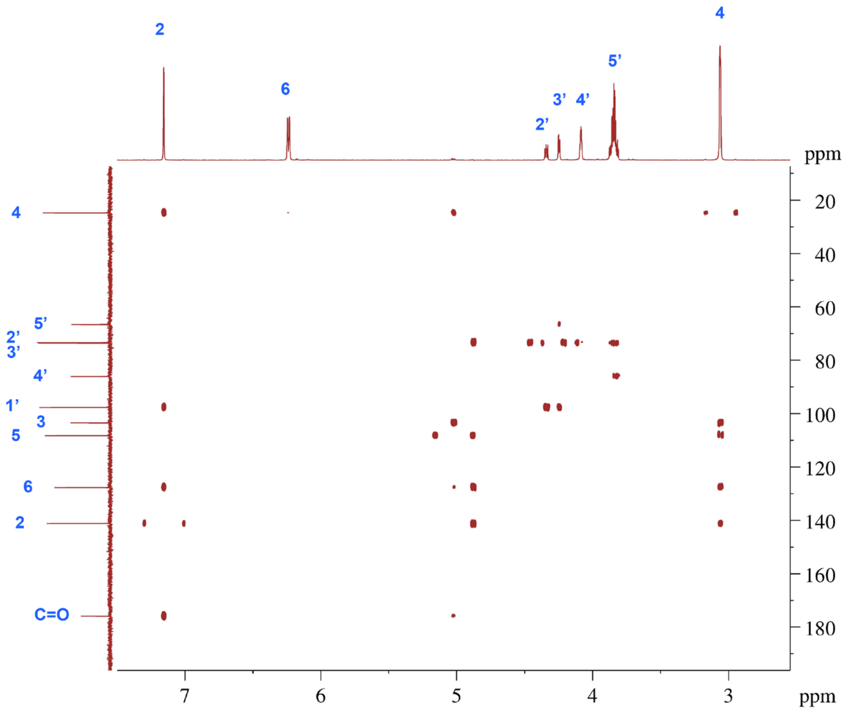

**SI Figure 18** The $^{1}H$-$^{13}C$ HMBC spectrum of 1,4-NMNH in $D_2O$ at 298 K, labelled according to **SI Table 3**

#### Cyclic Voltammetry standards

Standards were prepared at 1 mM of each compound in PBS (0.133 M, pH 8.5) at 25ºC, a three-electrode electrochemical cell was prepared for cyclic voltammetry with glassy carbon as working electrode, platinum wire as auxiliary electrode and $Ag/AgNO_3$ (0.01 M) as reference electrode.

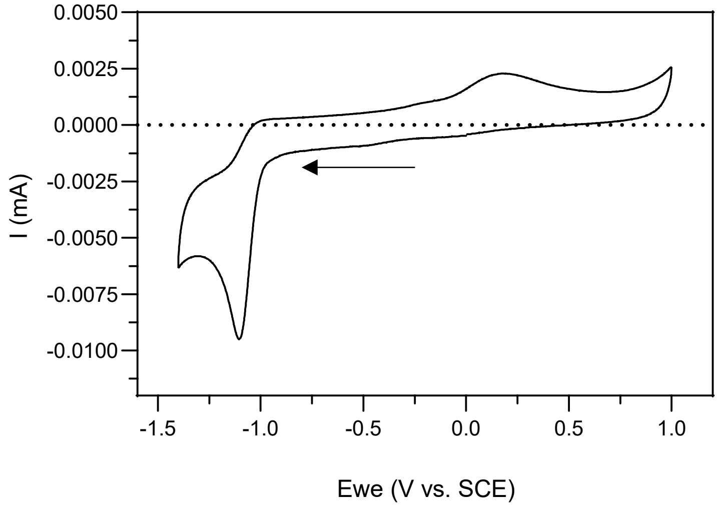

**SI Figure 19** Cyclic voltammogram of $NAD^+$ (1 mM) in PBS (pH 8.5) at 25 ºC.

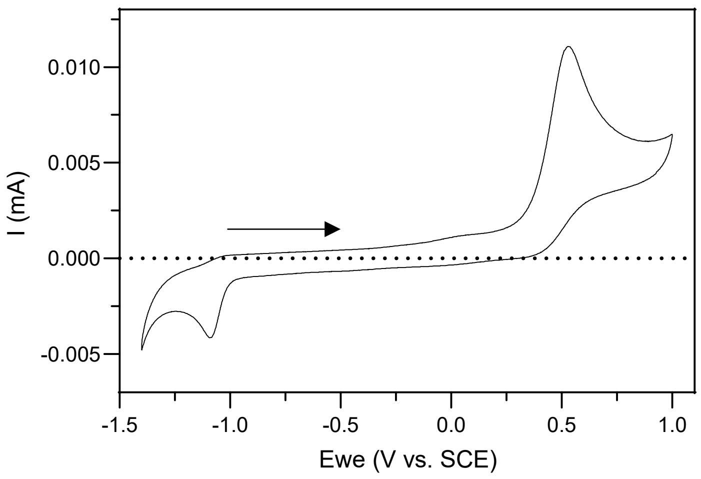

**SI Figure 20** Cyclic voltammogram of 1,4-NADH (1 mM) in PBS (pH 8.5) at 25 ºC.

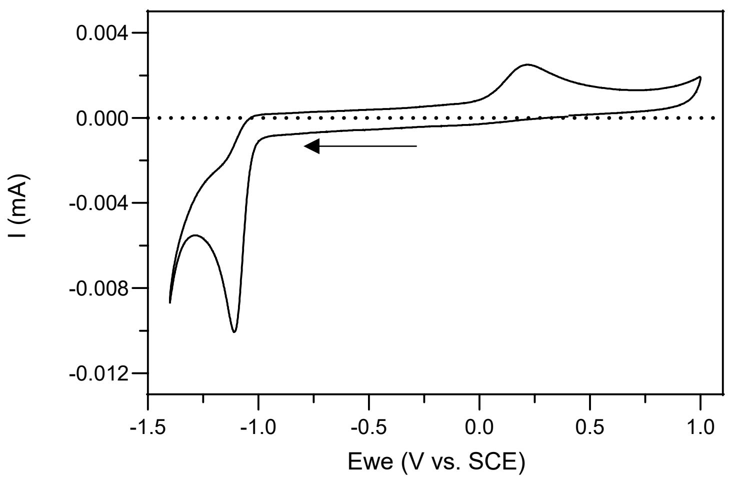

**SI Figure 21** Cyclic voltammogram of NMN (1 mM) in PBS (pH 8.5) at 25 ºC.

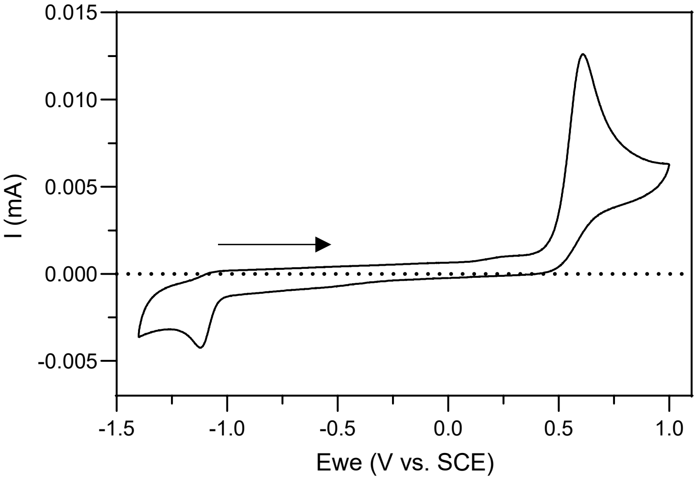

**SI Figure 22** Cyclic voltammogram of 1,4-NMNH (1 mM) in PBS (pH 8.5) at 25 ºC.

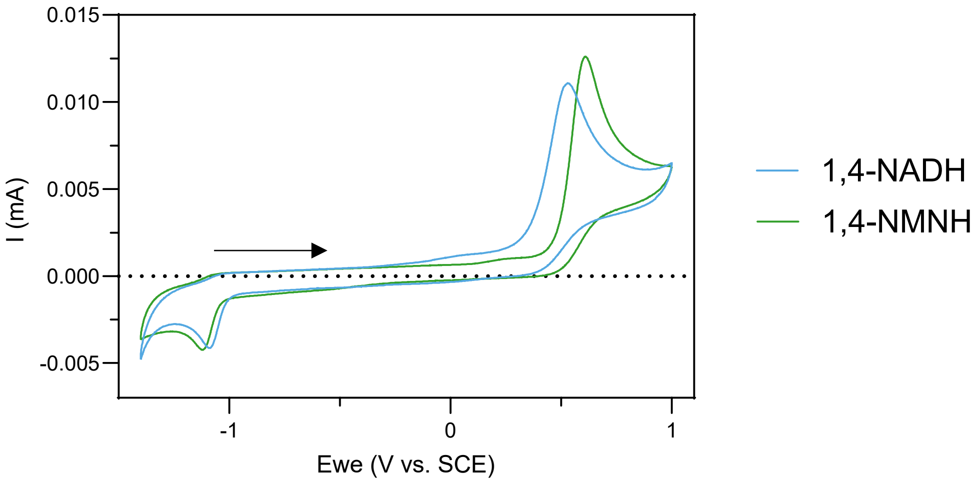

**SI Figure 23** Cyclic voltammogram of 1,4-NADH (1 mM; blue) and 1,4-NMNH (1 mM; green) in PBS (pH 8.5) at 25 ºC.

### Heterogeneous catalysis of $NAD^+$ reduction with $H_2$ and Ni/Fe alloys

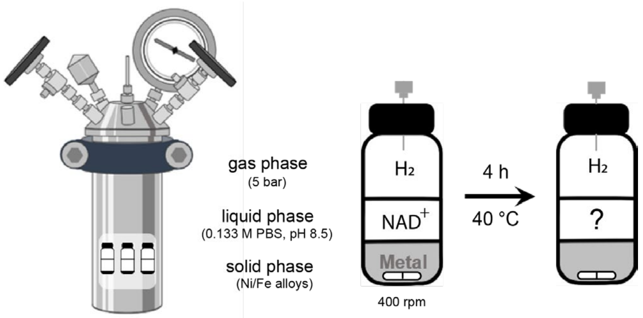

**SI Scheme 1** The reduction of $NAD^+$ with Ni/Fe alloys was tested with the protocol described in detail in Methods, and according to the scheme above. The amount of metal and $NAD^+$ was 36 μmol, which reacted together for 4 h, at 40 °C, under alkaline conditions and 5 bars of $H_2$. The same reaction was made under Ar as a control.

**SI Table 4** For quantification of each compound, a peak was selected in their spectra. The values listed a bellow indicate the ppm value where the peak can be found at approximately pH 8.5 (0.133M PBS).

| Molecule | $NAD^+$ | Nam | 1,6-NADH | 1,4-NADH |
|---|---|---|---|---|
| δ (ppm) for quantification | 9.12 | 8.93 | 7.11 | 6.94 |

**SI Table 5** After 4 h under 5 bar of $H_2$, samples with different nanoparticular Ni/Fe alloys yielded different amounts of 1,4-NADH, 1,6-NADH and nicotinamide (Nam), from the starting material $NAD^+$, as listed below. The starting metal and cofactor were 36 μmol mixed in 3 mL of 0.133 M PBS (pH 8.5), as shown in **SI Scheme 1**. The yields were calculated relative to the metal-free sample (100% $NAD^+$). Reactions with nNi, $nNi_3Fe$, and nNiFe were performed with six replicas; $Fe^0$ and w/o metal were done with five replicas; $nNiFe_3$ amounted to four. To determine the turn over frequency (TOF) of each reaction, 1,4-NADH and 1,6-NADH were considered as products and the total amount of metal atoms were considered as the amount of catalyst, instead of the number of molecules.

| | $H_2$ | $NAD^+$ | SD | 1,4-NADH | SD | 1,6-NADH | SD | Nam | SD | TOF [mol/s] |
|---|---|---|---|---|---|---|---|---|---|---|
| 4h | $nNi^0$ | 72.63% | 3.5% | 3.68% | 0.5% | 1.19% | 0.2% | 6.37% | 0.3% | 3.57E-06 |
| | $nNi_3Fe$ | 61.20% | 2.7% | 14.09% | 1.8% | 6.84% | 1.1% | 7.47% | 0.2% | 1.54E-05 |
| | nNiFe | 31.10% | 2.1% | 19.66% | 2.4% | 7.90% | 1.2% | 7.88% | 0.7% | 2.02E-05 |
| | $nNiFe_3$ | 25.59% | 4.4% | 40.60% | 1.8% | 16.70% | 0.6% | 4.48% | 0.1% | 3.02E-05 |
| | $nFe^0$ | 52.23% | 7.6% | 6.01% | 1.1% | 0.00% | 0.0% | 7.04% | 0.6% | 4.41E-06 |

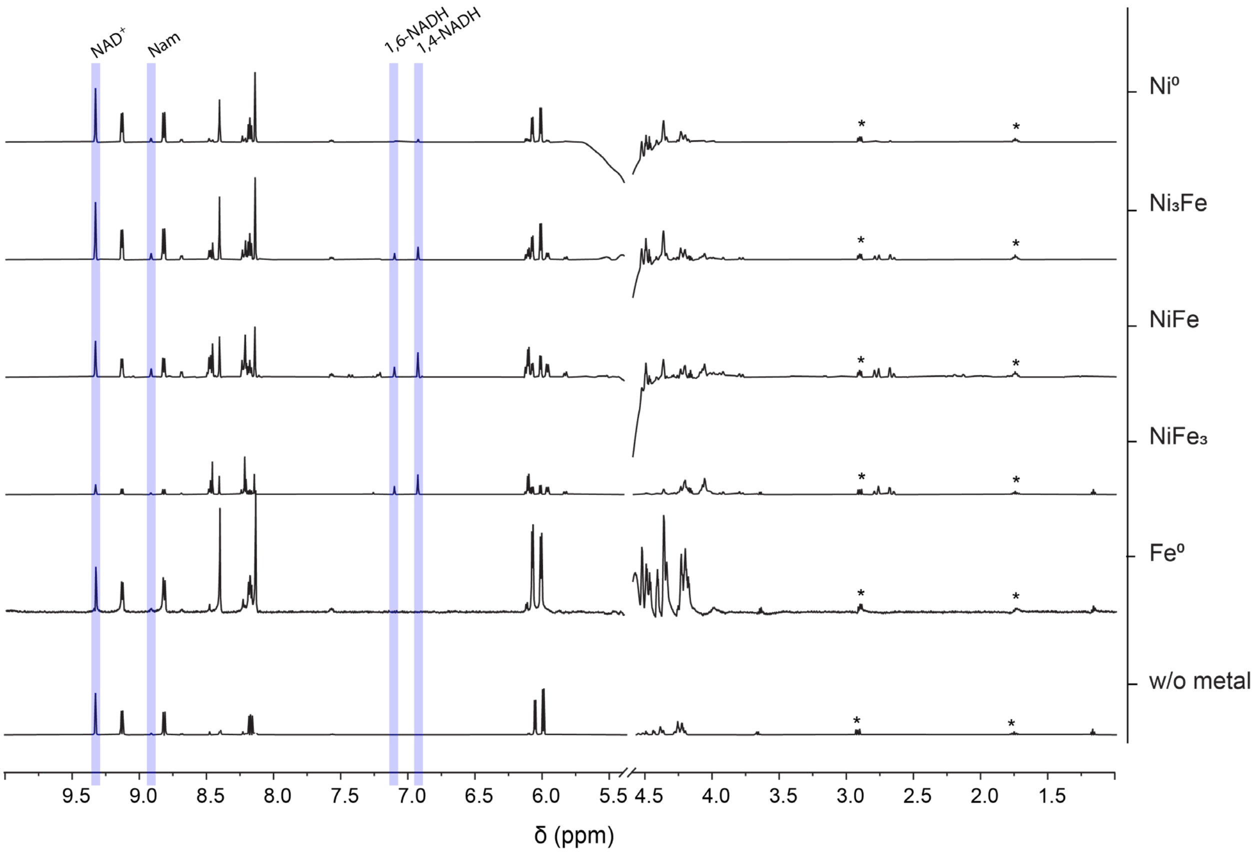

**SI Figure 24** The NMR spectra of replica samples of $NAD^+$ in PBS (0.133 M, pH 8.5) after 4 h with 5 bar of $H_2$ and a metal 1:1 cofactor ratio, as shown in **SI Scheme 1**, are stacked together in this figure. The metal used in each sample is indicated on the right side of each spectra. After 4h reacting, the supernatant was collected and DSS added as an internal standard. The spectra were edited to only include relevant peaks, having been removed a DSS peak at 0 ppm and water peak at 4.8 ppm. No other peaks were found in the areas removed. Some DSS peaks are still visible (*). The peaks used for qualitative analysis and subsequent qNMR are highlighted in blue, according to **SI Table 4**.

**SI Table 6** After 4 h under 5 bar of Ar, samples with different nanoparticular Ni/Fe alloys yielded different amounts of 1,4-NADH, 1,6-NADH and Nam, from the starting material $NAD^+$, as listed below. The starting metal and cofactor were 36 µmol mixed in 3 mL of 0.133 M PBS (pH 8.5), as shown in **SI Scheme 1** with Ar. The yields were calculated relative to the metal-free sample (100% $NAD^+$). Reactions with nNi, $nNi_3Fe$, and nNiFe were performed with six replicas; $nFe^0$ and w/o metal were done with five replicas; $nNiFe_3$ amounted to four. To determine the TOF of each reaction, 1,4-NADH and 1,6-NADH were considered as products and the total amount of metal atoms were considered as the amount of catalyst, instead of the number of molecules.

| | Ar | **$NAD^+$** | **SD** | **1,4-NADH** | **SD** | **1,6-NADH** | **SD** | **Nam** | **SD** | **TOF [mol/s]** |
|---|---|---|---|---|---|---|---|---|---|---|
| 4h | $nNi^0$ | 80.94% | 1.7% | 0.00% | 0.0% | 0.00% | 0.0% | 7.48% | 0.2% | 0.00E+00 |
| | $nNi_3Fe$ | 62.68% | 8.0% | 0.00% | 0.0% | 0.00% | 0.0% | 6.61% | 1.5% | 0.00E+00 |
| | nNiFe | 85.24% | 2.5% | 0.00% | 0.0% | 0.00% | 0.0% | 8.48% | 0.3% | 0.00E+00 |
| | $nNiFe_3$ | 88.84% | 1.2% | 4.43% | 0.9% | 1.64% | 0.3% | 4.55% | 0.0% | 3.20E-06 |
| | $nFe^0$ | 77.00% | 6.0% | 5.44% | 0.9% | 0.00% | 0.0% | 20.33% | 2.3% | 3.99E-06 |

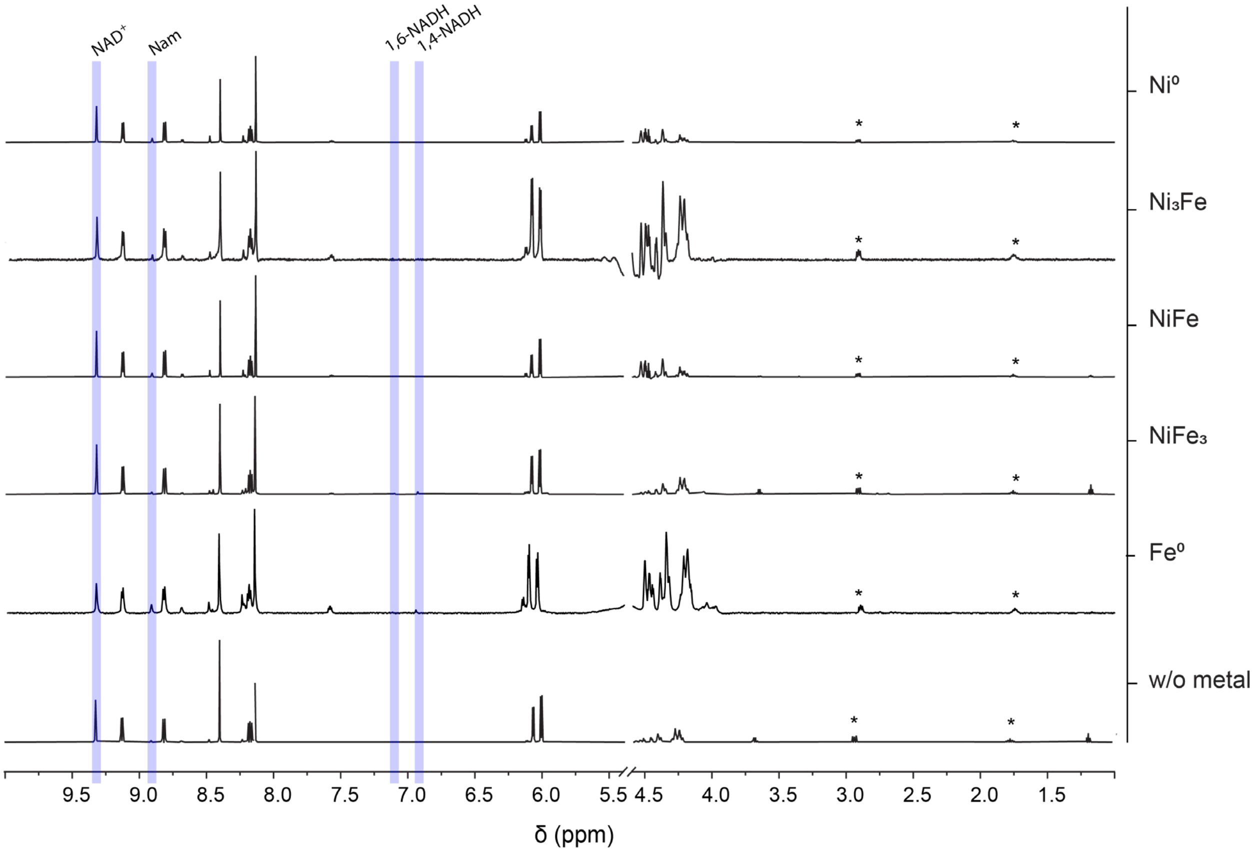

**SI Figure 25** The NMR spectra of replica samples of $NAD^+$ in PBS (0.133 M, pH 8.5) after 4 h with 5 bar of Ar and a metal 1:1 cofactor ratio, as shown in **SI Scheme 1** with Ar, are stacked together in this figure. The nanoparticular metalpowder used in each sample is indicated on the right. After 4h reacting, the supernatant was collected and DSS added as an internal standard. The spectra were edited to only include relevant peaks, having been removed a DSS peak at 0 ppm and water peak at 4.8 ppm. No other peaks were found in the areas removed. Some DSS peaks are still visible (*). The peaks used for qualitative analysis and subsequent qNMR are highlighted in blue, according to **SI Table 4**.

### Heterogeneous catalysis of NMN reduction with $H_2$ and Ni/Fe alloys

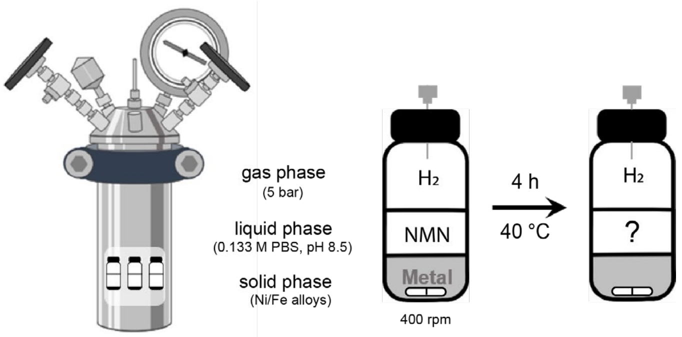

**SI Scheme 2** The reduction of NMN with nanoparticular Ni/Fe alloys was tested with the protocol described in detail in Methods, and according to the scheme above. The amount of metal and NMN was 36 or 18 µmol, which reacted together for 4 h, at 40 °C, under alkaline conditions and 5 bars of $H_2$. The same reaction was made under Ar as a control.

**SI Table 7** For quantification of each compound, a peak was selected in their spectra. The values listed a bellow indicate the ppm value where the peak can be found at approximately pH 8.5 (0.133M PBS).

| Molecule | NMN | Nam | 1,4,6-NMN | NMNH$_2$OH | 1,4-NMNH | 1,2,4,6-NMN |
|---|---|---|---|---|---|---|
| δ (ppm) for quantification | 9.58 | 8.93 | 7.41/7.45 | 7.34 | 7.15 | 2.85 |

**SI Table 8** After 4 h under 5 bar of $H_2$, as shown in **SI Scheme 2**, samples with different nanoparticular Ni/Fe alloys yielded different amounts of 1,4-NMNH, 1,4,6-NMNH, NMNH$_2$OH, 1,2,4,6-NMNH$_5$ and nicotinamide (Nam), from the starting material NMN, as listed below. The starting metal and cofactor were 36 or 18 µmol mixed in 3 mL of 0.133 M PBS (pH 8.5). The yields were calculated relative to the metal-free sample (100% NMN). To determine the TOF of each reaction, the sum of 1,4-NMNH, 1,4,6-NMNH$_3$ and 1,2,4,6-NMNH$_5$ was considered as the amount of product, and the total amount of metal atoms were considered as the mount of catalyst, instead of the number of molecules. Each condition was tested in duplicates and a metal-free control.

| | $H_2$ | NMN | SD | 1,4-NMNH | SD | 1,4,6-NMNH$_3$ | SD | NMNH$_2$OH | SD | 1,2,4,6-NMNH$_5$ | SD | Nam | SD | TOF [mol/s] |
|---|---|---|---|---|---|---|---|---|---|---|---|---|---|---|
| 4h | nNiFe | 9.61% | 6.4% | 7.74% | 0.5% | 21.63% | 3.2% | 25.39% | 0.6% | 3.24% | 0.6% | 1.93% | 0.3% | 2.12E-05 |
| | nNiFe$_3$ | 2.01% | 0.1% | 9.46% | 0.4% | 25.22% | 0.7% | 29.49% | 0.0% | 2.73% | 0.4% | 1.75% | 0.1% | 1.12E-05 |
| | nFe$^0$ | 40.51% | 5.1% | 13.23% | 1.0% | 2.29% | 0.3% | 19.47% | 2.1% | 0.00% | 0.0% | 4.63% | 0.2% | 2.63E-05 |

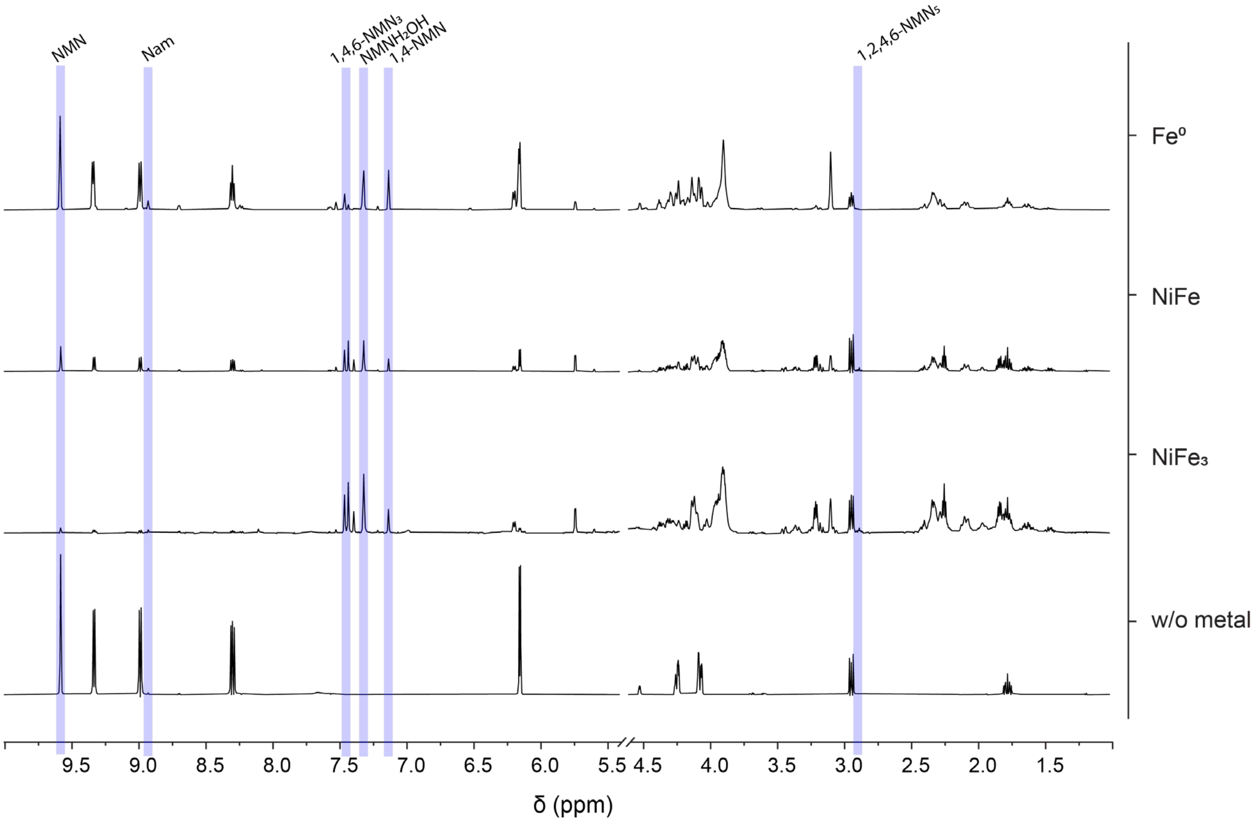

**SI Figure 26** The NMR spectra of replica samples of NMN in PBS (0.133 M, pH 8.5) reacted for 4h with 5 bar of $H_2$ and a metal 1:1 cofactor ratio, as shown in **SI Scheme 2**, are stacked together in this figure. The metal used in each reaction is specified on the right of each spectra. After the 4h reaction, the supernatant was collected and DSS added as an internal standard. The spectra were edited to only include relevant peaks, having been removed a DSS peak at 0 ppm and water peak at 4.8 ppm. No other peaks were found in the areas removed. Some DSS peaks are still visible (*). The peaks used for qualitative analysis and subsequent qNMR are highlighted in blue, according to **SI Table 7**.

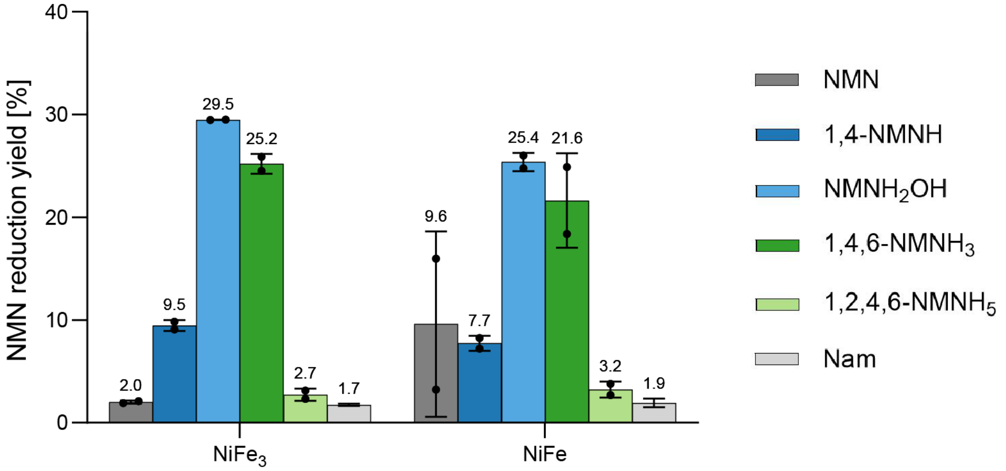

**SI Figure 27** Reduction of NMN in PBS (0.133 M, pH 8.5) after 4 h at 40 °C with equimolar amounts of $nNiFe_3$ or NiFe (**SI Table 1**) and 5 bar of $H_2$. The supernatant was separated from the metal for qNMR (600 MHz) and the yields calculated as for **SI Table 8** and **SI Table 9**.

**SI Table 9** After 4 h under 5 bar of Ar, as shown in **SI Scheme 2** with Ar, samples with different nanoparticular Ni/Fe alloys yielded different amounts of 1,4-NMNH, 1,4,6-NMNH, $NMNH_2OH$, 1,2,4,6-$NMNH_5$ and nicotinamide (Nam), from the starting material NMN, as listed below. The starting metal and cofactor were 36 or 18 µmol mixed in 3 mL of 0.133 M PBS (pH 8.5). The amount of metal atoms was the same as the cofactor. The yields were calculated relative to the metal-free sample (100% NMN). To determine the TOF of each reaction, the sum of 1,4-NMNH, 1,4,6-$NMNH_3$ and 1,2,4,6-$NMNH_5$ was considered as the amount of product, and the total amount of metal atoms were considered as the mount of catalyst, instead of the number of molecules.

| | Ar | NMN | SD | 1,4-NMNH | SD | 1,4,6-$NMNH_3$ | SD | $NMNH_2OH$ | SD | 1,2,4,6-$NMNH_5$ | SD | Nam | SD | TOF [mol/s] |
|---|---|---|---|---|---|---|---|---|---|---|---|---|---|---|
| 4h | nNiFe | 65.07% | 1.0% | 4.40% | 0.4% | 2.92% | 0.0% | 7.52% | 0.3% | 0.00% | 0.0% | 1.90% | 0.1% | 4.75E-06 |
| | $nNiFe_3$ | 54.40% | 0.2% | 6.72% | 0.4% | 3.39% | 0.4% | 13.31% | 0.4% | 0.00% | 0.0% | 2.38% | 0.2% | 6.56E-06 |
| | $nFe^0$ | 87.37% | 1.2% | 0.97% | 0.0% | 0.00% | 0.0% | 0.80% | 0.2% | 0.00% | 0.0% | 5.36% | 0.2% | 1.33E-06 |

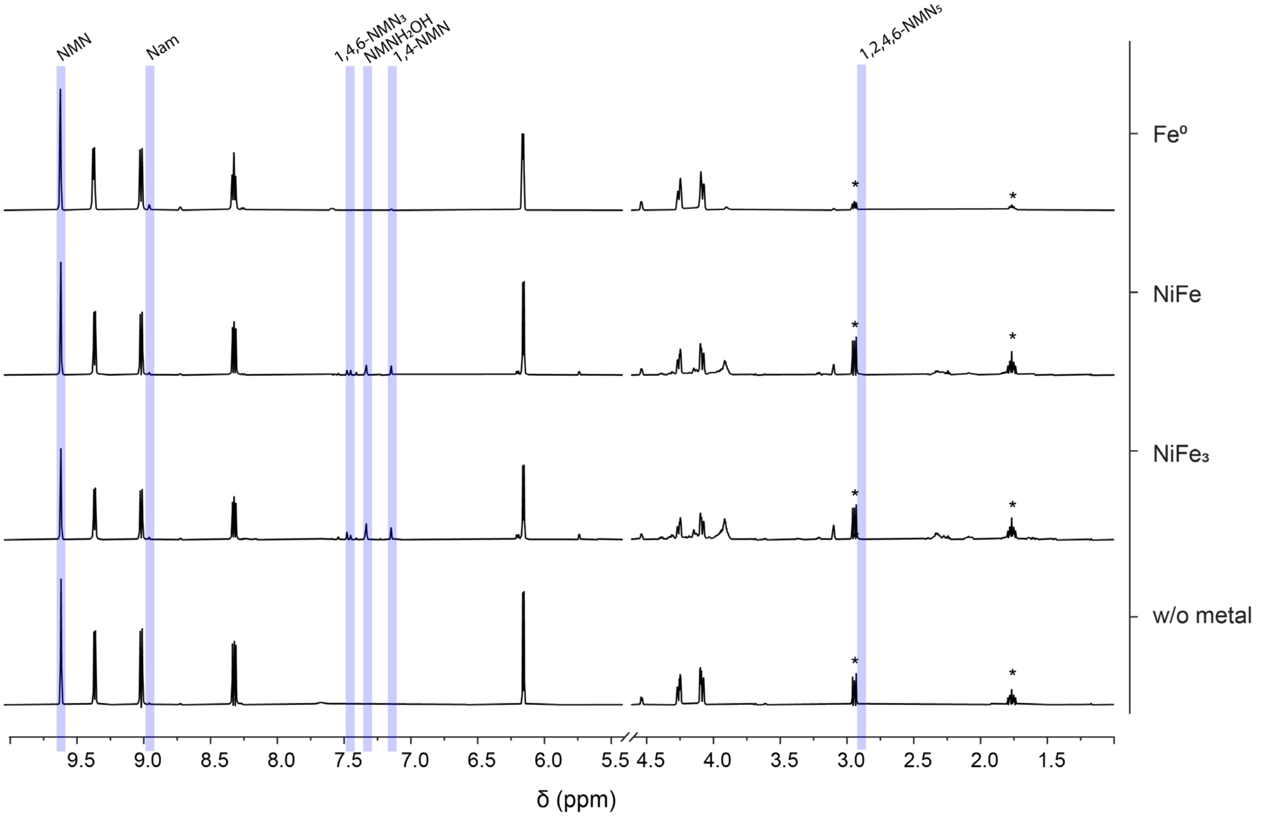

**SI Figure 28** The NMR spectra of replica samples of NMN in PBS (0.133 M. pH 8.5) reacted for 4h with 5 bar of Ar and a metal 1:1 cofactor ratio, as shown in **SI Scheme 2** with Ar, as indicated on the left side, are stacked together in this figure. The metal used in each reaction is specified on the right of each spectra. After the 4h reaction, the supernatant was collected and DSS added as an internal standard. The spectra were edited to only include relevant peaks, having been removed a DSS peak at 0 ppm and water peak at 4.8 ppm. No other peaks were found in the areas removed. Some DSS peaks are still visible (*). The peaks used for qualitative analysis and subsequent qNMR are highlighted in blue, according to **SI Table 7**.

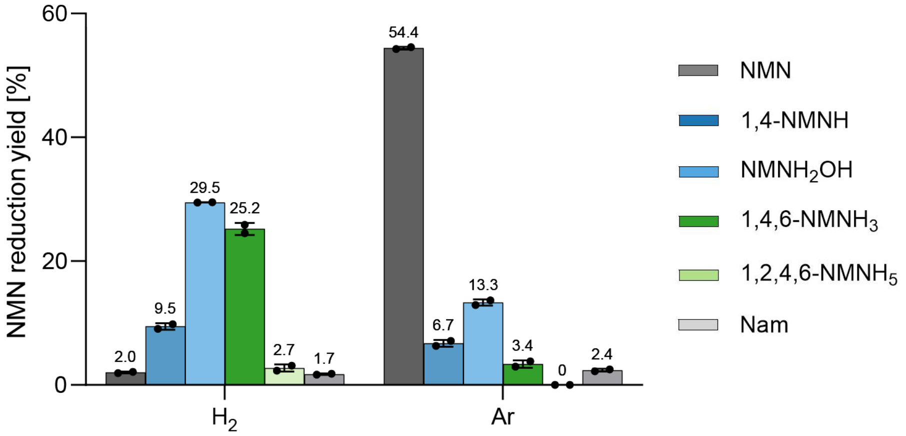

**SI Figure 29** Reduction of NMN in PBS (0.133 M. pH 8.5) after 4 h at 40 °C with equimolar amounts of $nNiFe_3$ (**SI Table 1**) and 5 bar of Ar or $H_2$. The supernatant was separated from the metal for qNMR (600 MHz) and the yields calculated as for **SI Table 8**, and **SI Table 9**. Each condition was tested with two replicas and a metal-free control.

### Overtime reduction of NAD/NMN with $H_2$ and $NiFe_3$

For quantification of each product, the ppm values of **SI Table 4**, and **SI Table 7** should be considered.

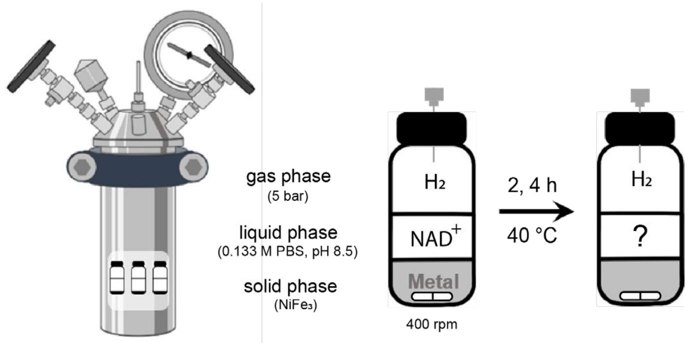

**SI Scheme 3** The reduction of $NAD^+$ with $nNiFe_3$ alloys was tested with the protocol described in detail in Methods, and according to the scheme above. The amount of metal and $NAD^+$ was 36 µmol, which reacted together for 2 and 4 h, at 40 °C, under alkaline conditions and 5 bars of $H_2$. The same reaction was made under Ar as a control.

**SI Table 10** After 2 and 4 h under 5 bar of $H_2$, as shown in **SI Scheme 3**, samples with $NiFe_3$ yielded different amounts of 1,4-NADH, 1,6-NADH, and Nam, from the starting material $NAD^+$. The starting metal and cofactor were 36 µmol mixed in 3 mL of 0.133 M PBS (pH 8.5). The amount of metal atoms was the same as the cofactor. The yields were calculated relative to the metal-free sample (100% $NAD^+$). To determine the TOF of each reaction, the sum of 1,4-NADH, and 1,6-NADH was considered as the amount of product, and the total amount of metal atoms was considered as the mount of catalyst, instead of the number of molecules. Experiments of 2h had duplicates while 4h long experiments had four replicas.

| | $H_2$ | **$NAD^+$** | **SD** | **1,4-NADH** | **SD** | **1,6-NADH** | **SD** | **Nam** | **SD** | **TOF [mol/s]** |
|---|---|---|---|---|---|---|---|---|---|---|
| 4h | $nNiFe_3$ | 25.59% | 4.4% | 40.60% | 1.8% | 16.70% | 0.6% | 4.48% | 0.1% | 3.02E-05 |
| 2h | | 68.38% | 2.5% | 15.05% | 1.3% | 6.61% | 0.6% | 7.41% | 0.4% | 3.18E-05 |

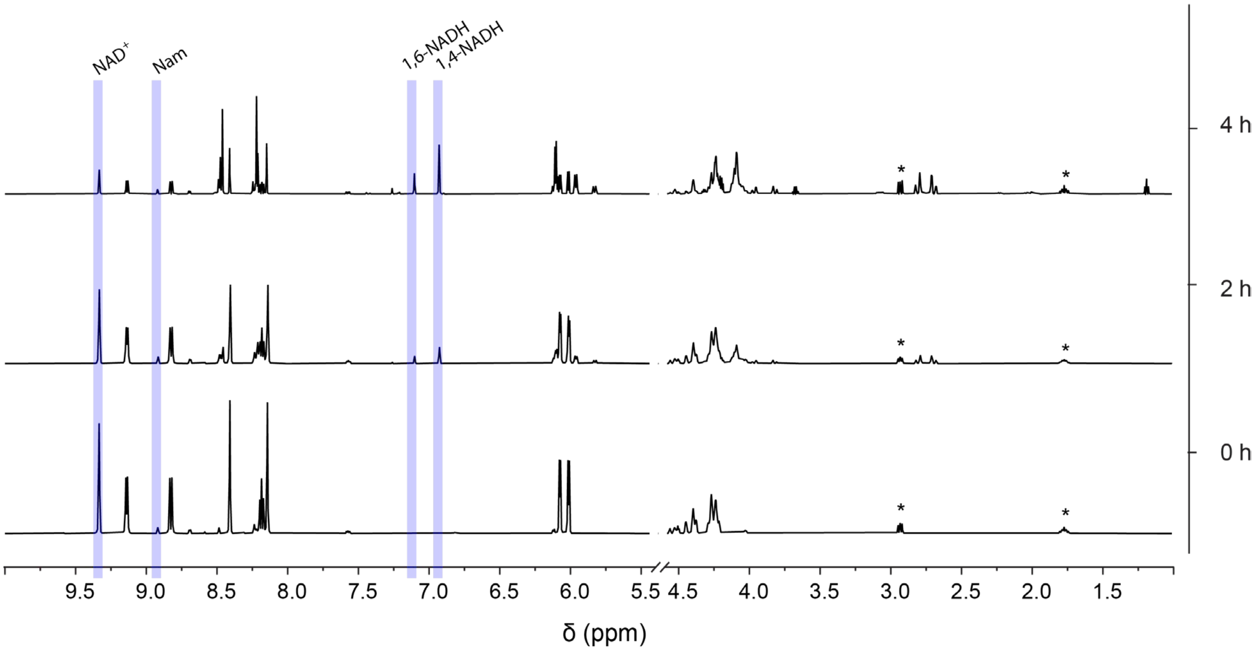

**SI Figure 30** The NMR spectra of replica samples of 36 µmol NAD+ in PBS (0.133 M, pH 8.5) with 5 bar of $H_2$ and 36 µmol $NiFe_3$ (1:1 cofactor ratio), as shown in **SI Scheme 3**, are stacked together in this figure. After 0, 2 and 4 h, the supernatant was collected and DSS added as an internal standard. The spectra were edited to only include relevant peaks, having been removed a DSS peak at 0 ppm and water peak at 4.8 ppm. No other peaks were found in the areas removed. Some DSS peaks are still visible (*). The peaks used for qualitative analysis and subsequent qNMR are highlighted in blue, according to **SI Table 4**.

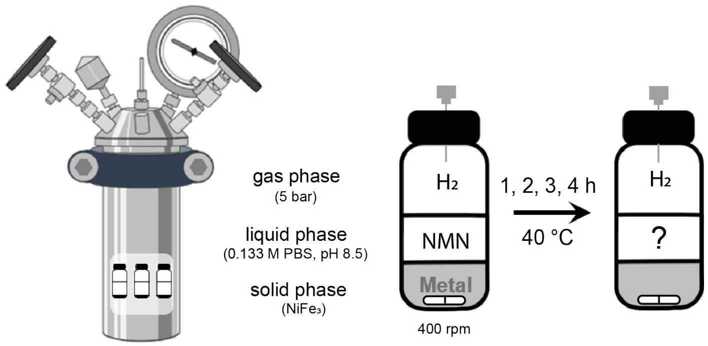

**SI Scheme 4** The reduction of NMN with $NiFe_3$ was tested with the protocol described in detail in Methods, and according to the scheme above. The amount of metal and NMN was 36 or 18 µmol, which reacted together for 1, 2, 3, and 4 h, at 40 °C, under alkaline conditions and 5 bars of $H_2$. The same reaction was made under Ar as a control.

**SI Table 11** After 1, 2, 3, and 4 h under 5 bar of $H_2$, as shown in **SI Scheme 4**, samples with $NiFe_3$ yielded different amounts of 1,4-NMNH, $NMNH_2OH$, 1,4,6-$NMNH_3$, 1,2,4,6-$NMNH_5$, and nicotinamide (Nam), from the starting material NMN. The starting metal and cofactor were 36 µmol mixed in 3 mL of 0.133 M PBS (pH 8.5). The amount of metal atoms was the same as the cofactor. The yields were calculated relative to the metal-free sample (100% NMN). To determine the TOF of each reaction, the sum of 1,4-NMNH, 1,4,6-$NMNH_3$, 1,2,4,6-$NMNH_5$ was considered as the amount of product, and the total amount of metal atoms was considered as the mount of catalyst, instead of the number of molecules. 2h and 4h were tested with duplicates while 1h and 3h with triplicates.

| | $H_2$ | NMN | SD | 1,4-NMNH | SD | 1,4,6-$NMNH_3$ | SD | $NMNH_2OH$ | SD | 1,2,4,6-$NMNH_5$ | SD | Nam | SD | TOF [mol/s] |
|---|---|---|---|---|---|---|---|---|---|---|---|---|---|---|
| 4h | $nNiFe_3$ | 2.01% | 0.1% | 9.46% | 0.4% | 25.22% | 0.7% | 29.49% | 0.0% | 2.73% | 0.4% | 1.75% | 0.1% | 1.12E-05 |
| 3h | | 1.21% | 0.6% | 24.74% | 0.0% | 19.72% | 2.6% | 21.32% | 1.4% | 2.82% | 0.6% | 2.05% | 0.2% | 7.10E-05 |
| 2h | | 9.38% | 3.5% | 34.03% | 0.7% | 14.48% | 1.4% | 16.64% | 0.3% | 1.78% | 0.4% | 2.54% | 0.2% | 7.55E-05 |
| 1h | | 30.80% | 16.7% | 29.84% | 7.1% | 7.40% | 2.7% | 12.67% | 2.7% | 0.73% | 0.5% | 2.87% | 0.3% | 1.14E-04 |

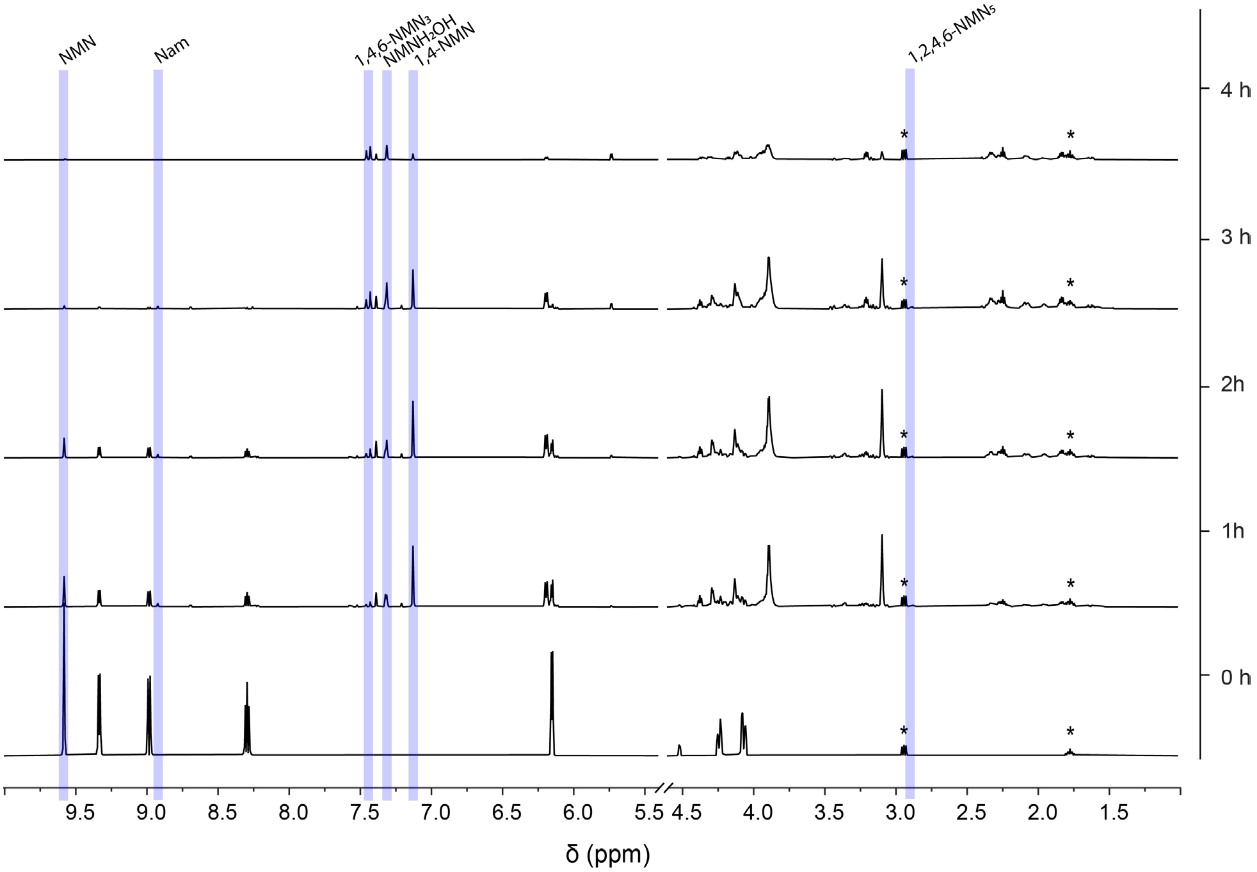

**SI Figure 31** The NMR spectra of replica samples of 36 or 18 µmol NMN in PBS (0.133 M, pH 8.5) with 5 bar of $H_2$ and equimolar amounts of $nNiFe_3$ (1:1 ratio), as shown in **SI Scheme 4**, are stacked together in this figure. After 0, 1, 2, 3 and 4 h, the supernatant was collected and DSS added as an internal standard. The spectra were edited to only include relevant peaks, having been removed a DSS peak at 0 ppm and water peak at 4.8 ppm. No other peaks were found in the areas removed. Some DSS peaks are still visible (*). The peaks used for qualitative analysis and subsequent qNMR are highlighted in blue, according to **SI Table 7**.

### Cyclic Voltametry of the reaction products

After a 4 h long reaction as described in **SI Scheme 3** and **SI Scheme 4**, the supernatant was diluted with PBS (0.133 M at pH 8.5) to obtain 15 mL of sample.

Maintaining the same experimental setup (oxygen-free 0.133 M PBS at pH 8.5), a three-electrode electrochemical cell was prepared for cyclic voltammetry with glassy carbon as working electrode, platinum wire as auxiliary electrode and $Ag/AgNO_3$ (0.01 M) as reference electrode. Samples were collected after 4h under the conditions described above, and diluted with buffer to reach a similar concentration to that of the standards (1 mM).

**SI Table 12** Table of oxidation potential

| Reagent | $E_a$ (V vs. SCE)[a] |
|---|---|
| 1,4-NADH | 0.531 |
| 1,4-NMNH | 0.609 |
| Reduced $NAD^+$ | 0.573 |
| Reduced NMN | 0.616; 1.000 |

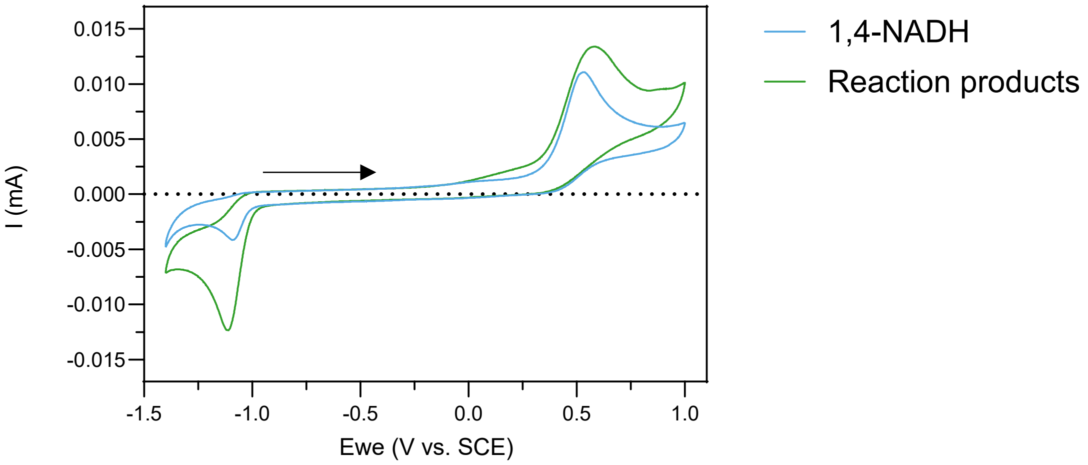

**SI Figure 32** Cyclic voltammogram of 1,4-NADH (1 mM in 0.133 M PBS pH 8.5; blue) and 12 mM of reduced $NAD^+$ with equimolar amounts of nNiFe (0.133 M PBS. 40 °C. 5 bar $H_2$.diluted 1:5 after reaction; green), measured at 25 ºC.

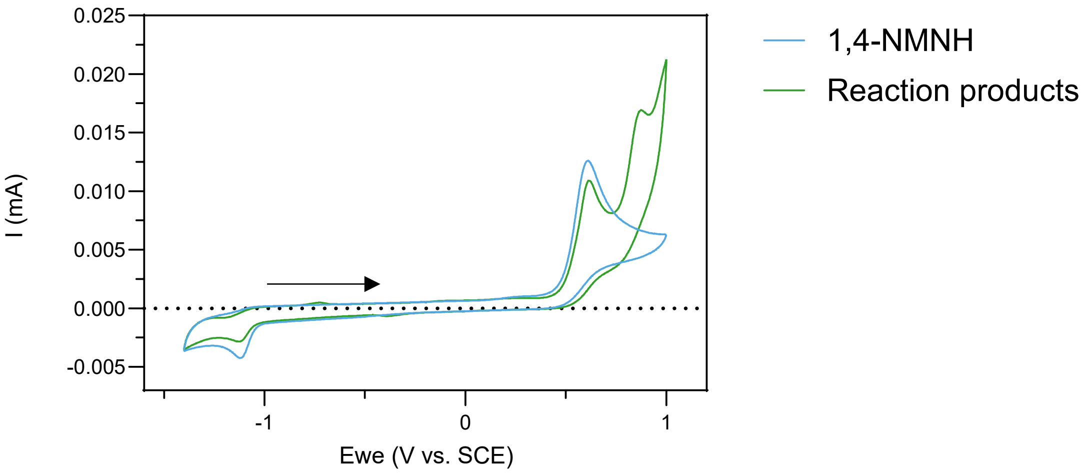

**SI Figure 33** Cyclic voltammogram of 1,4-NMNH (1 mM in 0.133 M PBS pH 8.5; blue) and 12 mM of reduced NMN with equimolar amounts of nNiFe (0.133 M PBS. 40 °C. 5 bar $H_2$.diluted with PBS 1:5 after reaction; green). measured at 25 ºC.

### Reduction of NAD/NMN with μNi$^0$ and μFe$^0$

In an attempt to further understand the differences observed in reactions with NMN and Ni/Fe alloys as catalysts, compared to NAD, experiments with the individual metals in micropowder form were performed.

For quantification of each product, the ppm values of **SI Table 4**, and **SI Table 7** should be considered.

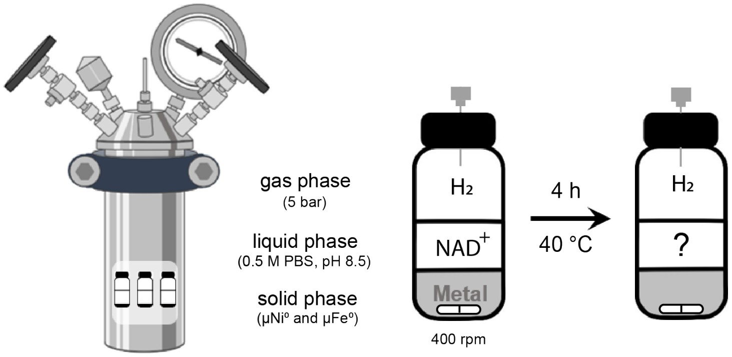

**SI Scheme 5** The reduction of NAD$^+$ with μFe$^0$ (particle size: <150 μm) and μNi$^0$ (particle size: 3–7 μm) was tested with the protocol described in detail in Methods, and according to the scheme above. The amount of NAD$^+$ was 36 μmol and metal was 1.8 mmol, which reacted together for 4 h, at 40 °C, under alkaline conditions and 5 bars of $H_2$.

**SI Table 13** After 4 h under 5 bar of $H_2$, as shown in **SI Scheme 5**, samples with μNi$^0$ or μFe$^0$ yielded different amounts of 1,4-NADH, 1,6-NADH, and Nam, from the starting material NAD$^+$. The starting metal and cofactor were 36 μmol mixed in 3 mL of 0.5 M PBS (pH 8.5). The amount of metal atoms was 50 times the moles of cofactor. The yields were calculated relative to the metal-free sample (100% NAD$^+$). To determine the TOF of each reaction, the sum of 1,4-NADH, and 1,6-NADH was considered as the amount of product. All conditions had duplicates and a metal-free control. In comparison to a former study with μNi[7] (333:1), where no 1,4-NADH was formed unless the powder was pretreated with $H_2$ before the reaction, the same powder produces a 13.6% yield of 1,4-NADH in the experiment described here (50:1). We attribute this different outcome to the degassing of this experiments' buffer opposed to no degassing in Pereira et al. 2022.[7] Degassing does not seem to have the same consequences to μFe, most likely due to the different reduction mechanism at place (Fe donating electrons while Ni depending on $H_2$ activation on its surface).

| | $H_2$ | NAD$^+$ | SD | 1,4-NADH | SD | 1,6-NADH | SD | Nam | SD | TOF [mol/s] |
|---|---|---|---|---|---|---|---|---|---|---|
| 4h | μNi$^0$ | 55.22% | 1.6% | 13.56% | 0.9% | 1.69% | 0.6% | 10.82% | 0.5% | 2.24E-07 |
| | μFe$^0$ | 52.07% | 4.3% | 5.75% | 0.1% | 0.30% | 0.3% | 31.24% | 3.4% | 8.88E-08 |

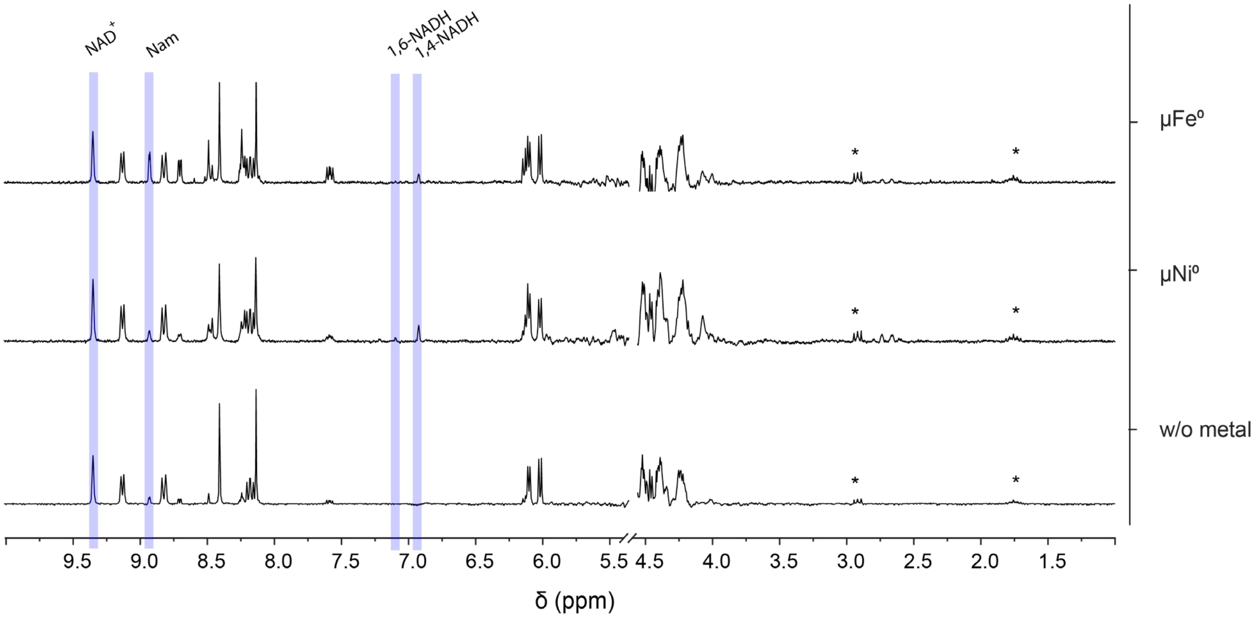

**SI Figure 34** The NMR spectra of replica samples of 36 μmol $NAD^+$ in PBS (0.5 M, pH 8.5) with 5 bar of $H_2$ and 1.8 mmol $\mu Fe^0$, $\mu Ni^0$ (50:1 cofactor ratio), or no metal, as shown in **SI Scheme 5**, are stacked together in this figure. After the 4h reaction, the supernatant was collected and DSS added as an internal standard. The spectra were edited to only include relevant peaks, having been removed a DSS peak at 0 ppm and water peak at 4.8 ppm. No other peaks were found in the areas removed. Some DSS peaks are still visible (*). The peaks used for qualitative analysis and subsequent qNMR are highlighted in blue, according to **SI Table 4**.

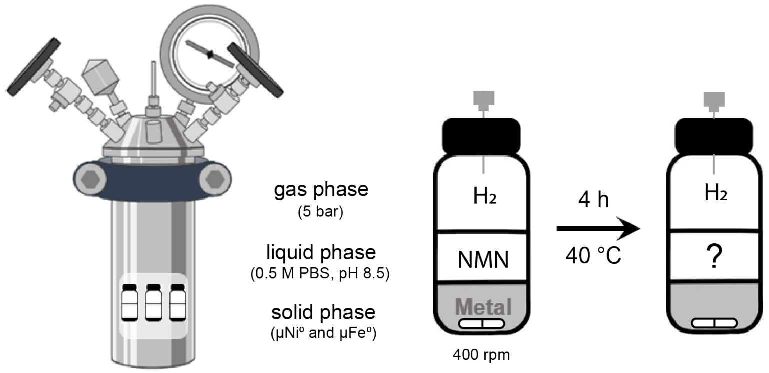

**SI Scheme 6** The reduction of NMN with $\mu Fe^0$ and μNi was tested with the protocol described in detail in Methods, and according to the scheme above. The amount of NMN was 36 μmol and metal was 1.8 mmol, which reacted together for 4 h, at 40 °C, under alkaline conditions and 5 bars of $H_2$. The same reaction was made under Ar as a control.

**SI Table 14** After 4 h under 5 bar of $H_2$, as shown in **SI Scheme 6**, samples with μ$Ni^0$ or μ$Fe^0$ yielded different amounts of 1,4-NMNH, 1,4,6-$NMNH_3$, $NMNH_2OH$, and nicotinamide (Nam), from the starting material NMN. The starting metal and cofactor were 1.8 mmol and 36 μmol, respectively, mixed in 3 mL of 0.5 M PBS (pH 8.5). The amount of metal atoms was fifty times of the cofactor. The yields were calculated relative to the metal-free sample (100% NMN). To determine the TOF of each reaction, the sum of 1,4-NMNH, 1,2,4,6-$NMNH_5$, and 1,4,6-$NMNH_3$ was considered as the amount of product. All conditions were tested in triplicate.

| | $H_2$ | NMN | SD | 1,4-NMNH | SD | 1,4,6-$NMNH_3$ | SD | $NMNH_2OH$ | SD | 1,2,4,6-$NMNH_5$ | SD | Nam | SD | TOF [mol/s] |
|---|---|---|---|---|---|---|---|---|---|---|---|---|---|---|
| 4h | μ$Ni^0$ | 6.27% | 4.5% | 7.06% | 1.6% | 23.76% | 7.0% | 18.03% | 2.6% | 0.00% | 0.0% | 1.46% | 2.1% | 4.63E-07 |
| | μ$Fe^0$ | 41.28% | 3.3% | 25.09% | 3.2% | 0.00% | 0.0% | 7.04% | 1.8% | 0.00% | 0.0% | 18.73% | 2.2% | 3.77E-07 |

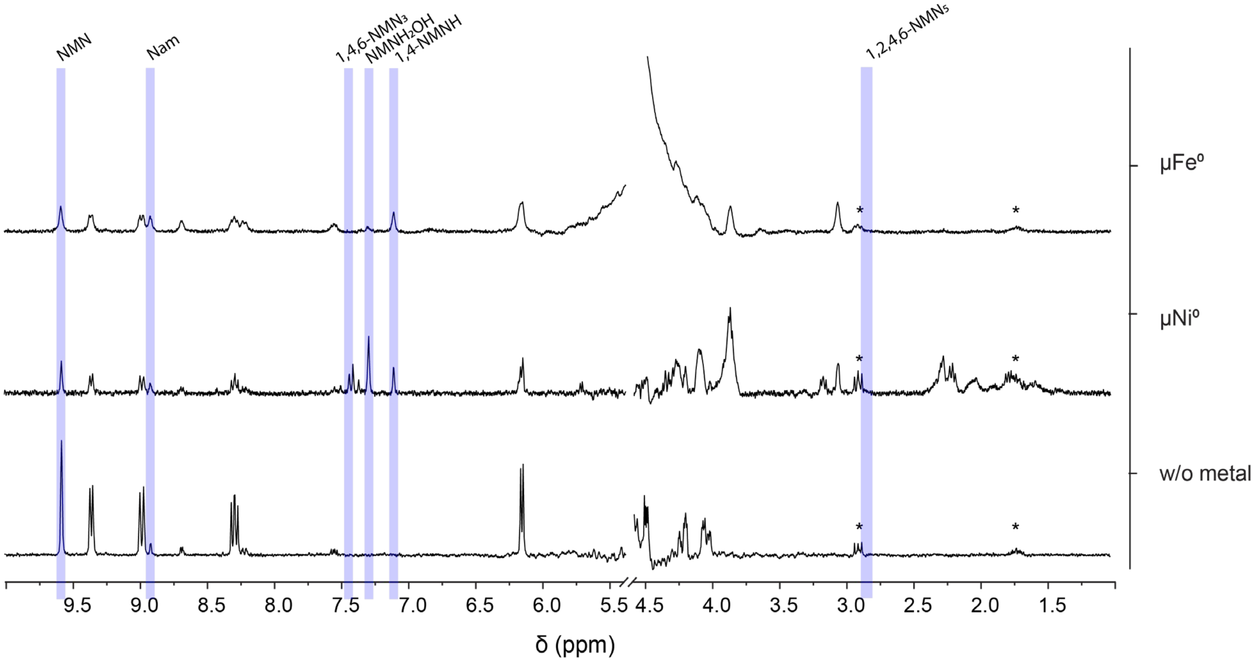

**SI Figure 35** The NMR spectra of replica samples of 36 μmol NMN in PBS (0.5 M, pH 8.5) with 5 bar of $H_2$ and 1.8 mmol μ$Fe^0$, μ$Ni^0$ (50:1 cofactor ratio), or no metal, as shown in **SI Scheme 6**, are stacked together in this figure. After the 4h reaction, the supernatant was collected and DSS added as an internal standard. The spectra were edited to only include relevant peaks, having been removed a DSS peak at 0 ppm and water peak at 4.8 ppm. No other peaks were found in the areas removed. Some DSS peaks are still visible (*). The peaks used for qualitative analysis and subsequent qNMR are highlighted in blue, according to **SI Table 7**.

**SI Table 15** After 4 h under 5 bar of Ar, as shown in **SI Scheme 6**, samples with μ$Ni^0$ or μ$Fe^0$ yielded different amounts of 1,4-NMNH, $NMNH_2OH$, and nicotinamide (Nam), from the starting material NMN. The starting metal and cofactor were 1.8 mmol and 36 μmol, respectively, mixed in 3 mL of 0.5 M PBS (pH 8.5). The amount of metal atoms was fifty times of the cofactor. The yields were calculated relative to the metal-free sample (100% NMN). To determine the TOF of each reaction, the sum of 1,4-NMNH, 1,4,6-$NMNH_3$, and 1,2,4,6-$NMNH_5$ was considered as the amount of product. All experiments were performed in duplicate.

| | Ar | NMN | SD | 1,4-NMNH | SD | 1,4,6-$NMNH_3$ | SD | $NMNH_2OH$ | SD | 1,2,4,6-$NMNH_5$ | SD | Nam | SD | TOF [mol/s] |
|---|---|---|---|---|---|---|---|---|---|---|---|---|---|---|
| 4h | μ$Ni^0$ | 86.34% | 3.1% | 0.00% | 0.0% | 0.00% | 0.0% | 0.00% | 0.0% | 0.00% | 0.0% | 5.40% | 5.4% | 0.00E+00 |
| | μ$Fe^0$ | 45.66% | 2.4% | 22.21% | 1.2% | 0.00% | 0.0% | 5.44% | 0.2% | 0.00% | 0.0% | 22.25% | 0.2% | 3.34E-07 |

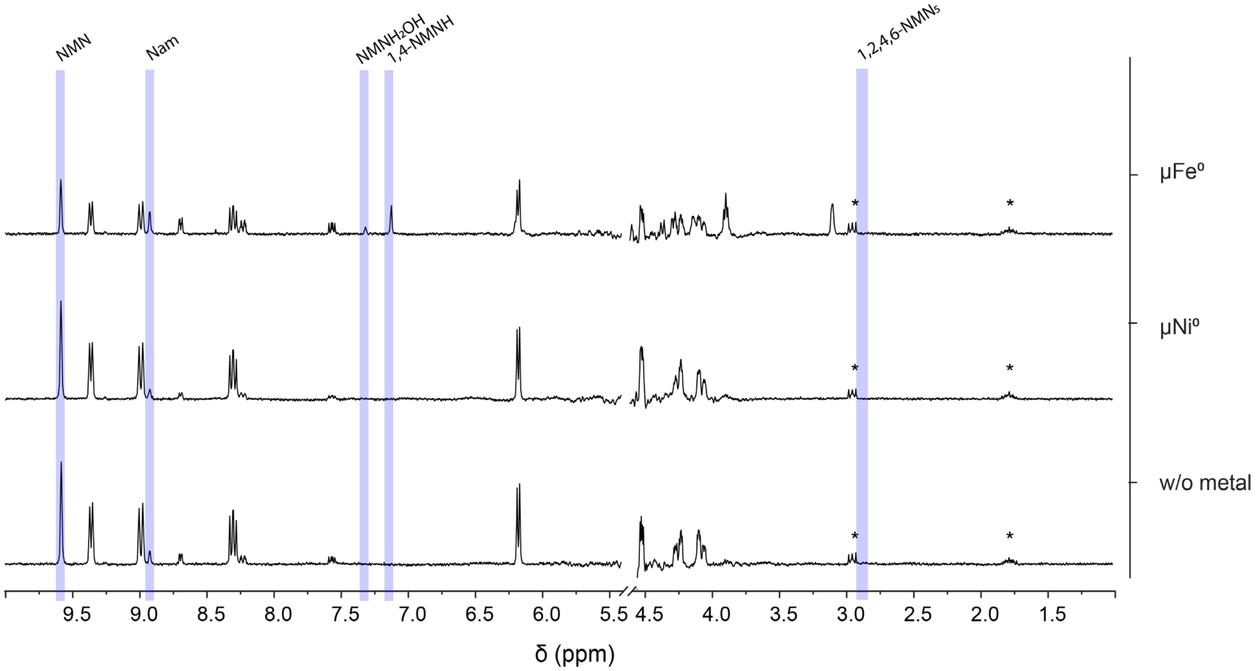

**SI Figure 36** The NMR spectra of replica samples of 36 μmol NMN in PBS (0.5 M, pH 8.5) with 5 bar of Ar and 1.8 mmol μ$Fe^0$, μ$Ni^0$ (50:1 cofactor ratio), or no metal, as shown in **SI Scheme 6**, are stacked together in this figure. After the 4h reaction, the supernatant was collected and DSS added as an internal standard. The spectra were edited to only include relevant peaks, having been removed a DSS peak at 0 ppm and water peak at 4.8 ppm. No other peaks were found in the areas removed. Some DSS peaks are still visible (*). The peaks used for qualitative analysis and subsequent qNMR are highlighted in blue, according to **SI Table 7**.

In the reactions with micropowder (**SI Scheme 6**), the product 1,2,4,6-$NMNH_5$ was never detected with any metal. Considering that the surface area of x amount of micropowder is considerably smaller than that of the same amount of nanopowders, it is possible that the amounts of micropowder used was insufficient to mimic the reaction conditions of nanopowder metals. Additionally, due to the difference in the NMR spectrometer used between samples with nano- (600 mHz) and micropowder (300 mHz), it is possible that higher yields are necessary to detect that product. Therefore, experiments with the same conditions described in **SI Scheme 6**, but with a metal to cofactor ratio of 200:1 instead of 50:1 were made. 7

**SI Table 16** After 4 h under 5 bar of $H_2$, as shown in **SI Scheme 6**, samples with $\mu Ni^0$ or $\mu Fe^0$ yielded different amounts of 1,4-NMNH, 1,4,6-$NMNH_3$, 1,2,4,6-$NMNH_5$, $NMNH_2OH$, and nicotinamide (Nam), from the starting material NMN. The starting metal and cofactor were 3.2 mmol and 18 µmol, respectively, mixed in 3 mL of 0.5 M PBS (pH 8.5). The amount of metal atoms was two hundred times of the cofactor. The yields were calculated relative to the metal-free sample (100% NMN). To determine the TOF of each reaction, the sum of 1,4-NMNH, 1,4,6-$NMNH_3$, and 1,2,4,6-$NMNH_5$ was considered as the amount of product. Fe-containing experiments were performed in duplicate and Ni-containing in triplicate.

| | $H_2$ | NMN | SD | 1,4-NMNH | SD | 1,4,6-$NMNH_3$ | SD | $NMNH_2OH$ | SD | 1,2,4,6-$NMNH_5$ | SD | Nam | SD | TOF [mol/s] |
|---|---|---|---|---|---|---|---|---|---|---|---|---|---|---|
| 4h | $\mu Ni^0$ | 0,70% | 0,5% | 1,04% | 1,2% | 37,05% | 3,0% | 4,02% | 3,1% | 11,49% | 3,1% | 0,21% | 0,3% | 1,5082E-07 |
| | $\mu Fe^0$ | 20,07% | 17,8% | 44,73% | 20,3% | 0,23% | 0,0% | 9,53% | 4,2% | 0,81% | 0,3% | 25,43% | 5,5% | 1,3925E-07 |

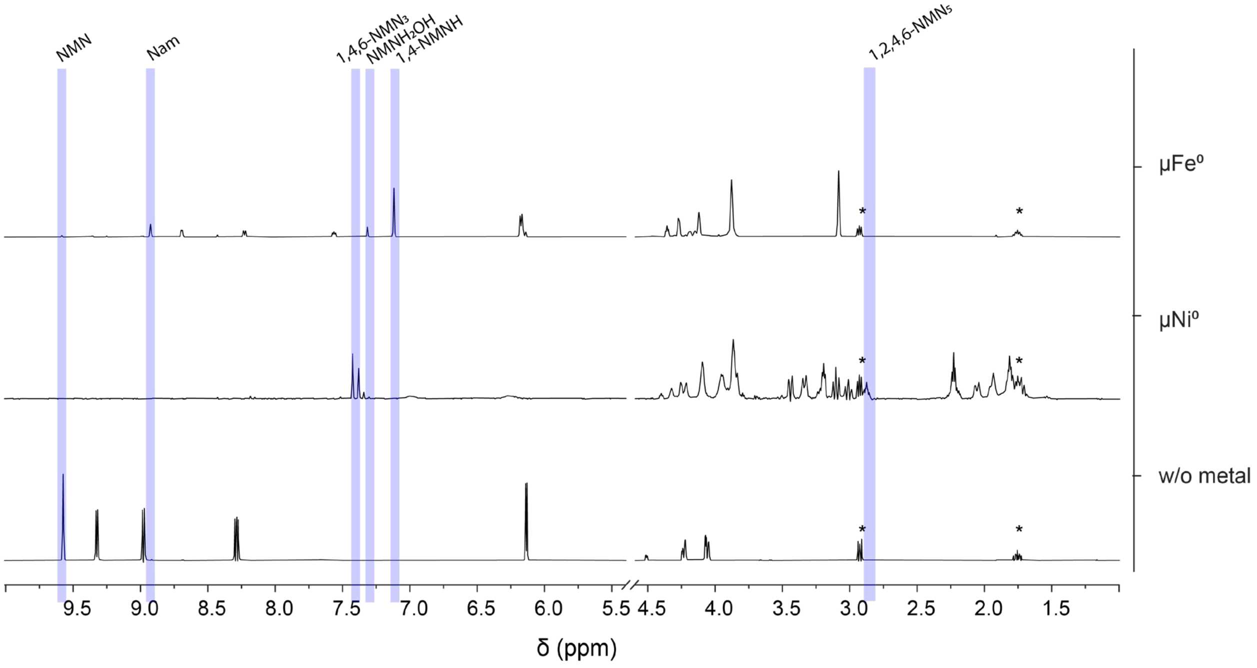

**SI Figure 37** The NMR spectra of replica samples of 18 µmol NMN in PBS (0.5 M, pH 8.5) with 5 bar of $H_2$ and 3.2 mmol $\mu Fe^0$, $\mu Ni^0$ (200:1 cofactor ratio), or no metal, as shown in **SI Scheme 6**, are stacked together in this figure. After the 4h reaction, the supernatant was collected and DSS added as an internal standard. The spectra were edited to only include relevant peaks, having been removed a DSS peak at 0 ppm and water peak at 4.8 ppm. No other peaks were found in the areas removed. Some DSS peaks are still visible (*). The peaks used for qualitative analysis and subsequent qNMR are highlighted in blue, according to **SI Table 7**.

In this experiment, nickel yielded significant amounts of 1,2,4,6-$NMNH_5$, detected at 2.85 ppm as observed in **SI Figure 37**.

#### Specificity of each metal to 1,4-NADH production

In the experiments described with NAD, the only other reduction product made other than 1,4-NADH, in significant amounts and identified through 2D-NMR, was 1,6-NADH. The amount of 1,6-NADH relative to the total amount of NADH detected in each sample was calculated and presented in the table below.

**SI Table 17** After 4 h under 5 bar of $H_2$, as shown in **SI Scheme 1**, samples with different Ni/Fe alloys yielded different amounts of 1,4-NADH and 1,6-NADH, from the starting material $NAD^+$. The starting amount of cofactor was 36 µmol mixed in 3 mL of PBS (pH 8.5), with equimolar amounts of metal nanopowder (0.133 M PBS buffer) or 50 times more if using micropowder (0.5 M PBS buffer). The yields were calculated relative to the metal-free sample (100% $NAD^+$) as in **SI Table 5** and **SI Table 13**. To determine the specificity of each reaction to reduce NAD in the $4^{th}$ of $6^{th}$ carbon of the nicotinamide moiety, the amount of 1,6-NADH was normalized to the total amount of NADH in the sample (1,4-NADH + 1,6-NADH). Reactions with nNi, $nNi_3Fe$, and nNiFe were performed with six replicas; $nFe^0$ and w/o metal were done with five replicas; $nNiFe_3$ amounted to four; $nNiFe_3$, $µNi^0$, and $µFe^0$ had duplicates. Calculating mere surface area differences between nNi and µNi powder results in a ratio of roughly 50:1, for nFe and µFe the ratio is 1:1400.

| $H_2$ | 1,6-NADH/NADH | SD |
|---|---|---|
| $µNi^0$ | 11,16% | 3,91% |
| $nNi^0$ | 24,34% | 1,36% |
| $nNi_3Fe$ | 32,63% | 0,82% |
| nNiFe | 28,58% | 1,19% |
| $nNiFe_3$ | 29,15% | 0,20% |
| $nFe^0$ | 0,00% | 0,00% |
| $µFe^0$ | 4,67% | 4,67% |

According to **SI Table 17**, micropowder metals tend to lead to the accumulation of less 1,6-NADH compared to nanopowders. 1,6-NADH was not detected in samples with $nFe^0$, most likely due to the low NADH yield and high amounts of dissolved paramagnetic metal ions leading to line broadening. Contrastingly, $nNiFe_3$ is the only metal to produce 1,6-NADH under Ar, due to its comparably high TOF (s. SI Tab.9). The ratio of 1,6-NADH to NADH is maintained independently of the gas phase and yield at approximately 30%.

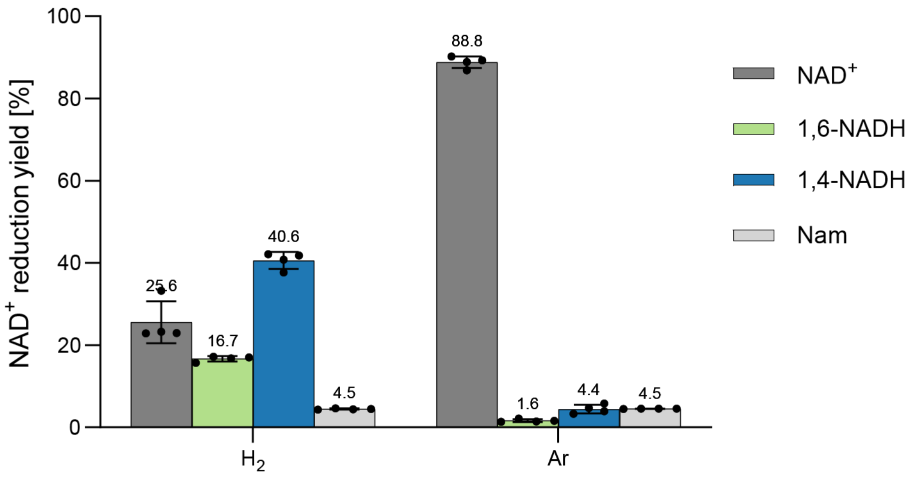

**SI Figure 38** 36 µmol of $NAD^+$ reacted in PBS (0.133 M, pH 8.5) for 4 h at 40 °C with equimolar amounts of $nNiFe_3$ and 5 bar of $H_2$ or Ar. The supernatant was separated from the metal for qNMR (600 MHz) and the yields calculated according to **SI Table 5** and **SI Table 6**. Each condition was tested with four replicas and a metal-free control.

To further understand the accumulation of 1,6-NADH, the reaction above was reproduced under an inert $N_2$ atmosphere, and starting with a solution of 1,4-NADH instead of $NAD^+$ and with duplicates.

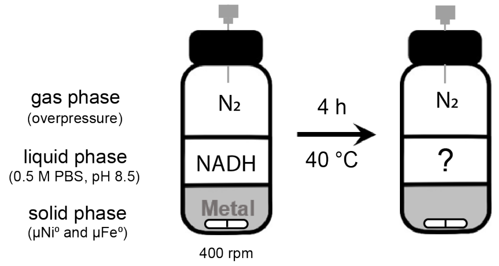

**SI Scheme 7** The conversion of 1,4-NADH to 1,6-NADH was tested according to the scheme above. The amount of 1,4-NADH was 36 µmol, and 50 times more metal was added to each vial ($\mu Fe^0$ and $\mu Ni^0$) which reacted together for 4 h, at 40 °C, under alkaline conditions and a $N_2$ overpressure.

**SI Table 18** After 4 h under an overpressure of $N_2$, as shown in **SI Scheme 7**, samples with µNi and µFe yielded different amounts of 1,4-NADH and 1,6-NADH, from the starting material 1,4-NADH. The starting cofactor was 36 µmol mixed in 3 mL of 0.5 M PBS (pH 8.5) and with 1.8 mmol of µFe or µNi. To determine the specificity of each reaction to reduce NAD in the 4$^{th}$ of 6$^{th}$ carbon of the nicotinamide moiety, the amount of 1,6-NADH was normalized to the total amount of NADH in the sample (1,4-NADH + 1,6-NADH). All conditions had duplicates.

| $N_2$ | 1,6-NADH/NADH | SD |
|---|---|---|
| control | 15.3% | 0,00% |
| $\mu Fe^0$ | 10,11% | 0,78% |
| $\mu Ni^0$ | 12,91% | 0,33% |

Starting from a solution of only 1,4-NADH, all samples yielded more 1,6-NADH than when starting from $NAD^+$ (**SI Table 17**). Additionally, the metal-containing experiments did not accumulate more 1,6-NADH than the control. Given the differences observed between metals in **SI Table 17**, it is clear that metals can influence the production of 1,6-NADH, but it also seems that in the case of micropowders, it mostly results from an equilibrium reaction of NADH in solution.

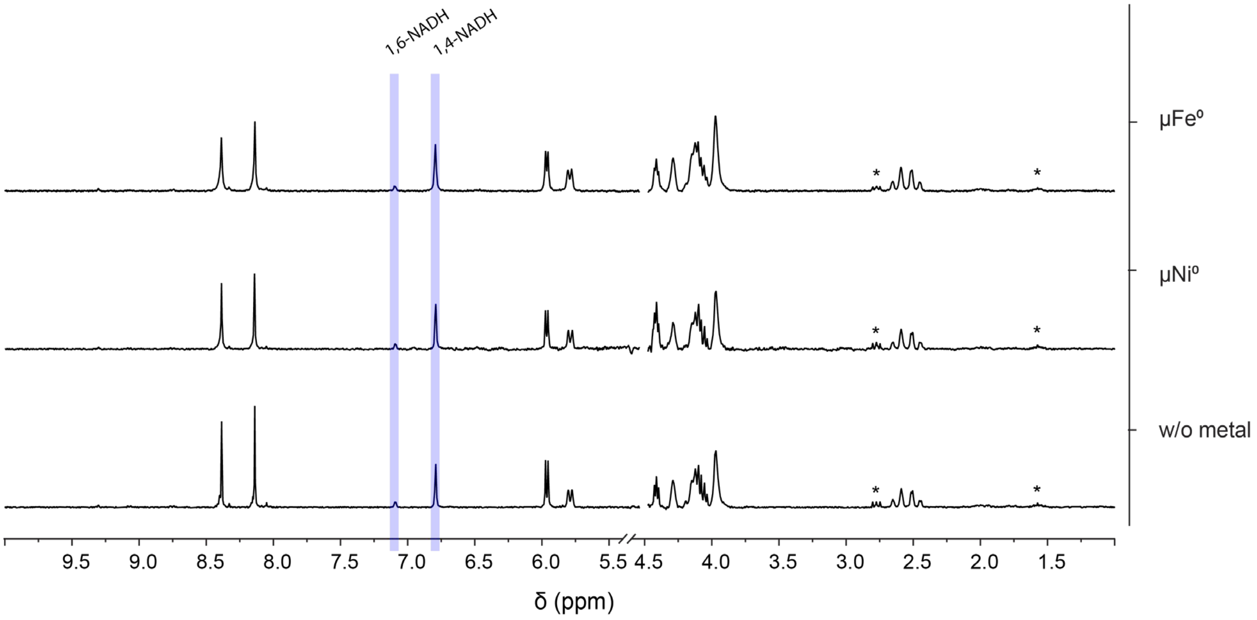

**SI Figure 39** The NMR spectra of replica samples of 1,4-NADH in PBS (0.5 M, pH 8.5), as shown in **SI Scheme 7**, are stacked together in this figure. The metal used in each sample is the micropowder indicated on the right. After 4h reacting, the supernatant was collected and DSS added as an internal standard. The spectra were edited to only include relevant peaks, having been removed a DSS peak at 0 ppm and water peak at 4.8 ppm. No other peaks were found in the areas removed. Some DSS peaks are still visible (*). The peaks used for qualitative analysis and subsequent qNMR are highlighted in blue, according to **SI Table 4**.

### "Competition" experiments

To further test the reactivity of NMN and NAD under prebiotic conditions, experiments with both cofactors in the same reaction mixture were designed.

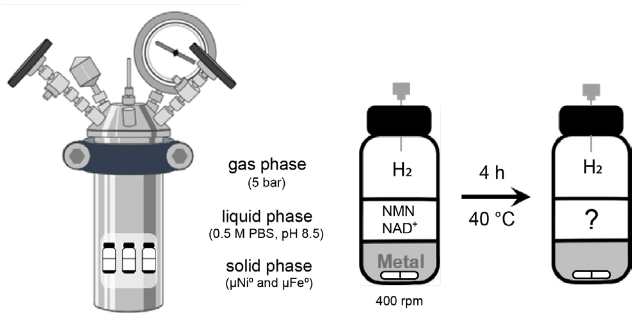

**SI Scheme 8** The reduction of a mixture of NMN and $NAD^+$ with $\mu Fe^0$ or $\mu Ni^0$ was tested with the general protocol described in detail in Methods, and according to the scheme above. The amount of cofactor was 36 or 72 μmol (18 or 36 each, respectively) in 3 mL of pH 8.5 0.5 M PBS buffer, the metal amount was constantly 1.8 mmol (leading to a metal-cofactor ratio of 50:1 and 25:1, respectively). The reaction took place for 4 h at 40 °C under slightly alkaline conditions and 5 bars of $H_2$.

**SI Table 19** After 4 h under 5 bar of $H_2$, as shown in **SI Scheme 8**, samples with $\mu Fe^0$ or $\mu Ni^0$ yielded different amounts of 1,4-NADH, 1,6-NADH, 1,4-NMNH, $NMNH_2OH$, 1,4,6-$NMNH_3$, and nicotinamide (Nam), from the starting material NMN and $NAD^+$. The starting cofactor mixture was 36 μmol (18 μmol each nucleotide) mixed in 3 mL of 0.5 M PBS (pH 8.5), with a metal-cofactor ratio of 50:1. The yields were calculated separately and relative to the metal-free sample (100% NMN and 100% $NAD^+$). To determine the TOF of each reaction, the sum of 1,4-NMNH, 1,4,6-$NMNH_3$, 1,4-NADH, and 1,6-NADH was considered as part of the reduction products. The NMN and NAD products are presented separately in order for the table to fit the page, but belong to the same set of triplicates.

| | $H_2$ | **1,4-NMNH** | **SD** | **1,4,6-$NMNH_3$** | **SD** | **$NMNH_2OH$** | **SD** | **1,2,4,6-$NMNH_5$** | **SD** | **NMN** | **SD** |
|---|---|---|---|---|---|---|---|---|---|---|---|
| 4h | $\mu Ni^0$ | 12.52% | 0.9% | 2.35% | 0.1% | 8.52% | 0.5% | 0.00% | 0.0% | 20.02% | 1.4% |
| | | **1,4-NADH** | **SD** | **1,6-NADH** | **SD** | **$NAD^+$** | **SD** | **TOF [mol/s]** | | | |
| | | 17.85% | 1.2% | 7.15% | 0.5% | 10.36% | 1.4% | 5.54E-07 | | | |
| | $H_2$ | **1,4-NMNH** | **SD** | **1,4,6-$NMNH_3$** | **SD** | **$NMNH_2OH$** | **SD** | **1,2,4,6-$NMNH_5$** | **SD** | **NMN** | **SD** |
| | $\mu Fe^0$ | 3.17% | 0.2% | 0.00% | 0.0% | 0.73% | 0.1% | 0.00% | 0.0% | 34.71% | 3.2% |
| | | **1,4-NADH** | **SD** | **1,6-NADH** | **SD** | **$NAD^+$** | **SD** | **TOF [mol/s]** | | | |
| | | 6.15% | 0.1% | 1.16% | 0.0% | 22.68% | 3.1% | 1.54E-07 | | | |

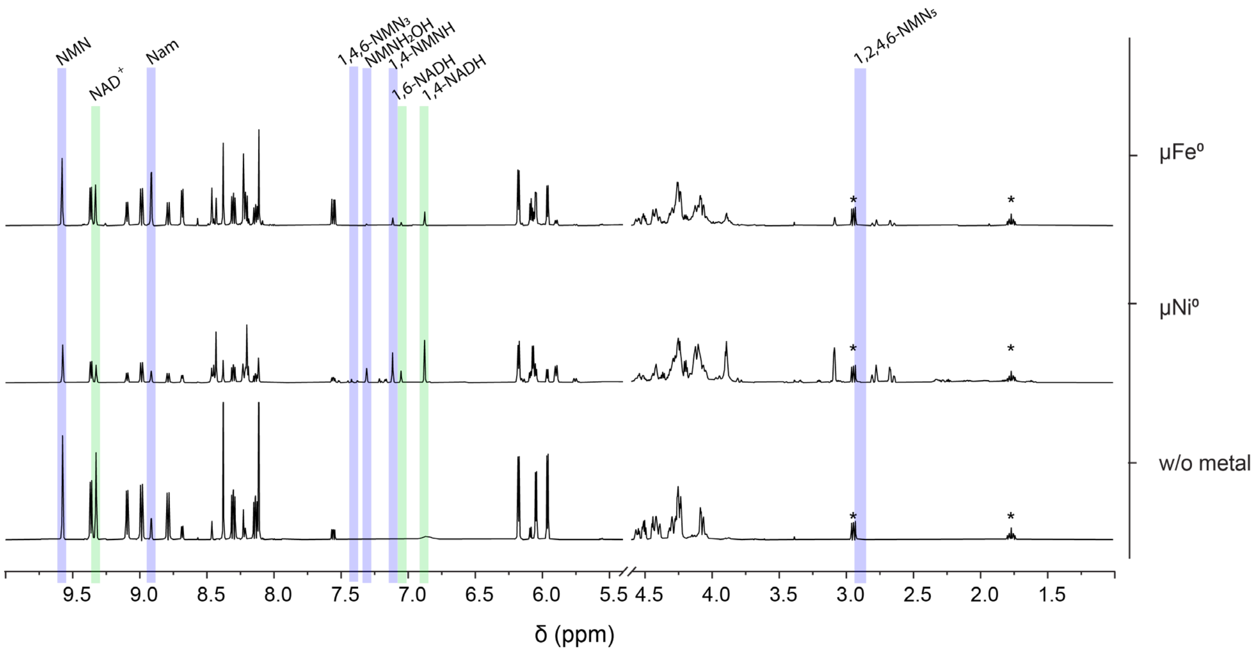

**SI Figure 40** The NMR spectra of replica samples with a mixture of 18 μmol NMN and 18 μmol $NAD^+$ in PBS (0.5 M, pH 8.5) with 5 bar of $H_2$ and 1.8 mmol $\mu Fe^0$, $\mu Ni^0$ (50:1 cofactor ratio), or no metal, as shown in **SI Scheme 8**, are stacked together in this figure. After the 4h reaction, the supernatant was collected and DSS added as an internal standard. The spectra were edited to only include relevant peaks, a DSS peak at 0 ppm and the water peak at 4.8 ppm have been removed. No other peaks were found in the areas removed. Other DSS peaks are still visible (*). The peaks used for qualitative analysis and subsequent qNMR are highlighted in blue (NMN products) and green (NAD products), according to **SI Table 7**.

**SI Table 20** After 4 h under 5 bar of $H_2$, as shown in **SI Scheme 8**, samples with $\mu Fe^0$ or $\mu Ni^0$ yielded different amounts of 1,4-NADH, 1,4-NMNH, $NMNH_2OH$, and nicotinamide (Nam), from the starting material NMN and $NAD^+$. The starting cofactor mixture was 72 μmol (36 μmol each nucleotide) mixed in 3 mL of 0.5 M PBS (pH 8.5), with a metal-cofactor ratio of 25:1. The yields were calculated separately and relative to the metal-free sample (100% NMN and 100% $NAD^+$). To determine the TOF of each reaction, the sum of 1,4-NMNH, 1,4,6-$NMNH_3$, 1,2,4,6-$NMNH_5$, 1,4-NADH, and 1,6-NADH was considered as part of the reduction products. The NMN and NAD products are presented separately in order for the table to fit the page, but belong to the same set of duplicates.

| | $H_2$ | **1,4-NMNH** | **SD** | **1,4,6-$NMNH_3$** | **SD** | **$NMNH_2OH$** | **SD** | **1,2,4,6-$NMNH_5$** | **SD** | **NMN** | **SD** |
|---|---|---|---|---|---|---|---|---|---|---|---|
| 4h | $\mu Ni^0$ | 3.83% | 0.4% | 0.00% | 0.0% | 4.34% | 0.5% | 0.00% | 0.0% | 66.81% | 1.1% |
| | | **1,4-NADH** | **SD** | **1,6-NADH** | **SD** | **$NAD^+$** | **SD** | **TOF [mol/s]** | | | |
| | | 8.62% | 0.3% | 0.00% | 0.0% | 50.02% | 1.9% | 5.61E-08 | | | |
| | $H_2$ | **1,4-NMNH** | **SD** | **1,4,6-$NMNH_3$** | **SD** | **$NMNH_2OH$** | **SD** | **1,2,4,6-$NMNH_5$** | **SD** | **NMN** | **SD** |
| | $\mu Fe^0$ | 4.52% | 0.4% | 0.00% | 0.0% | 0.00% | 0.0% | 0.00% | 0.0% | 68.06% | 7.6% |
| | | **1,4-NADH** | **SD** | **1,6-NADH** | **SD** | **$NAD^+$** | **SD** | **TOF [mol/s]** | | | |
| | | 8.16% | 1.3% | 1.64% | 0.1% | 49.78% | 2.5% | 6.63E-08 | | | |

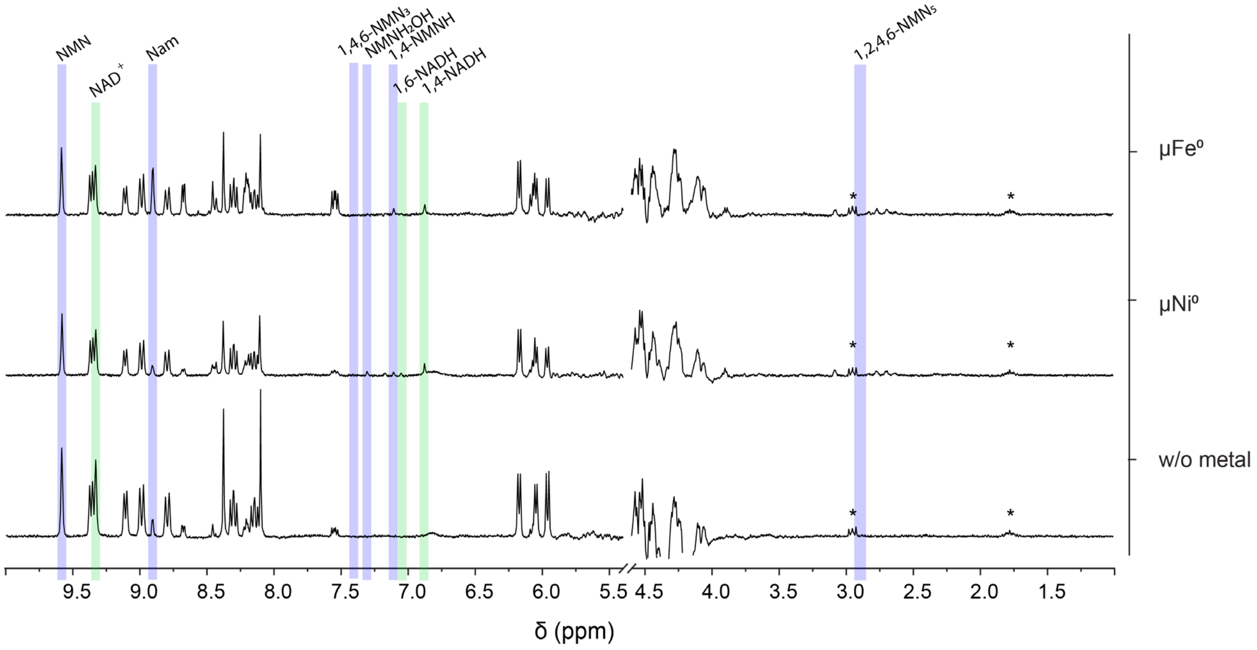

**SI Figure 41** The NMR spectra of replica samples with a mixture of 36 μmol NMN and 36 μmol $NAD^+$ in PBS (0.5 M, pH 8.5) with 5 bar of $H_2$ and 1.8 mmol $\mu Fe^0$, $\mu Ni^0$ (25:1 metal-cofactor ratio), or no metal, as shown in **SI Scheme 8**, are stacked together in this figure. After the 4h reaction, the supernatant was collected and DSS added as an internal standard. The spectra were edited to only include relevant peaks, a DSS peak at 0 ppm and the water peak at 4.8 ppm have been removed. No other peaks were found in the areas removed. Other DSS peaks are still visible (*). The peaks used for qualitative analysis and subsequent qNMR are highlighted in blue (NMN products) and green (NAD products), according to **SI Table 4** and **SI Table 7**.

### Reduction products of NMN and NAD characterized through NMR spectroscopy

**SI Figure 42** shows the $^1$H spectrum of substrate **1** in 0.133 M PBS at 298 K. Characteristic signals at 9.581, 9.337, 8.990, 8.308, 6.187, 4.472 ppm were observed for H-2, H-6, H-4, H-5, H-1'and H-4' respectively. Furthermore, two double doublets at 4.197 and 4.025 ppm were detected for the diastereotopic methylene $CH_2$-5'.

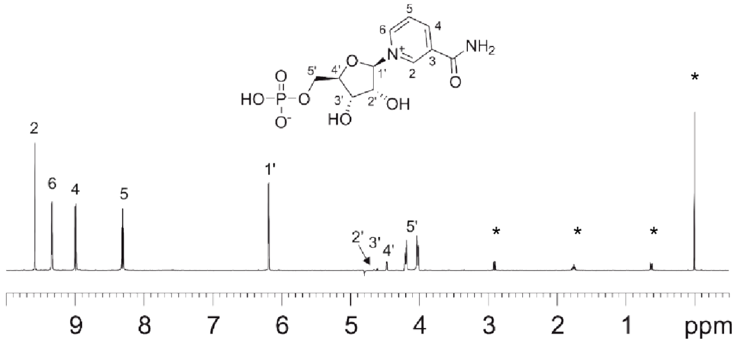

**SI Figure 42** $^1$H spectrum of substrate $NMN^+$ (**1**) in 0.133 M PBS with the internal reference DSS (*) at 298 K.

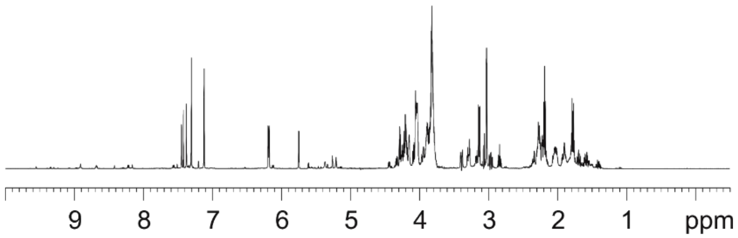

**SI Figure 43** $^1$H spectrum of the products NMNH in 0.133 M PBS at 298 K.

#### Identification of products of substrate 1

**SI Figure 43** presents $^1$H spectrum of products solution of substrate **1** after 4 h of incubation. This spectrum shows a strong reduction in the intensity of signals from 8.3 to 9.6 ppm. which means a large degradation of the substrate. A large number of signals from 1.3 to 7.5 ppm were observed, which reveals the formation of a group of more than three different species of products. As shown in **SI Scheme 9**, three single reduction products, the 1.2-NMNH. 1,4-NMNH and 1,6-NMNH may form. Further reduction of the nicotineamide may follow, thus 1,2,4-, 1,2,5-, 1,2,6-, 1,4,6-, and 1,2,4,6-$NMNH_x$ may form, too.

Due to the large number of species in the reaction solution, efforts to separate the products through LC-MS turned out to be in vain. The different NMN reduced species share the same retention time with the implemented protocol. Nevertheless, with enough amount of substance in hands we managed to use NMR spectroscopy to characterize the obtained products. It is well-known that multidimensional NMR spectroscopy plays an important role in the structure determination in synthetic chemistry, molecular biology and biochemistry and catalysis. By using two-dimensional NMR spectra, not only the funtional groups but also the connectivity among them within a molecule can be characterised. Thus, the structure of peptides and biomacromolecules can be determined.[8–10]

The great challenge we were facing was to deal with a huge signal overlap caused by the mixture of various species. Fortunately, by using TOCSY signals belong to the same molecule (within one spin network) can be identified. The edited $^{1}H$ – $^{13}C$ HSQC spectra extend the scope of information, where characteric $^{13}C$ chemical schifts can be made used for the structure identification. Furthermore, the sign of crosspeaks in the edited HSQC spectra provides unambiguous identification of methylene and methine groups. By careful analysis of the TOCSY and edited HSQC spectra we wre able to determine the main species of those reduction products.

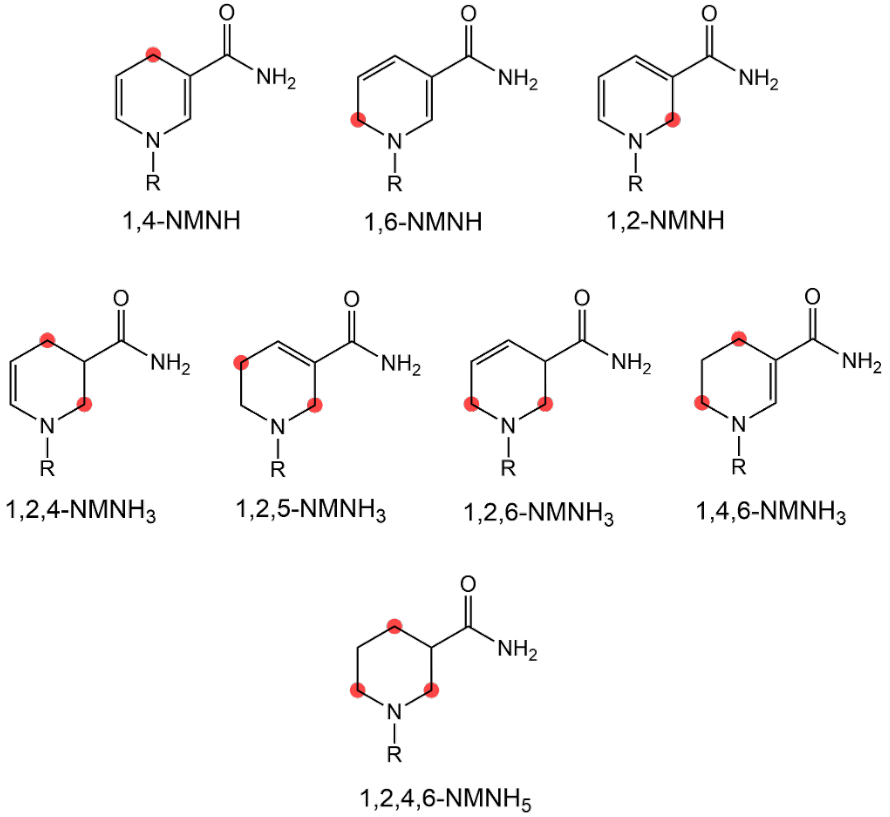

**SI Scheme 9** Tentative reduction products of NMN reduction. Red marks the reduced carbon.

##### The single reduction product 1.1

Starting from the well-resolved signal at 3.057 ppm, TOCSY crosspeaks (**SI Figure 44**) show connectivities with signals at 7.147, 6.214, 5.774, and 5.022 ppm. The edited HSQC spectrum (**SI Figure 45**) shows crosspeaks between 3.057 – 22.0 ppm for a $CH_2$ group, 7.147 – 138.2 ppm, 6.214 – 124.8 ppm and 5.022 – 105.3 ppm for three aromatic CH groups, and 5.774 – 91.7 ppm for a CH group at the anomeric position of a sugar. Those defined fragments correspond well to the three species of single reduction as shown in **SI Scheme 9**. Upon a close inspection into the $^{13}C$ chemical shift at 22.0 ppm, we concluded product **1.1** to be 1,4-NMNH. The $^{13}C$ signal of the $CH_2$ group in both 1,2- and 1,6-NMNH (at the $\alpha$-position of a tertiary amine) would appear at a much lower field (in the range 45 – 50 ppm).[11] Since no such crosspeaks were detected, we excluded the formation of 1,2- and 1,6-NMNH from our products. Two well resolved major peaks at 7.15 ppm (d, 1.5 Hz) and 6.24 ppm (ddt, 8.1, 1.9, 1.7 Hz) were observed. In addition, a pair of very close doublets were observed at 3.056 ppm (1.6 Hz) and 3.062 ppm (1.7 Hz). Those peaks correspond well to $^{1}H$ signals at positions 2, 6 and 4 in the **SI Table 3**, respectively.

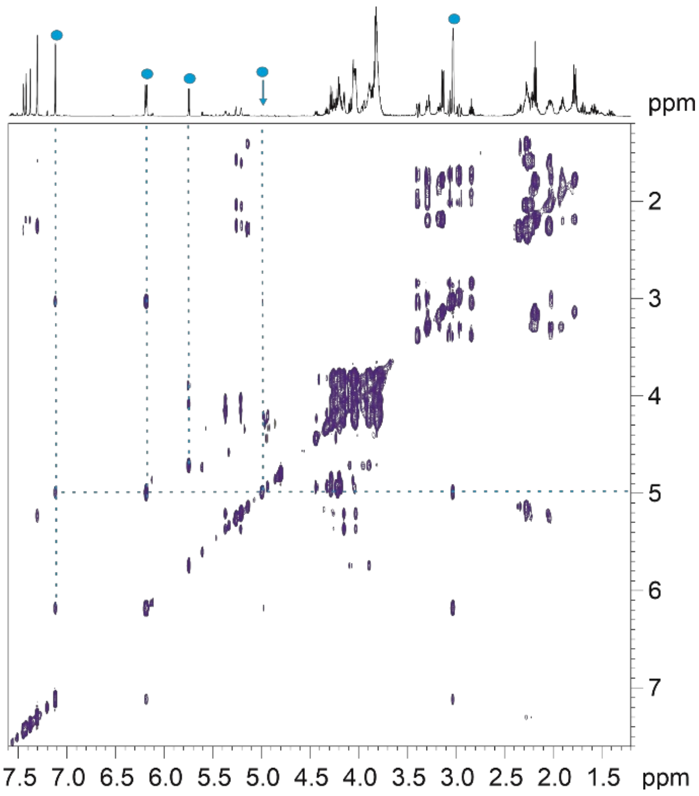

**SI Figure 44** $^{1}H$-$^{1}H$ TOCSY spectrum of NMN products in 0.133 M PBS at 298 K.

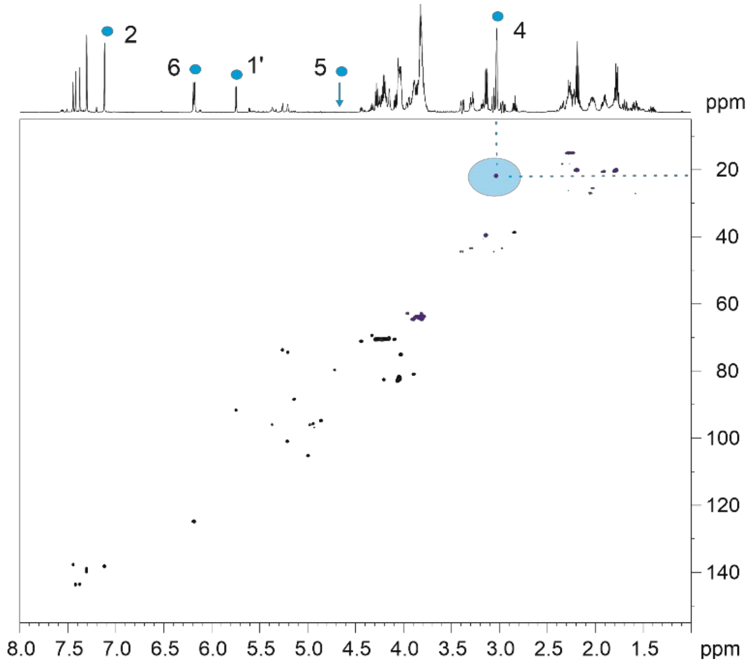

**SI Figure 45**.The edited $^{1}H$-$^{13}C$ HSQC spectrum of products of **1** at 298 K.

#### Products of follow up reactions

As shown in **SI Scheme 9**, two of the tetrahydro-products, 1,2,4- and 1,2,6-$NMNH_3$ contain a CH group within the hetero cyclic ring. This CH group should show a characteristic crosspeak in the edited HSQC spectrum of $^{13}C$ chemical shift in the region 40 – 50 ppm (38), with a sign opposite to a $CH_2$ group. A close inspection of the spectrum shows none of such signals detected. We thus excluded these two species from our products.

Nevertheless, one crosspeak at 2.868 – 38.8 ppm was detected. Close inspection of the crosspeaks in the corresponding region of the TOCSY spectrum (**SI Figure 47**) revealed a connectivity pattern for a meta-substituted piperidine. Therefore, a hexahydro product **1.2** was identified, which is 1,2,4,6-$NMNH_5$. The highlighted peak is for the methine CH and the $^{1}H$ signal at 2.87 ppm stands for proton at carbon 3.

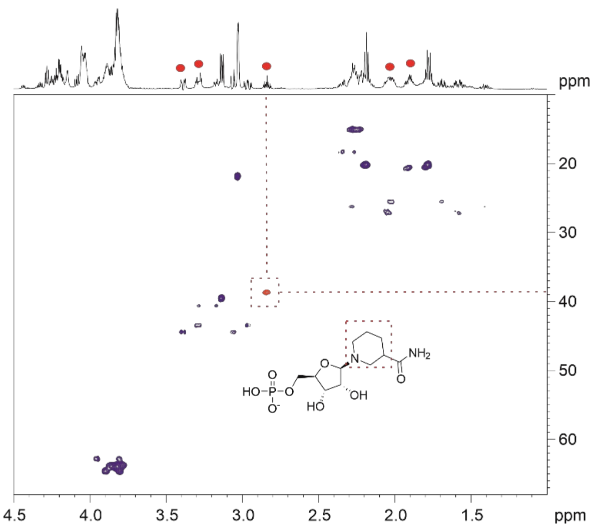

**SI Figure 46**. Section of edited $^{1}H$-$^{13}C$ HSQC spectrum of products of **1** at 298 K.

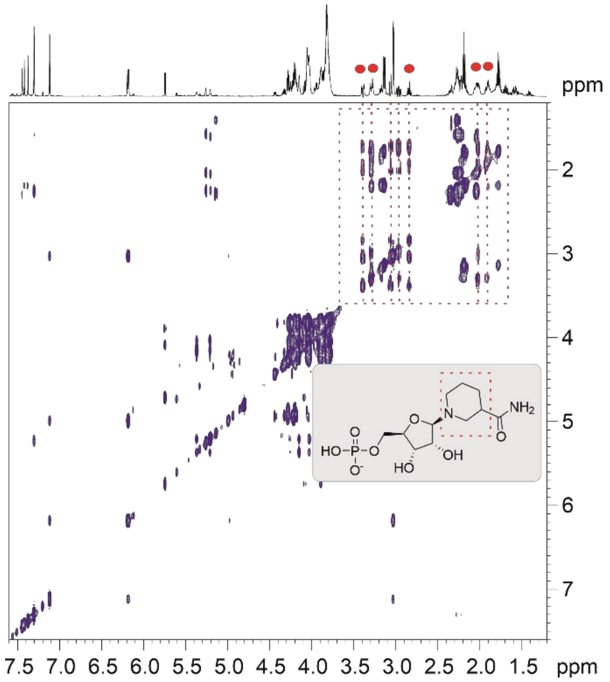

**SI Figure 47** Section of $^{1}H$-$^{1}H$ TOCSY spectrum of products of **1** at 298 K.

Similarly. crosspeaks at 7.448/7.405 – 143.7 (not shown **SI Figure 48**), 3.165 – 39.7, 2.215 – 20.3, and 1.808 – 20.3 ppm were detected in the edited HSQC spectrum (**SI Figure 48**). together with connectivities observed with TOCSY. The tetrahydro-product **1.3** was identified to be 1,4,6-$NMNH_3$.

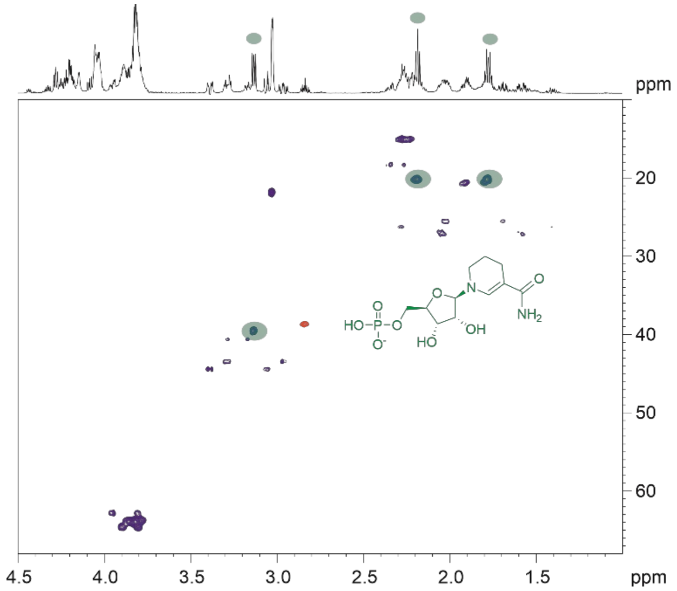

**SI Figure 48** Section of edited $^{1}H$-$^{13}C$ HSQC spectrum of products of **1** at 298 K.

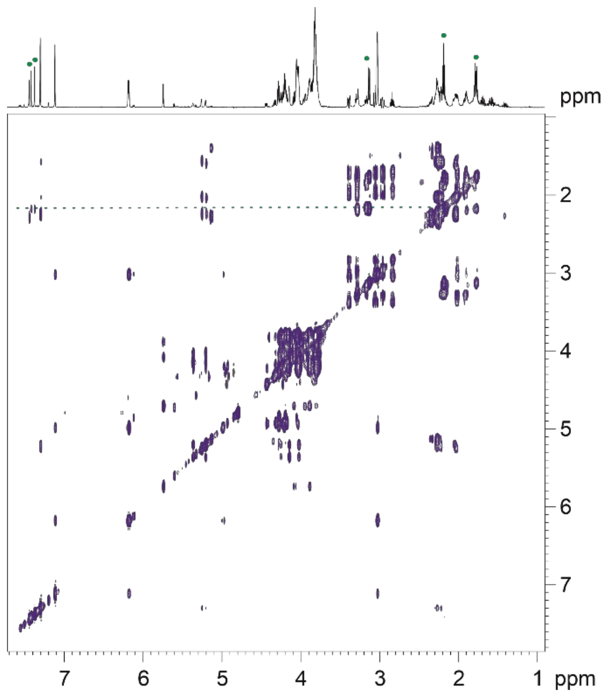

**SI Figure 49** Section of $^{1}H$-$^{1}H$ TOCSY spectrum of products of **1** at 298 K.

In summary, in the reaction mixture of substrate **1.** we were able to identify product **1.1** (1,4-NMNH), product **1.2** (1,2,4,6-$NMNH_5$), and product **1.3** (1,4,6-$NMNH_3$).

#### Identification of hydration product $NMNH_2OH$.

In order to identify the peak at 7.35 ppm in the $^{1}H$ spectrum we put further efforts. Via LC-MS a hydration product with molecular formula $C_{11}H_{18}N_2O_9P^-$ and 353.07323 m/z (mass accuracy of 5 ppm) was revealed (**SI Figure 58**). Following the first reduction step, hydration upon 1,4-NMNH took place and products 1,4-$NMNH_2OH$ formed. Thus, crosspeaks at 7.35 – 138.9. 7.35 – 139.9.

5.29 – 73.9, 5.24 – 74,6, 2.25/2.31 – 15.3 and 2.07/1.06 – 27.1 ppm were detected in the edited HSQC spectrum (**SI Figure 51**). Together with connectivity in TOCSY spectrum (**SI Figure 50**) hydration products as revealed by LC-MS were verified.

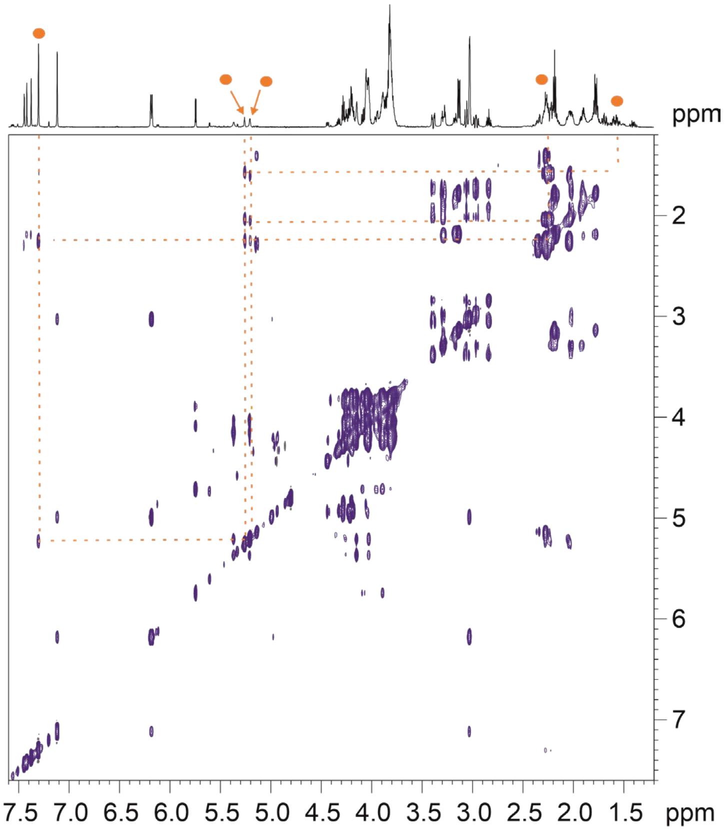

**SI Figure 50** Section of [1]H-[1]H TOCSY spectrum of products of 1 at 298 K. with highlight for the hydration products.

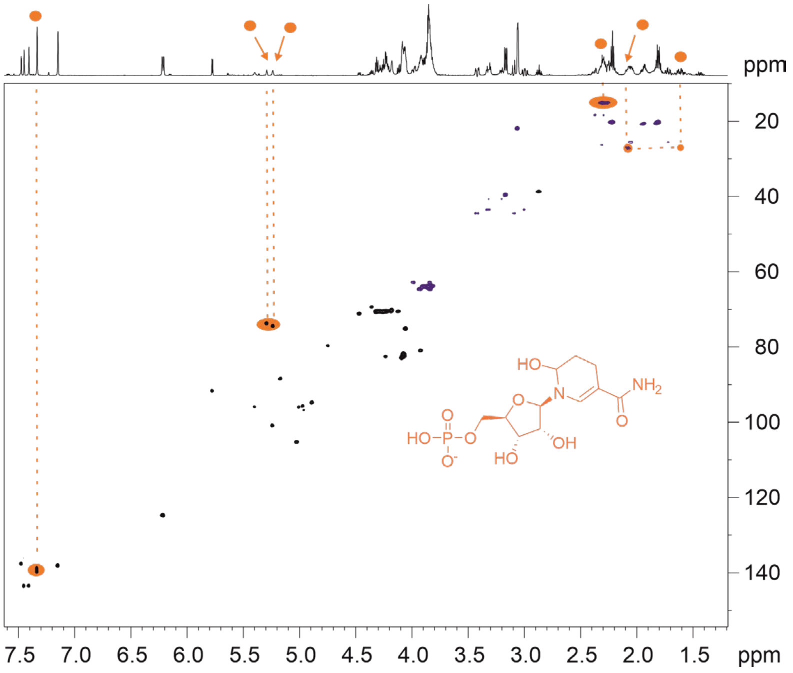

**SI Figure 51** Section of edited $^{1}H$-$^{13}C$ HSQC spectrum of products of 1 at 298 K. with highlight for the hydration products.

In summary. in the reaction mixture of substrate 1 we were able to identify product 1.1 (1,4-NMNH). product 1.2 (1,2,4,6-$NMNH_5$), product 1.3 (1,4,6-$NMNH_3$) and verify the hydration product 1,4 ($NMNH_2OH$).

#### Identification of products of substrate 2

**SI Figure 52** shows the $^{1}H$ spectrum of substrate **2** in 0.133 M PBS at 298 K. Characteristic signals at 9.339, 9.156, 8.825, 8.198 ppm were observed for H-2. H-6. H-4 and H-5 of the nicotinamide unit, signals at 8.379 and 8.019 ppm were detected for H-8 and H-2 of the adenine unit, while two signals at 6.097 and 5.993 ppm for H-1'of nicotinamide and adenine. Respectively, negligible impurities may exist.

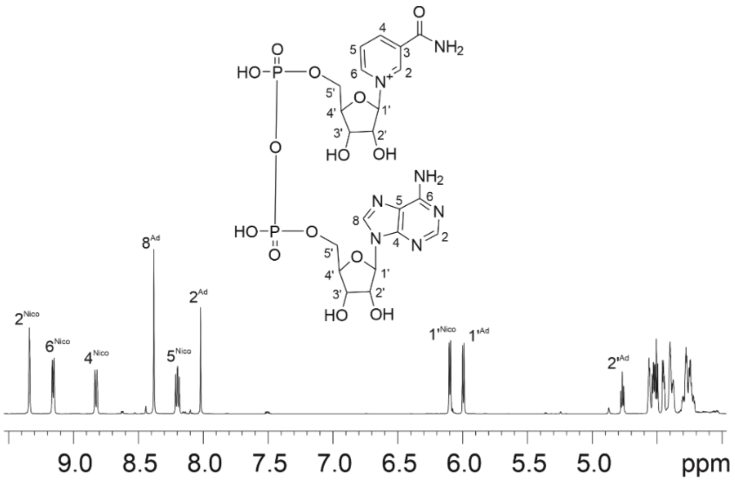

**SI Figure 52** $^{1}H$ spectrum of substrate $NAD^{+}$ (**2**) in 0.133 M PBS at 298 K.

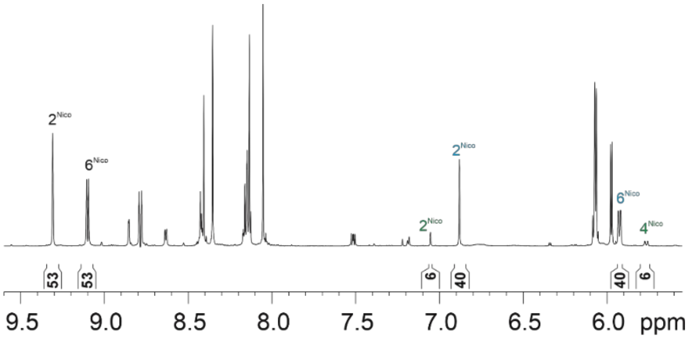

**SI Figure 53** $^1$H spectrum in the region 5.6 – 9.6 ppm of products of substrate **2** at 298 K. The labels show signal assignment (cyan for the major and green for the minor product) and the integral values show the conversion rates.

The $^1$H spectrum in the region 5.6 – 9.6 ppm of the reaction solution of substrate **2** after 4 hours of incubation in 0.133 M PBS is shown in **SI Figure 53**. Different to the situation of substrate **1**. remaining of the substrate **2** was observed. One major and one minor product could be detected. Signal integral revealed a conversion rate of 40% and6 % for the major and the minor product. respectively. Further products of a scale comparable to impurities may form. too.

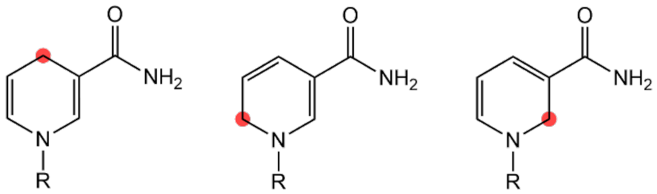

**SI Scheme 10** Tentative reduction products of substrate 2

**SI Scheme 10** shows the possible products of a single reduction of substrate **2**. Section of the TOCSY spectrum is shown in **SI Figure 54**. To start. the double doublet characteristic of a methylene at 2.647 and 2.616 ppm. crosspeaks with signals at 6.903. 5.946. and 4.725 ppm were detected. As shown in **SI Figure 53**. these signals correspond to the main product. In the edited HSQC spectrum. crosspeaks 2.647 – 21.8 and 2.616 – 21.8 ppm were observed. This $^{13}$C chemical shift corresponds to the $C_{N\text{-}4}$ (22.3 ppm) of 1,4-NADH.[7] Similarly. a minor double doublet at 3.913 and 3.776 ppm was observed. whose crosspeaks with signals at 7.075. 5.785 and 5.000 ppm were detected. These signals were assigned to the second product. A close inspection of the edited HSQC spectrum revealed crosspeaks at 3.913 – 41,6 and 3.776 – 41,6 ppm. This $^{13}$C chemical shift corresponds to the $C_{N\text{-}6}$ (42.1 ppm) published for 1,6-NADH.[12] We thus concluded 1,4-NADH and 1,6-NADH to be major and minor products of substrate **2**.

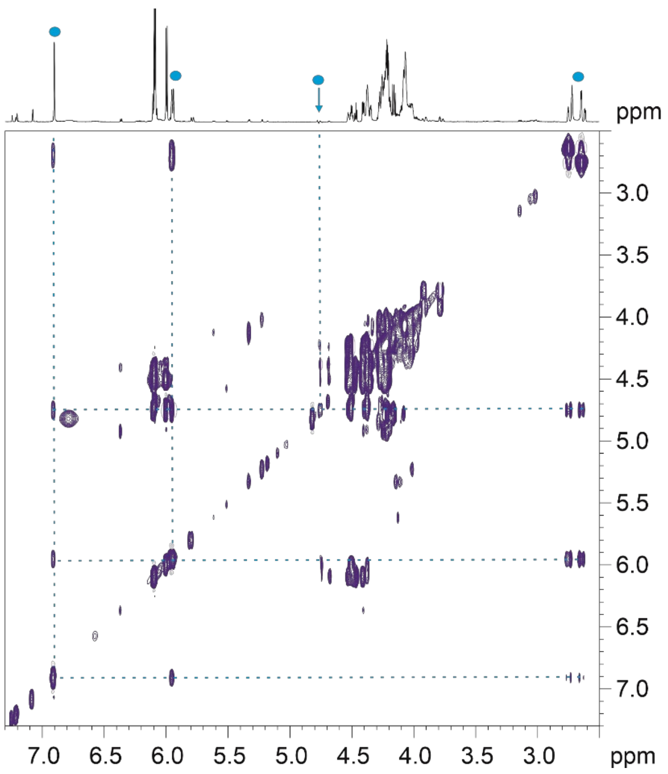

**SI Figure 54** Section of [1]H-[1]H TOCSY spectrum of products of **2** at 298 K. Connectivity among the signals of the major product is highlighted.

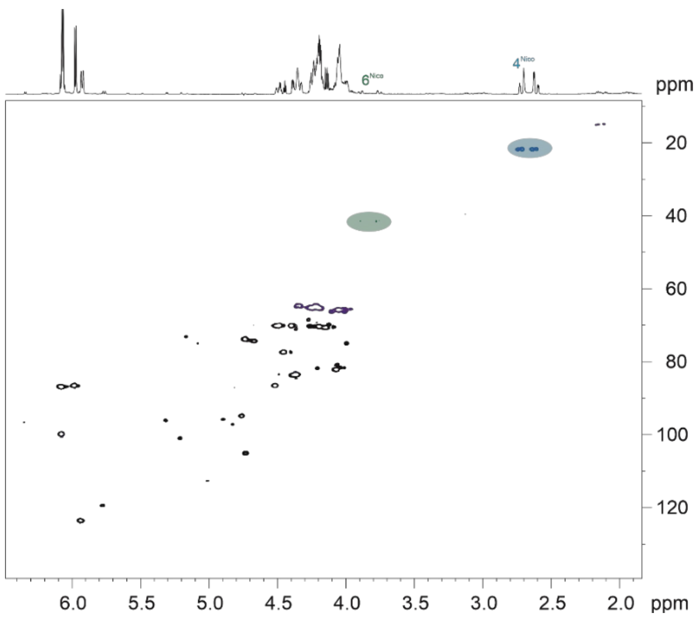

**SI Figure 55** Section of edited $^{1}$H-$^{13}$C HSQC spectrum of products of **2** at 298 K.

### Characterization of the 2nd reaction product with $Fe^0$ and LC-MS (7.35 ppm in $^1$H-NMR)

In order to identify the second product of NMN reduction with $H_2$ and $Fe^0$, sample from such reaction (NMN 18 µmol, nanopowder $Fe^0$ 18 µmol, 4 h. 40 °C. 5 bar $H_2$) was subjected to 2D-NMR and LC-MS qualitative analysis.

LC-MS analysis is in agreement that the main product is 1,4-NMNH. Three different molecules seem to be detected through chromatography, and one matches the retention time of the 1,4-NMN standard. It is possible that we have different conformations of this product in our samples. Its absence in the control confirms it to be a product obtained during the described reaction.

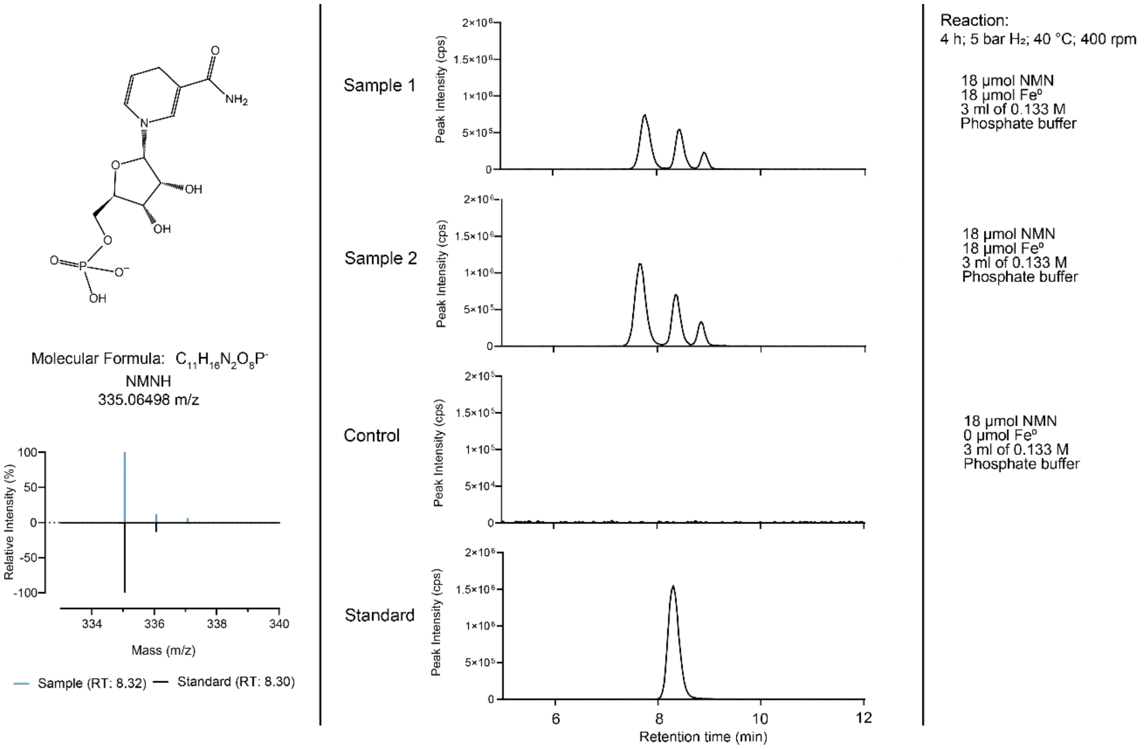

**SI Figure 56** The Extracted Ion Chromatogram (EIC) of mass 335.06368 (mass accuracy of 5 ppm) against the Retention Time (RT) reveals molecules of the mass 335.0649 m/z in the samples from the reaction described on the right. Samples were diluted 1:20 with waster after the reaction for measurement. The natural isotope distribution of the 1,4-NMNH standard (335;06498 m/z) matches that of the samples.

All Diels Alder products suggested in **Scheme 11** share the same mass and charge, which is also the same mass-to-charge of NMNH. Even though molecules of mass 335.06498, were revealed in the chromatogram, analysis of the isotope distribution revealed no doubled charged molecules, thus excluding Diels Alder reactions as products of the reaction with $Fe^0$.

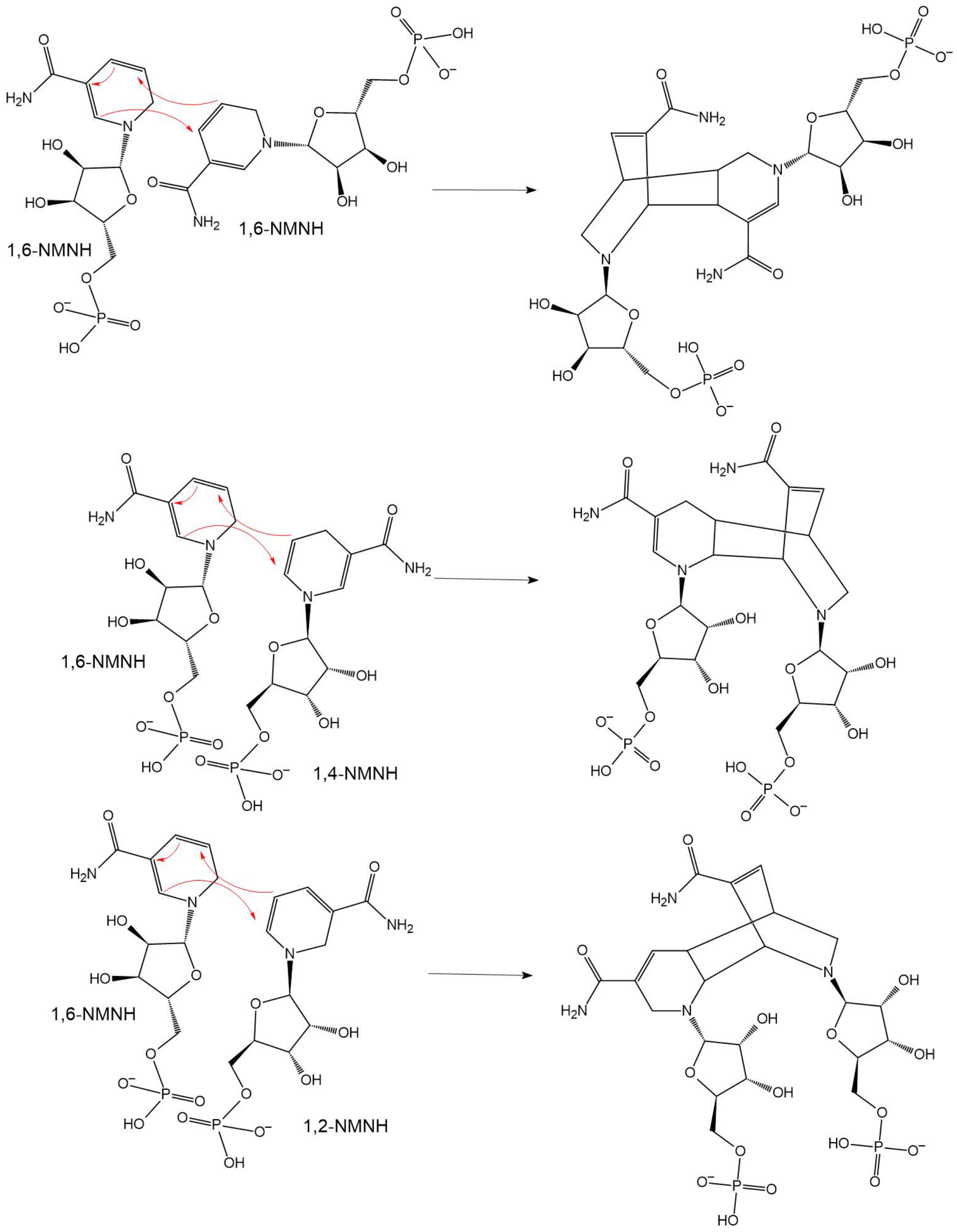

**SI Scheme 11** Possible Diels-Alder reactions to consume 1,4-NMNH

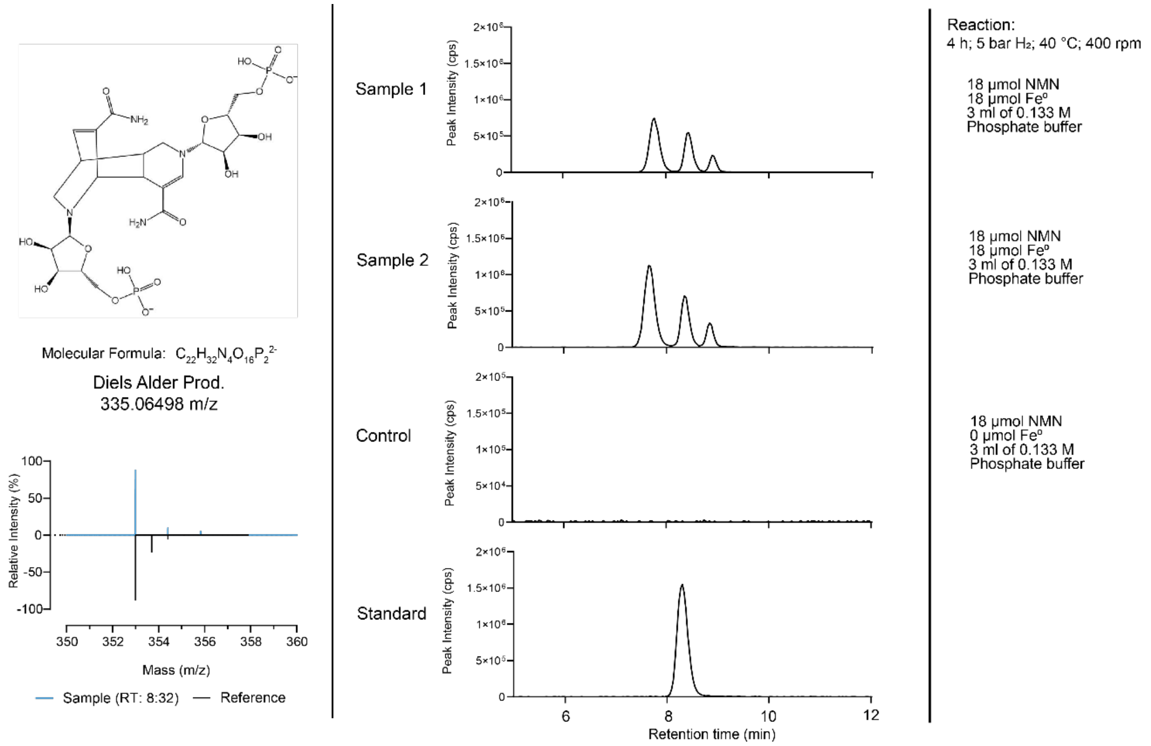

**SI Figure 57** The Extracted Ion Chromatogram (EIC) of mass 335.06368 (mass accuracy of 5 ppm) against the Retention Time (RT) reveals molecules of the mass 335.0649 m/z in the samples from the reaction described on the right. Samples were diluted 1:20 with waster after the reaction for measurement. The standard was prepared in water at similar final concentrations. The theoretical natural isotope distribution of the proposed Diels Alder products (335;06498 m/z), extracted from the website ChemCalc, is compared to that of the samples.

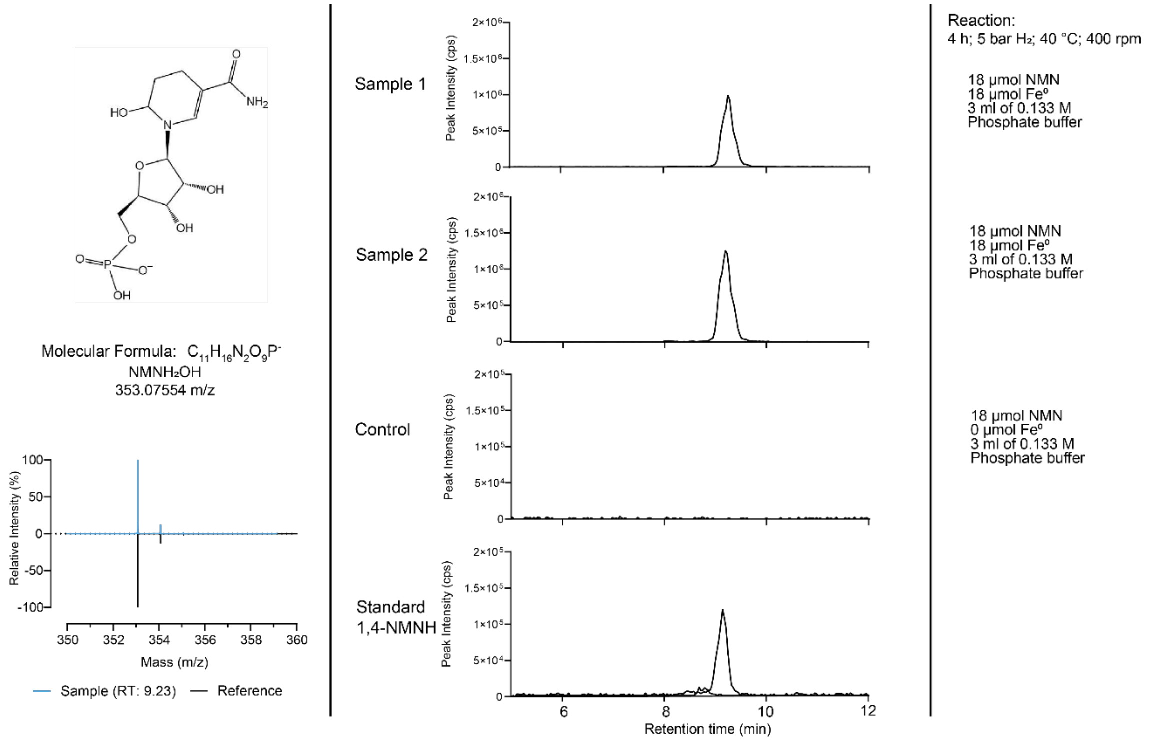

**SI Figure 58** The Extracted Ion Chromatogram (EIC) of mass 353.07323 (mass accuracy of 5 ppm) against the Retention Time (RT) reveals molecules of the mass 353.07323 m/z in the samples from the reaction described on the right. Samples were diluted 1:20 with waster after the reaction for measurement. The standard was prepared in water at similar final concentrations. The theoretical natural isotope distribution of $NMNH_2OH$ (353.07554 m/z), extracted from the website ChemCalc, is compared to that of the samples.

Further analysis of the samples revealed high amounts of a molecule of mass 353.07554 in the samples, which were not in the control. The mass and the isotope distribution of the sample matches that of $NMNH_2OH$. The same signal was detected in the 1,4-NMNH standard, however, the relative abundance to the standard itself, shows it is likely a small contamination from the synthesis of the standard or a consequence of sample preparation and injection into the LC-MS column. The relative abundance of this molecule to 1,4-NMNH, in the samples, is much more significant, being likely a product of the $Fe^0$ reaction. Its abundance suggests it could be the molecule detected at 7.35 ppm ($^1$H-NMR). 2D-NMR analysis confirmed the proposed structure to be of $NMNH_2OH$ (**SI Figure 50** and **SI Figure 51**).

### Scanning Transmission Electron Microscope of Ni-Fe-nanopowders

A

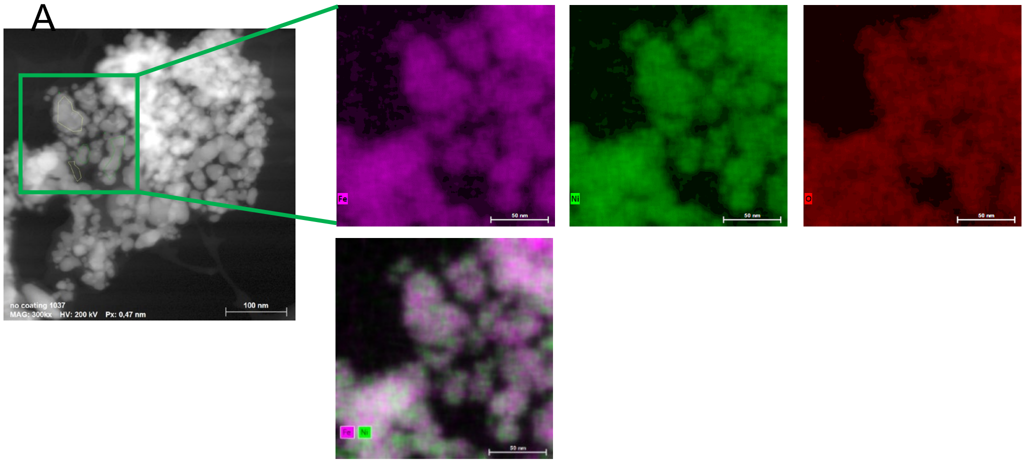

B

| | | | | | | | | | | | | | |
|---|---|---|---|---|---|---|---|---|---|---|---|---|---|
| ✓ | EDS | 1 | | 0,10 | C | O 22,0 | Na | Mg 0,7 | P 3,1 | S 2,1 | Cl 0,7 | Ca 3,3 | Fe 36,1 | Ni 32,0 |
| ✓ | EDS | 2 | | 0,00 | C | O 50,7 | Na 2,4 | Mg 0,9 | P 6,9 | S | Cl 0,5 | Ca 11,3 | Fe 17,6 | Ni 9,7 |
| ✓ | EDS | 3 | | 0,00 | C | O 13,2 | Na 0,5 | Mg 0,7 | P 1,5 | S 0,8 | Cl 0,0 | Ca 2,6 | Fe 46,3 | Ni 34,3 |
| ✓ | EDS | 4 | | 0,02 | C | O 14,0 | Na 1,0 | Mg 1,0 | P 1,2 | S 0,7 | Cl | Ca 3,6 | Fe 45,7 | Ni 32,8 |

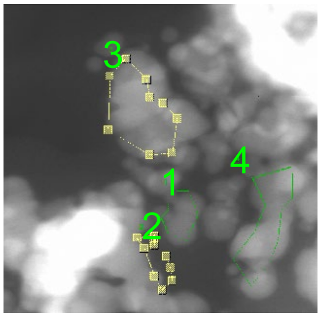

**SI Figure 59** NiFe (1:1) nanopowder pre-reaction STEM observation. (A) STEM-EDS analyses show that NiFe nanopowder shows a mostly even distribution of nickel an iron throughout the nanoparticles. (B) On average, there is slightly more Fe than Ni. A thin oxide layer covers the particles, probably shielding the particles from further oxidation. In comparison with $NiFe_3$ particles (**SI Figure 60**). NiFe shows less Fe-rich regions.

A

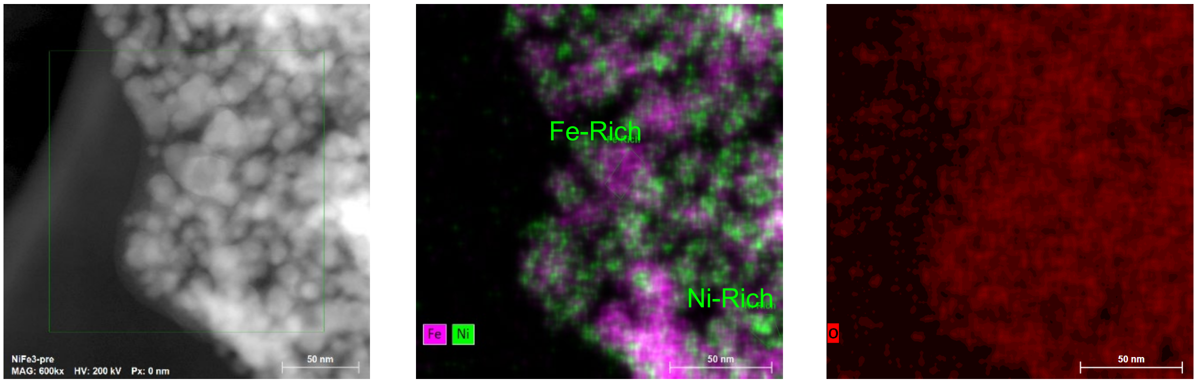

| | | | | | | | | | | | | | | | | |
|---|---|---|---|---|---|---|---|---|---|---|---|---|---|---|---|---|
| EDS | In Total | 0,01 | 100,0 | C | O 6,6 | Na | Mg | Al | Si | P | S | Cl | Ca | Fe 64,6 | Ni 28,8 | Cu |
| EDS | Ni-Rich | 0,00 | 100,0 | C | O 6,8 | Na | Mg | Al | Si | P | S | Cl | Ca | Fe 44,9 | Ni 48,3 | Cu |
| EDS | Fe-Rich | 0,00 | 100,0 | C | O 5,1 | Na | Mg | Al | Si | P | S | Cl | Ca | Fe 83,8 | Ni 11,0 | Cu |

B

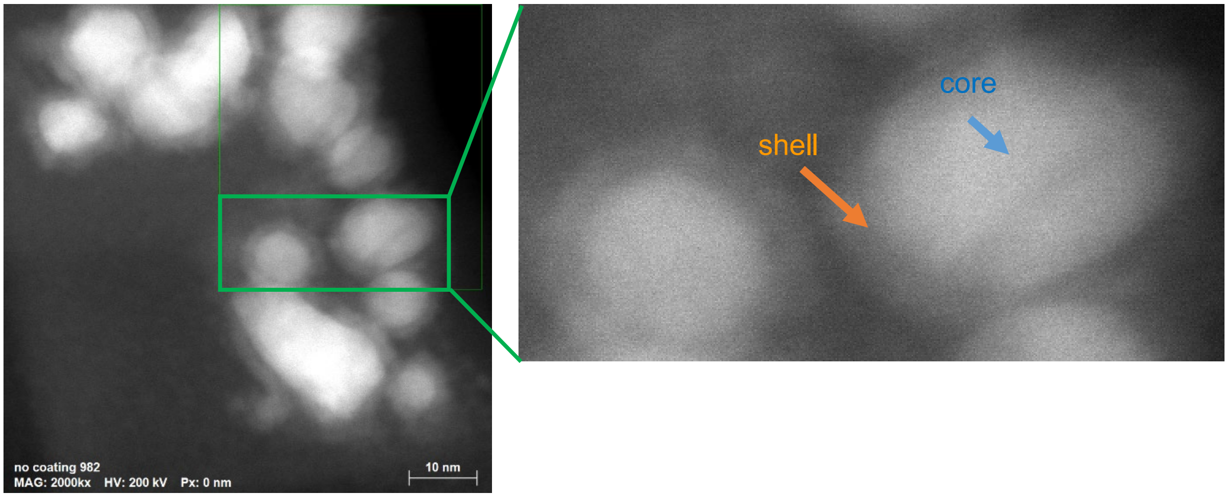

| | | | | | | | | | | | | | |
|---|---|---|---|---|---|---|---|---|---|---|---|---|---|
| EDS | shell | 0,12 | 100,0 | C | O 37,5 | Na 14,6 | Mg 4,8 | P 0,5 | S 1,7 | Cl 4,2 | Ca 8,8 | Fe 16,8 | Ni 11,1 |
| EDS | core | 0,08 | 100,0 | C | O 13,3 | Na 3,4 | Mg | P 1,7 | S 1,7 | Cl 1,5 | Ca 5,2 | Fe 35,3 | Ni 37,8 |

**SI Figure 60** $NiFe_3$ (1:3) nanopowder pre-reaction STEM observation. (A) STEM-EDS analyses show that $NiFe_3$ nanopowder shows the overall distribution of Ni and Fe is 1:3, but we find both metals are not distributed completely evenly, probably leading to the well hydrogenation yields in the reactions. There are iron rich areas that can predominantly be oxidized, delivering nascent $H_2$ to the NiFe regions of the same powder. (B) A thin oxide layer covers the particles, probably shielding them from further oxidation before the reaction.

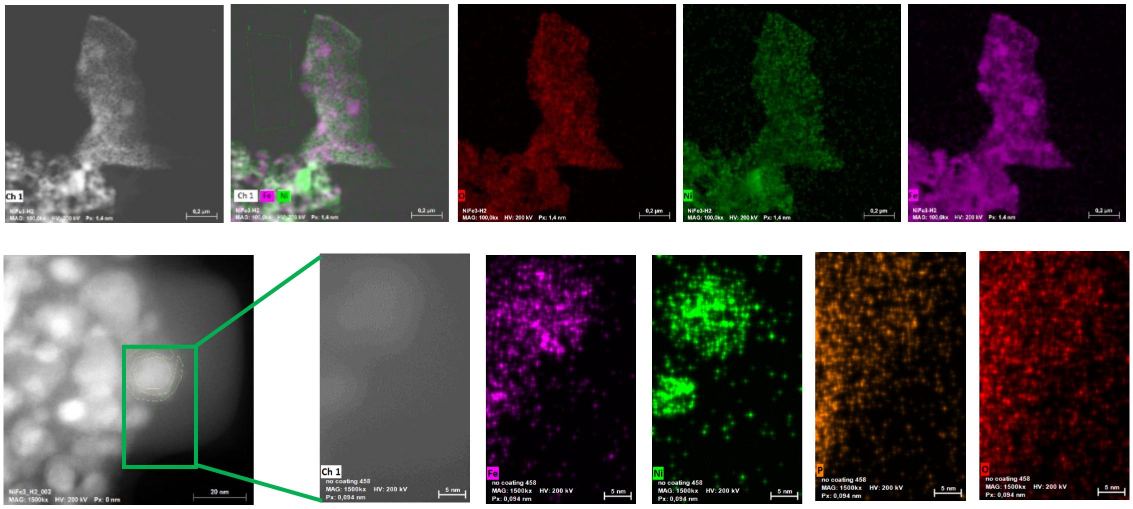

| | | | | | | | | | | | | | | | |
|---|---|---|---|---|---|---|---|---|---|---|---|---|---|---|---|
| | EDS | ROI | 0,01 | C | O 42,9 | Na 1,4 | Mg 1,7 | Al 0,5 | Si 0,6 | P 9,4 | S 0,3 | Cl 1,0 | Ca 4,9 | Fe 24,8 | Ni 12,5 |
| ✓ | EDS | shell | 0,01 | C | O 52,9 | Na 2,1 | Mg 2,0 | Al 1,6 | Si 0,3 | P 10,8 | S 0,0 | Cl 1,0 | Ca 5,8 | Fe 20,5 | Ni 3,1 |
| ✓ | EDS | core | 0,01 | C | O 29,1 | Na 0,3 | Mg 0,4 | Al | Si 0,3 | P 6,7 | S 0,6 | Cl 0,7 | Ca 3,3 | Fe 34,8 | Ni 23,8 |

**SI Figure 61** $NiFe_3$ (1:3) nanopowder after reaction in $H_2$ atmosphere – STEM observation. STEM-EDS analyses show that the Fe of $NiFe_3$ nanopowder gets associated with the phosphate used in the buffer. This is being determined via the ratios between Fe, P and O (1.5:1:4) in the elemental analysis of the shells forming around the nanoparticles. From previous work [7] we know, that Fe is capable of producing nascent hydrogen while being oxidized, these findings are a more direct proof of this happening. The growing phosphate layer could also ultimately lead to the Fe(0) containing minerals to decrease their reaction-promoting ability – we posit that the $H_2$ in the atmosphere can re-cycle the oxidized Fe back to $Fe^0$ and thus keep the reaction going for a longer period of time.

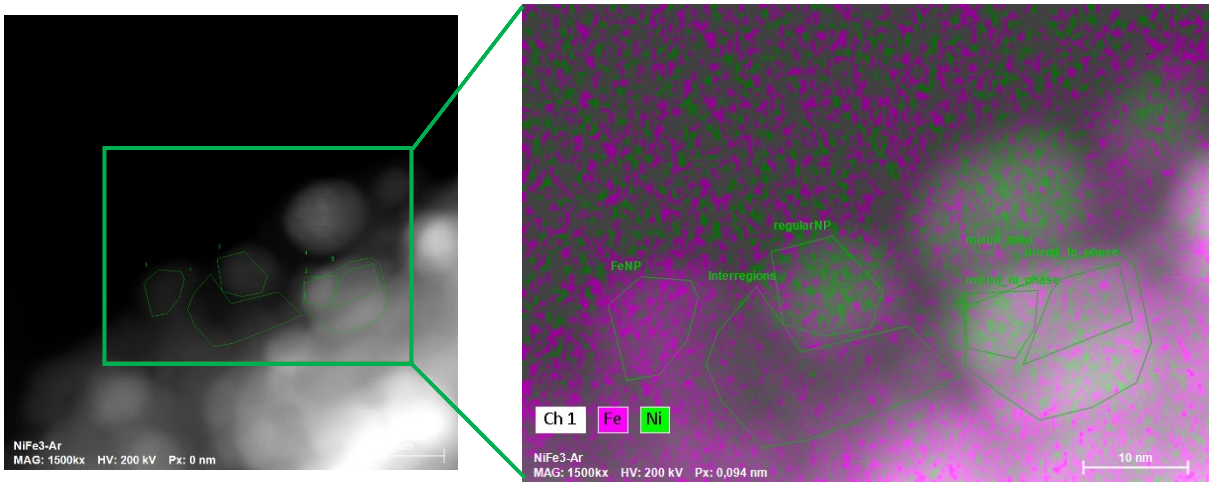

| | | | | | | | | | | | | | | | |
|---|---|---|---|---|---|---|---|---|---|---|---|---|---|---|---|
| ✓ | EDS | Interregions | 0,04 | C | O 49,3 | Na 2,8 | Mg 2,3 | Al 0,9 | Si 0,1 | P 12,8 | S 0,1 | Cl 0,1 | Ca 9,1 | Fe 22,0 | Ni 0,5 |
| ✓ | EDS | FeNP | 0,04 | C | O 48,6 | Na 1,2 | Mg 1,4 | Al 0,5 | Si 0,7 | P 2,0 | S | Cl 0,1 | Ca 1,1 | Fe 44,4 | Ni 0,0 |
| ✓ | EDS | regularNP | 0,06 | C | O 16,2 | Na 1,1 | Mg 1,1 | Al 0,4 | Si 0,2 | P 4,8 | S 0,7 | Cl | Ca 3,3 | Fe 42,4 | Ni 29,9 |
| ✓ | EDS | mixed_ni_phase | 0,18 | C | O 29,4 | Na 2,2 | Mg 1,8 | Al 0,4 | Si 0,5 | P 7,6 | S 0,2 | Cl | Ca 5,4 | Fe 32,3 | Ni 20,1 |
| ✓ | EDS | mixed_total | 0,16 | C | O 34,0 | Na 1,9 | Mg 1,4 | Al 1,1 | Si 0,3 | P 7,2 | S 0,4 | Cl | Ca 5,2 | Fe 33,6 | Ni 14,8 |
| ✓ | EDS | mixed_fe_phase | 0,18 | C | O 37,8 | Na 2,0 | Mg 1,4 | Al 0,7 | Si | P 8,8 | S 0,3 | Cl | Ca 6,6 | Fe 30,6 | Ni 11,9 |

**SI Figure 62** $NiFe_3$ (1:3) nanopowder after reaction in Ar atmosphere – STEM observation. STEM-EDS analyses indicate that the Fe of $NiFe_3$ nanopowder likely gets oxidized, forming $Fe_3(PO_4)_2$ with the phosphate buffer. The cleaning process of the nanoparticles after the reaction (s. Methods) strengthens the assumption that Fe and phosphate are more than just loosely associated. This is being determined via the ratios between Fe. P and O. This mapping shows that the regions between the nanoparticles almost consist exclusively of iron-phosphates precipitating from the reactions.

### Abiotic oxidation of organic cofactors and reduction of pyruvate

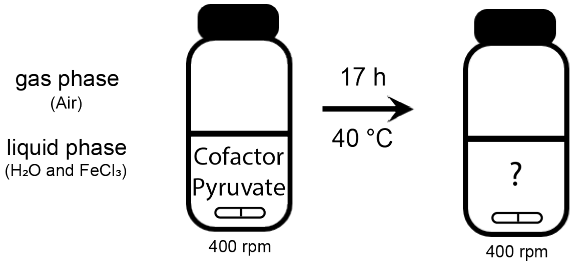

**SI Scheme 12** An aqueous mixture of 0.1 mL with 100 mM pyruvate, twice as much 1,4-NADH or 1,4-NMNH, and 60 mM $FeCl_3$ reacted overnight at 40 °C and 400 rpm to produce lactate, according to the scheme. The pH before and after the reaction is <5.

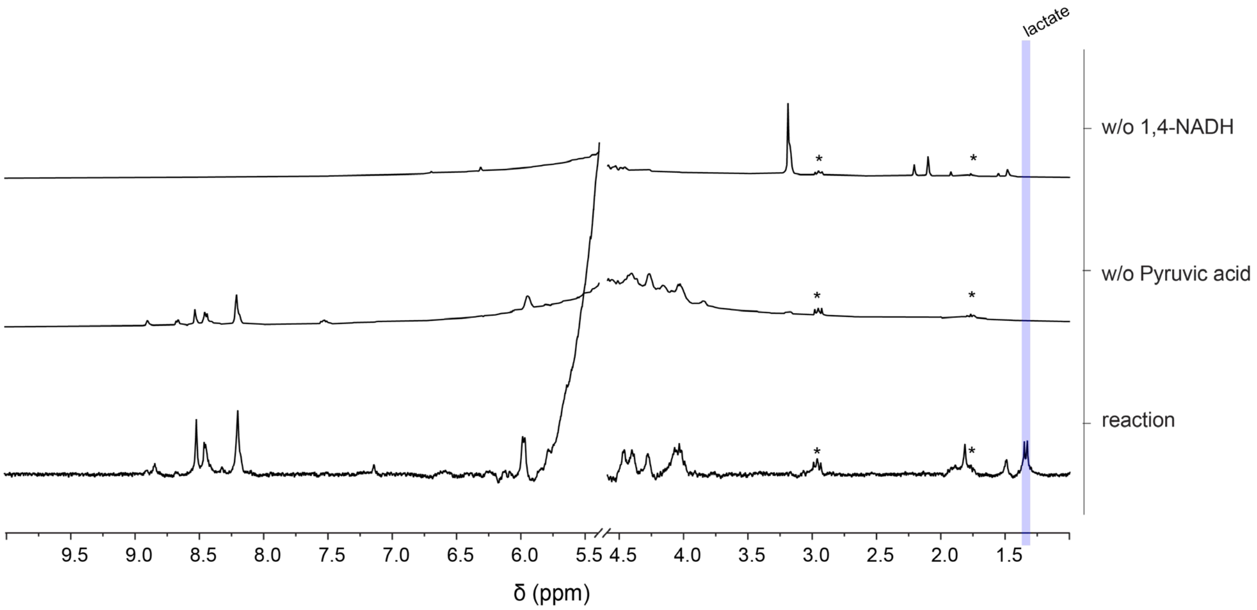

**SI Figure 63** A NMR spectrum of a replica sample of aqueous 1,4-NADH (20 mmol) were mixed in an Eppendorf tube with 10 mmol of pyruvic acid, and 60 mmol of $FeCl_3$, as indicated by **SI Scheme 12**, is stacked together in this figure with the respective controls. The latter were without NADH and without pyruvic acid as described on the right side of each spectra. The metals were precipitated with 0.2 mL of a thiolate/phosphate solution. The supernatant was collected and DSS added as an internal standard. The spectra were edited to only include relevant peaks, having been removed a DSS peak at 0 ppm and water peak at 4.8 ppm. No other peaks were found in the areas removed. Some DSS peaks are still visible (*). The peaks used for qualitative analysis and subsequent qNMR are highlighted in blue (1.33 ppm).

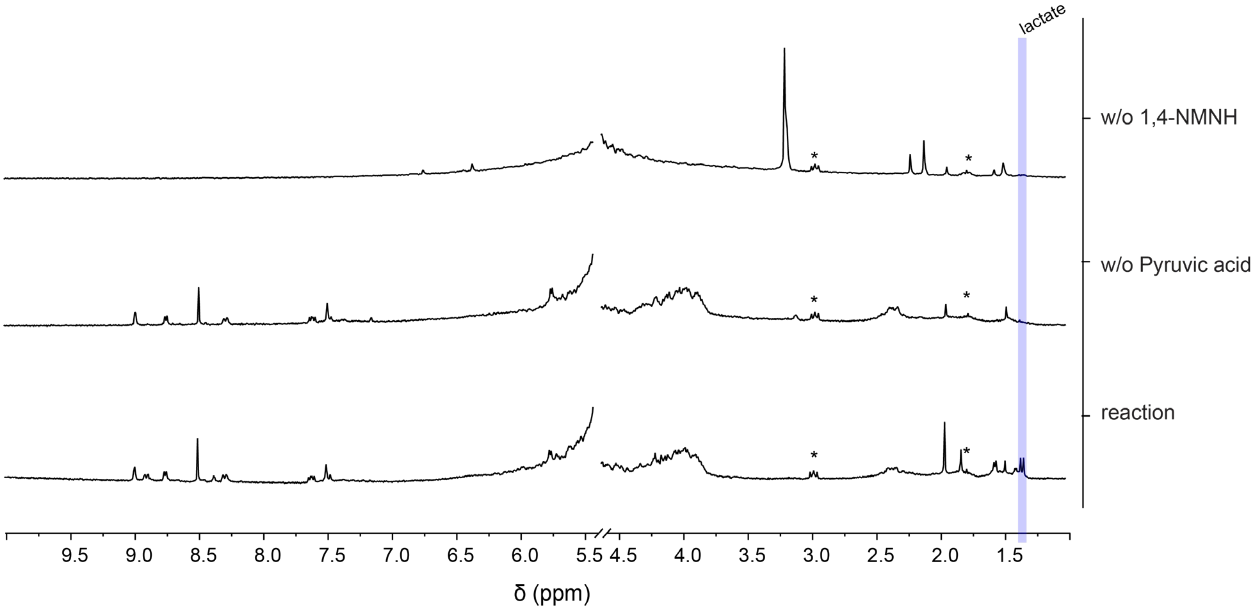

**SI Figure 64** A NMR spectrum of a replica sample of aqueous NMNH (20 mmol) were mixed in an Eppendorf tube with 10 mmol of pyruvic acid, and 6 mmol of $FeCl_3$, as indicated by **SI Scheme 10**, is stacked together in this figure with the respective controls. The latter were without NADH and without pyruvic acid as described on the right side of each spectra. The metals were precipitated with 0.2 mL of a thiolate/phosphate solution. The supernatant was collected and DSS added as an internal standard. The spectra were edited to only include relevant peaks, having been removed a DSS peak at 0 ppm and water peak at 4.8 ppm. No other peaks were found in the areas removed. Some DSS peaks are still visible (*). The peaks used for qNMR are highlighted in blue (1.33 ppm).

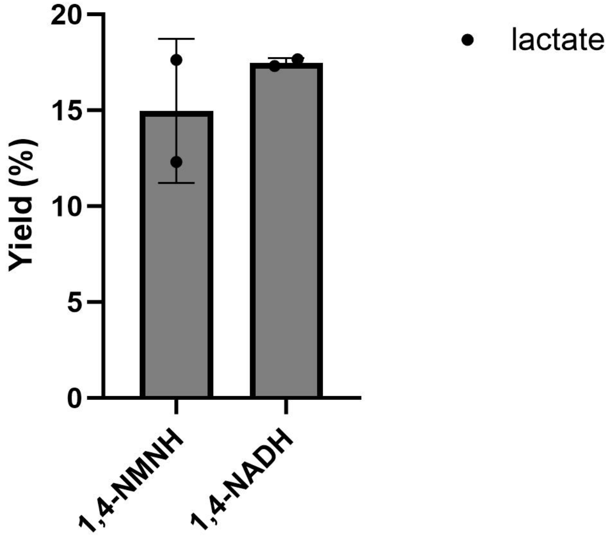

**SI Figure 65** From duplicates of the reaction setup described in **SI Scheme 10**, similar amounts of lactate were obtained starting from 1,4-NADH and 1,4-NMNH as hydride donors. The yields were obtained by integration of the lactate duplet at approximately

### Proposed surface interaction between NAD/NMN and Ni/Fe minerals

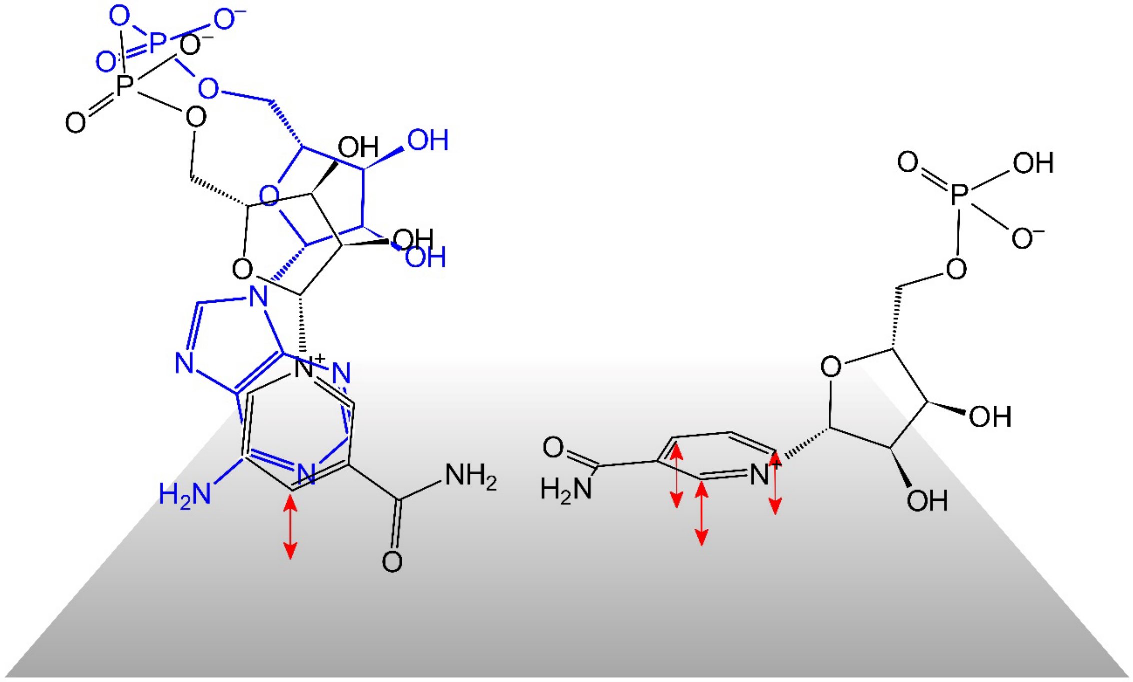

**SI Scheme 13** Proposed surface interaction with $NAD^+$ and NMN. Due to its dinucleotide structure. NAD alternates between the folded and open conformation. This could prevent surface hydrides from reaching all carbons of the nicotinamide ring. NMN does not have such conformation and would be able to interact more directly with the surface.
